# Hybridization chain reaction enables a unified approach to multiplexed, quantitative, high-resolution immunohistochemistry and in situ hybridization

**DOI:** 10.1101/2021.06.02.446311

**Authors:** Maayan Schwarzkopf, Mike C. Liu, Samuel J. Schulte, Rachel Ives, N. Husain, Harry M.T. Choi, Niles A. Pierce

## Abstract

RNA in situ hybridization (RNA-ISH) based on the mechanism of hybridization chain reaction (HCR) enables multiplexed, quantitative, high-resolution RNA imaging in highly autofluorescent samples including whole-mount vertebrate embryos, thick brain slices, and formalin-fixed paraffin-embedded (FFPE) tissue sections. Here, we extend the benefits of 1-step, multiplexed, quantitative, isothermal, enzyme-free HCR signal amplification to immunohistochemistry (IHC), enabling accurate and precise protein relative quantitation with subcellular resolution in an anatomical context. Moreover, we provide a unified framework for simultaneous quantitative protein and RNA imaging with 1-step HCR signal amplification performed for all target proteins and RNAs simultaneously.

**SUMMARY:** Signal amplification based on the mechanism of hybridization chain reaction enables multiplexed, quantitative, high-resolution imaging of protein and RNA targets in highly autofluorescent tissues.

## INTRODUCTION

Biological circuits encoded in the genome of each organism direct development, maintain integrity in the face of attacks, control responses to environmental stimuli, and sometimes malfunction to cause disease. RNA in situ hybridization (RNA-ISH) methods (Gall & Pardue, 1969; Cox *et al*., 1984; Tautz & Pfeifle, 1989; Rosen & Beddington, 1993; Wallner *et al*., 1993; Nieto *et al*., 1996; Thisse & Thisse, 2008) and immunohistochemistry (IHC) methods (Coons *et al*., 1941; Takakura *et al*., 1997; Sillitoe & Hawkes, 2002; Ahnfelt-Ronne *et al*., 2007; Psychoyos & Finnell, 2009; Ramos-Vara & Miller, 2014; Fujisawa *et al*., 2015; Staudt *et al*., 2015) provide biologists, drug developers, and pathologists with critical windows into the spatial organization of this circuitry, enabling imaging of RNA and protein expression in an anatomical context. While it is desirable to perform multiplexed experiments in which a panel of targets are imaged quantitatively at high resolution in a single specimen, using traditional RNA-ISH and IHC methods in highly autofluorescent samples including whole-mount vertebrate embryos and FFPE tissue sections, multiplexing is cumbersome, staining is non-quantitative, and spatial resolution is routinely compromised by diffusion of reporter molecules. These multi-decade technological shortcomings are significant impediments to biological research as well as to advancement of drug development and pathology assays, impeding high-dimensional, quantitative, high-resolution analyses of developmental and disease-related regulatory networks in an anatomical context.

RNA-ISH methods detect RNA targets using nucleic acid probes (Qian *et al*., 2004; Silverman & Kool, 2007) and IHC methods detect protein targets using antibody probes (Ramos-Vara & Miller, 2014). In either case, probes can be direct-labeled with reporter molecules (Kislauskis *et al*., 1993; Femino *et al*., 1998; Levsky *et al*., 2002; Kosman *et al*., 2004; Capodieci *et al*., 2005; Chan *et al*., 2005; Raj *et al*., 2008), but to increase the signal-to-background ratio, are more often used to mediate signal amplification in the vicinity of the probe (Qian *et al*., 2004; Silverman & Kool, 2007; Ramos-Vara & Miller, 2014). A variety of in situ amplification approaches have been developed including immunological methods (Macechko *et al*., 1997; Hughes & Krause, 1998; Kosman *et al*., 2004), branched DNA methods (Collins *et al*., 1997; Bushnell *et al*., 1999; Player *et al*., 2001; Qian & Lloyd, 2003; Wang *et al*., 2012; Kishi *et al*., 2019; Saka *et al*., 2019), in situ PCR methods (Wiedorn *et al*., 1999; Qian & Lloyd, 2003), and rolling circle amplification methods (Zhou *et al*., 2001; Schweitzer & Kingsmore, 2001; Larsson *et al*., 2004; Zhou *et al*., 2004; Larsson *et al*., 2010). However, for both RNAISH (Tautz & Pfeifle, 1989; Harland, 1991; Lehmann & Tautz, 1994; Kerstens *et al*., 1995; Nieto *et al*., 1996; Pernthaler *et al*., 2002; Kosman *et al*., 2004; Thisse *et al*., 2004; Denkers *et al*., 2004; Clay & Ramakrishnan, 2005; Barroso-Chinea *et al*., 2007; Acloque *et al*., 2008; Piette *et al*., 2008; Thisse & Thisse, 2008; Weiszmann *et al*., 2009; Wang *et al*., 2012) and IHC (Takakura *et al*., 1997; Sillitoe & Hawkes, 2002; Ahnfelt-Ronne *et al*., 2007; Psychoyos & Finnell, 2009; Ramos-Vara & Miller, 2014; Fujisawa *et al*., 2015; Staudt *et al*., 2015), traditional in situ amplification based on enzyme-mediated catalytic reporter deposition (CARD) remains the dominant approach for achieving high signal-to-background in highly autofluorescent samples including whole-mount vertebrate embryos and FFPE tissue sections. CARD is widely used despite three significant drawbacks: multiplexing is cumbersome due to the lack of orthogonal deposition chemistries, necessitating serial amplification for one target after another (Lehmann & Tautz, 1994; Nieto *et al*., 1996; Thisse *et al*., 2004; Denkers *et al*., 2004; Kosman *et al*., 2004; Clay & Ramakrishnan, 2005; Barroso-Chinea *et al*., 2007; Tóth & Mezey, 2007; Acloque *et al*., 2008; Piette *et al*., 2008; Glass *et al*., 2009; Stack *et al*., 2014; Mitchell *et al*., 2014; Tsujikawa *et al*., 2017), staining is qualitative rather than quantitative, and spatial resolution is routinely compromised by diffusion of reporter molecules prior to deposition (Tautz & Pfeifle, 1989; Takakura *et al*., 1997; Sillitoe & Hawkes, 2002; Thisse *et al*., 2004; Thisse & Thisse, 2008; Acloque *et al*., 2008; Piette *et al*., 2008; Weiszmann *et al*., 2009; Psychoyos & Finnell, 2009).

In the context of RNA-ISH, in situ amplification based on the mechanism of hybridization chain reaction (HCR; Figure 1A) (Dirks & Pierce, 2004) overcomes the longstanding shortcomings of CARD to enable multiplexed, quantitative, high-resolution imaging of RNA expression in diverse organisms and sample types including highly autofluorescent samples (Choi *et al*., 2010; Choi *et al*., 2014; Choi *et al*., 2016; Shah *et al*., 2016; Trivedi *et al*., 2018; Choi *et al*., 2018) (e.g., see Table S1). To image RNA expression, targets are detected by nucleic acid probes that trigger isothermal enzyme-free chain reactions in which fluorophore-labeled HCR hairpins self-assemble into tethered fluorescent amplification polymers (Figure 1B). Orthogonal HCR amplifiers operate independently within the sample so the experimental timeline for multiplexed experiments is independent of the number of target RNAs (Choi *et al*., 2010; Choi *et al*., 2014). The amplified HCR signal scales approximately linearly with the number of target molecules (Figure 1E), enabling accurate and precise RNA relative quantitation with subcellular resolution in the anatomical context of whole-mount vertebrate embryos (Trivedi *et al*., 2018; Choi *et al*., 2018). Amplification polymers remain tethered to their initiating probes, enabling imaging of RNA expression with subcellular or single-molecule resolution as desired (Choi *et al*., 2014; Shah *et al*., 2016; Choi *et al*., 2016; Choi *et al*., 2018).

**Fig. 1:**
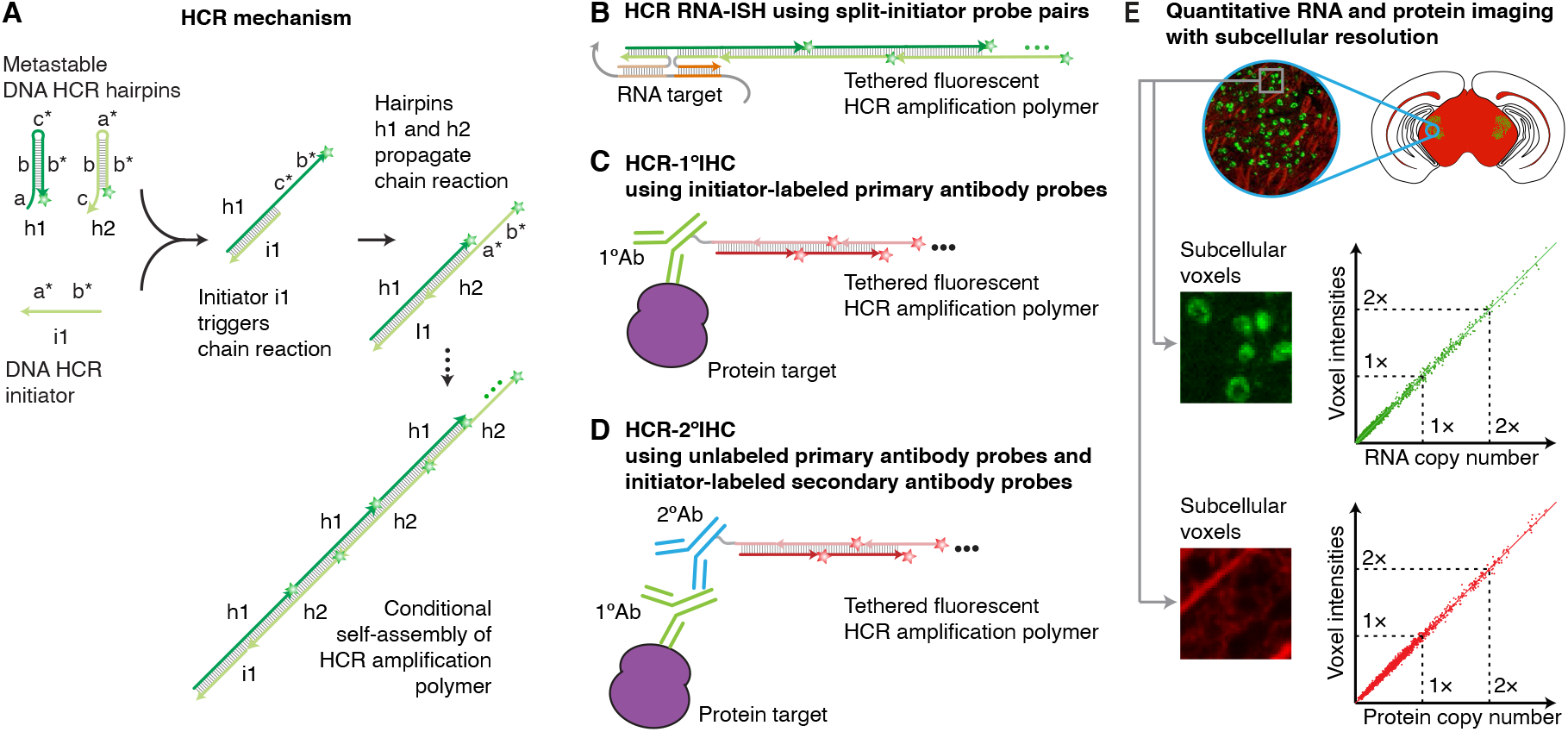
A unified framework for multiplexed, quantitative, high-resolution protein and RNA imaging using HCR 1° IHC + HCR RNA-ISH or HCR 2° IHC + RNA-ISH. (A) 1-step, isothermal, enzyme-free signal amplification via hybridization chain reaction (HCR) (Dirks & Pierce, 2004). Kinetically trapped hairpins h1 and h2 co-exist metastably in solution on lab time scales, storing the energy to drive a conditional self-assembly cascade upon exposure to a cognate initiator sequence i1. Stars denote fluorophores. (B) HCR RNA-ISH using split-initiator probe pairs that hybridize to adjacent binding sites on the target RNA to colocalize a full HCR initiator and trigger HCR. (C) HCR 1° IHC using initiator-labeled primary antibody probes. (D) HCR 2° IHC using unlabeled primary antibody probes and initiator-labeled secondary antibody probes. (E) Conceptual schematic: HCR signal scales approximately linearly with the abundance of a target RNA (green channel) or protein (red channel), enabling accurate and precise relative quantitation with subcellular resolution in an anatomical context.

These properties that make HCR signal amplification well-suited for RNA-ISH appear equally favorable in the context of IHC, suggesting the approach of combining HCR signal amplification with antibody probes (Koos *et al*., 2015; Husain, 2016; Lin *et al*., 2018). Here, we extend the benefits of 1-step, quantitative, enzyme-free signal amplification from RNA-ISH to IHC, validating multiplexed, quantitative, high-resolution imaging of protein expression with high signal-to-background in highly autofluorescent samples, thus overcoming the long-standing shortcomings of IHC using CARD. Moreover, we establish a unified framework for simultaneous multiplexed, quantitative, high-resolution IHC and RNA-ISH, with 1-step HCR signal amplification performed for all targets simultaneously.

## RESULTS

For protein imaging with HCR, we pursue two complementary approaches. Using HCR 1°IHC, protein targets are detected using primary antibody probes labeled with one or more HCR initiators (Figure 1C). For multiplexed experiments, the probes for different targets are labeled with different HCR initiators that trigger orthogonal HCR amplifiers labeled with spectrally distinct fluorophores. Researchers have the flexibility to detect different targets using primary antibody probes raised in the same host species (or a variety of host species as convenient). On the other hand, antibody-initiator conjugation must be validated for each new primary antibody probe. Using HCR 2°IHC, protein targets are detected using unlabeled primary antibody probes that are in turn detected by secondary antibody probes labeled with one or more HCR initiators (Figure 1D). This approach has the advantage that validation of a small library of initiator-labeled secondary antibodies (e.g., 5 secondaries targeting different host species) enables immediate use of large libraries of primary antibody probes (e.g., 10^5^ commercially available primaries) without modification. On the other hand, for multiplexed experiments, each target must be detected using a primary antibody raised in a different host species to enable subsequent detection by an anti-host secondary antibody probe that triggers an orthogonal spectrally distinct HCR amplifier. Hence, depending on the available antibody probes, one may prefer HCR 1°IHC in one instance and HCR 2°IHC in another.

### Multiplexed protein imaging using HCR 1°IHC or HCR 2°IHC

Figure 2 demonstrates multiplexed protein imaging via HCR 1°IHC using initiator-labeled primary antibody probes. Figure 3 demonstrates multiplexed protein imaging via HCR 2°IHC using unlabeled primary antibody probes and initiator-labeled secondary antibody probes. Both methods achieve high signal-to-background for 3-plex protein imaging in mammalian cells and for 4-plex protein imaging in FFPE mouse brain sections. Across 21 protein imaging scenarios (6 in mammalian cells + 10 in FFPE mouse brain sections + 4 in FFPE human breast tissue sections + 1 in whole-mount zebrafish embryos; 9 using 1°IHC HCR + 13 using 2°IHC HCR; 11 using confocal microscopy + 10 using epifluorescence microscopy), the estimated signal-to-background ratio for protein targets ranged from 20 to 610 with a median of 87 (see Tables S9 and S10 for additional details). This level of performance was achieved for all targets simultaneously in 4-channel and 5-channel images (including a DAPI channel in each case) using fluorophores that compete with lower autofluorescence (Alexa647) as well as with higher autofluorescence (Alexa488) and in samples with lower autofluorescence (mammalian cells) and higher autofluorescence (FFPE mouse brain sections).

**Fig. 2:**
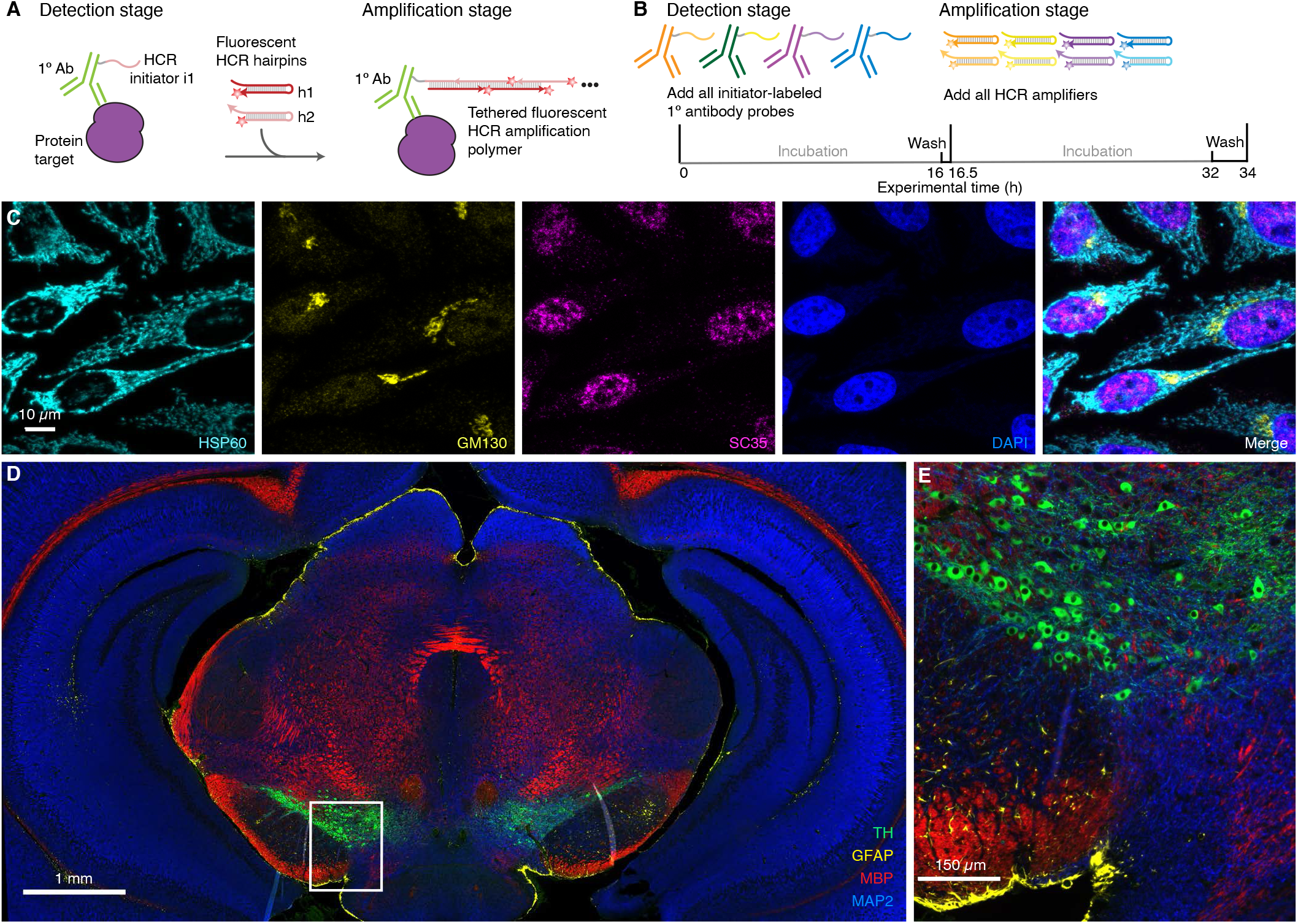
Multiplexed protein imaging via HCR 1° IHC using initiator-labeled primary antibody probes and simultaneous HCR signal amplification for all targets. (A) 2-stage IHC protocol. Detection stage: initiator-labeled primary antibody probes bind to protein targets; wash. Amplification stage: initiators trigger self-assembly of fluorophore-labeled HCR hairpins into tethered fluorescent amplification polymers; wash. (B) Multiplexing timeline. The same 2-stage protocol is used independent of the number of target proteins. (C) Confocal image of 3-plex protein imaging in mammalian cells on a slide; 0.2 × 0.2 *µ*m pixels; maximum intensity z-projection. Target proteins: HSP60 (Alexa488), GM130 (Alexa647), SC35 (Alexa546). Sample: HeLa cells. (D) Epifluorescence image of 4-plex protein imaging in FFPE mouse brain sections; 0.3 × 0.3 *µ*m pixels. Target proteins: TH (Alexa488), GFAP (Alexa546), MBP (Alexa647), MAP2 (Alexa750). (E) Zoom of depicted region of panel D. Sample: FFPE C57BL/6 mouse brain section (coronal); thickness: 5 *µ*m. See Section S5.2 for additional data.

**Fig. 3:**
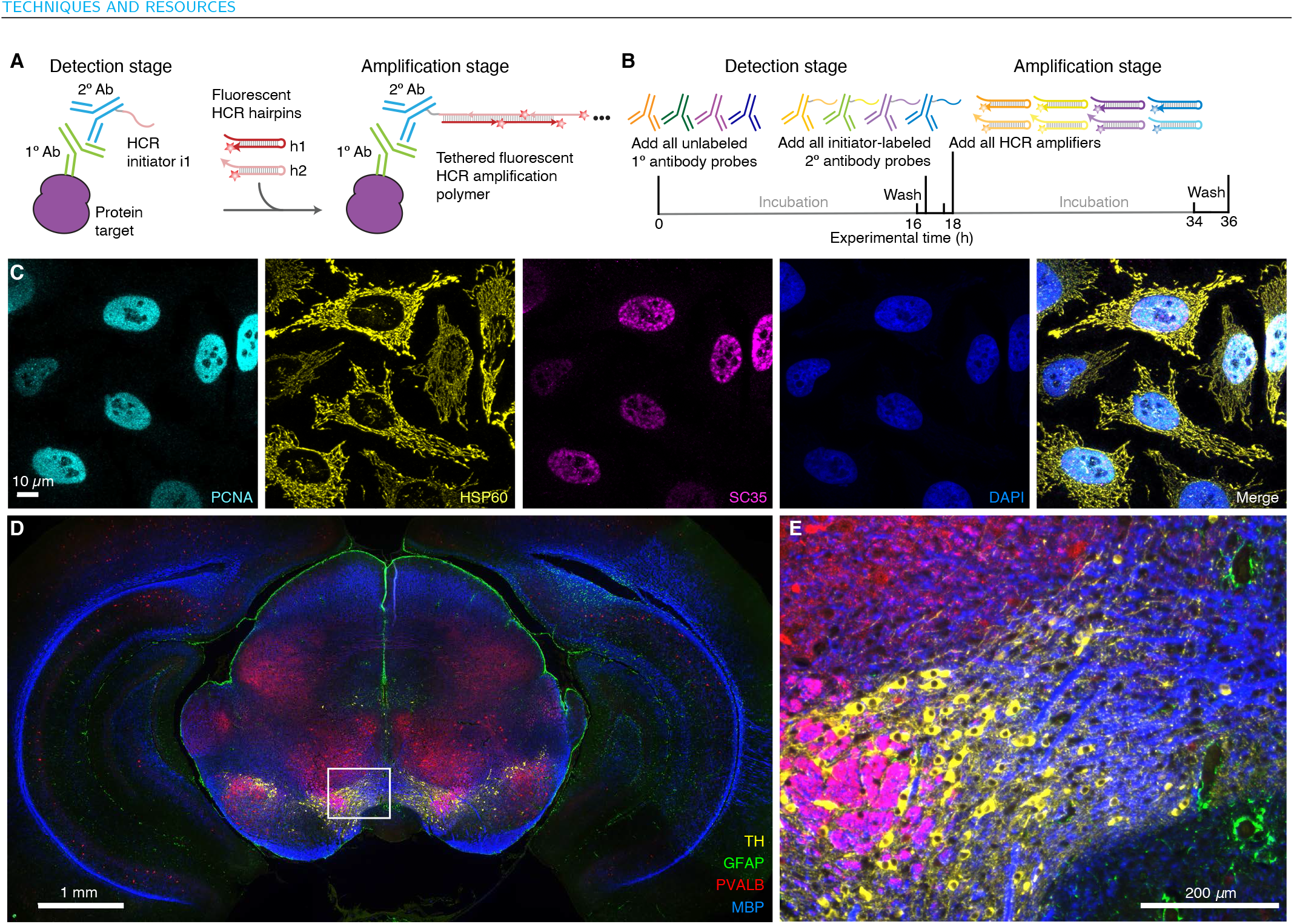
Multiplexed protein imaging via HCR 2°IHC using unlabeled primary antibody probes, initiator-labeled secondary antibody probes, and simultaneous HCR signal amplification for all targets. (A) 2-stage IHC protocol. Detection stage: unlabeled primary antibody probes bind to protein targets; wash; initiator-labeled secondary antibody probes bind to primary antibody probes; wash. Amplification stage: initiators trigger self-assembly of fluorophore-labeled HCR hairpins into tethered fluorescent amplification polymers; wash. (B) Multiplexing timeline. The same 2-stage protocol is used independent of the number of target proteins. (C) Confocal image of 3-plex protein imaging in mammalian cells on a slide; 0.14 × 0.14 *µ*m pixels; maximum intensity z-projection. Target proteins: PCNA (Alexa647), HSP60 (Alexa546), SC35 (Alexa488). Sample: HeLa cells. (D) Epifluorescence image of 4-plex protein imaging in FFPE mouse brain sections; 0.6×0.6 *µ*m pixels. Target proteins: TH (Alexa488), GFAP (Alexa546), PVALB (Alexa647), MBP (Alexa750). (E) Zoom of depicted region of panel D. Sample: FFPE C57BL/6 mouse brain section (coronal); thickness: 5 *µ*m. See Section S5.3 for additional data.

Using HCR signal amplification, the amplification gain corresponds to the number of fluorophore-labeled hairpins per amplification polymer. Hence, we were curious to measure the mean HCR polymer length in the context of HCR 1°IHC and HCR 2°IHC experiments. We can estimate HCR amplification gain by comparing the signal intensity in HCR experiments using h1 and h2 hairpins together (enabling polymerization to proceed as normal) vs using only hairpin h1 (so that each HCR initiator can bind only one HCR hairpin and polymerization cannot proceed). Across 4 measurement scenarios (2 in mammalian cells + 2 in FFPE mouse brain sections; 2 using HCR 1°IHC and 2 using HCR 2°IHC) we observed a median polymer length of ≈180 hairpins (Section S5.5). It is this amplification gain that boosts the signal above autofluorescence to yield high signal-to-background even in FFPE tissues and whole-mount vertebrate embryos.

### qHCR imaging: protein relative quantitation with subcellular resolution in an anatomical context

We previously demonstrated that HCR RNA-ISH overcomes the historical tradeoff between RNA quantitation and anatomical context, enabling mRNA relative quantitation (qHCR imaging) with subcellular resolution within whole-mount vertebrate embryos (Trivedi *et al*., 2018; Choi *et al*., 2018). Here, we demonstrate that HCR IHC enables analogous subcellular quantitation of proteins in an anatomical context. To test protein relative quantitation, we first redundantly detected a target protein using two primary antibody probes that bind different epitopes on the same protein and trigger different spectrally-distinct HCR amplifiers (Figure 4A; top), yielding a 2-channel image (Figure 4B; top). If HCR signal scales approximately linearly with the number of target proteins per voxel, a 2-channel scatter plot of normalized voxel intensities will yield a tight linear distribution with approximately zero intercept (Trivedi *et al*., 2018). Conversely, observing a tight linear distribution with approximately zero intercept (Figure 4C; top), we conclude that the HCR signal scales approximately linearly with the number of target proteins per imaging voxel, after first ruling out potential systematic crowding effects that could permit pairwise voxel intensities to slide undetected along a line (Supplementary Figure S24). Using one initiator-labeled primary antibody probe per channel, we observe high accuracy (linearity with zero intercept) and precision (scatter around the line) for subcellular 2×2 *µ*m voxels within FFPE mouse brain sections. Note that this redundant detection experiment provides a conservative characterization of quantitative performance since there is the risk that two antibody probes may interfere with each other to some extent when attempting to bind different epitopes on the same target protein. As a further test of quantitative imaging characteristics, we detected a protein target with unlabeled primary antibody probes that are subsequently detected by two batches of secondary antibody probes that trigger different spectrally-distinct HCR amplifiers (Figure 4A; bottom). This experiment is testing the accuracy and precision of the secondary antibody probes and HCR signal amplification, but not that of the primary antibody probes. In FFPE human breast tissue sections (Figure 4B; bottom), a 2-channel scatter plot of voxel intensities for subcellular 2×2 *µ*m voxels again reveals a tight linear distribution with approximately zero intercept (Figure 4C; bottom). Based on these two studies, we conclude that qHCR imaging enables accurate and precise relative quantitation of protein targets in an anatomical context with subcellular resolution, just as it does for mRNA targets (Trivedi *et al*., 2018; Choi *et al*., 2018).

**Fig. 4:**
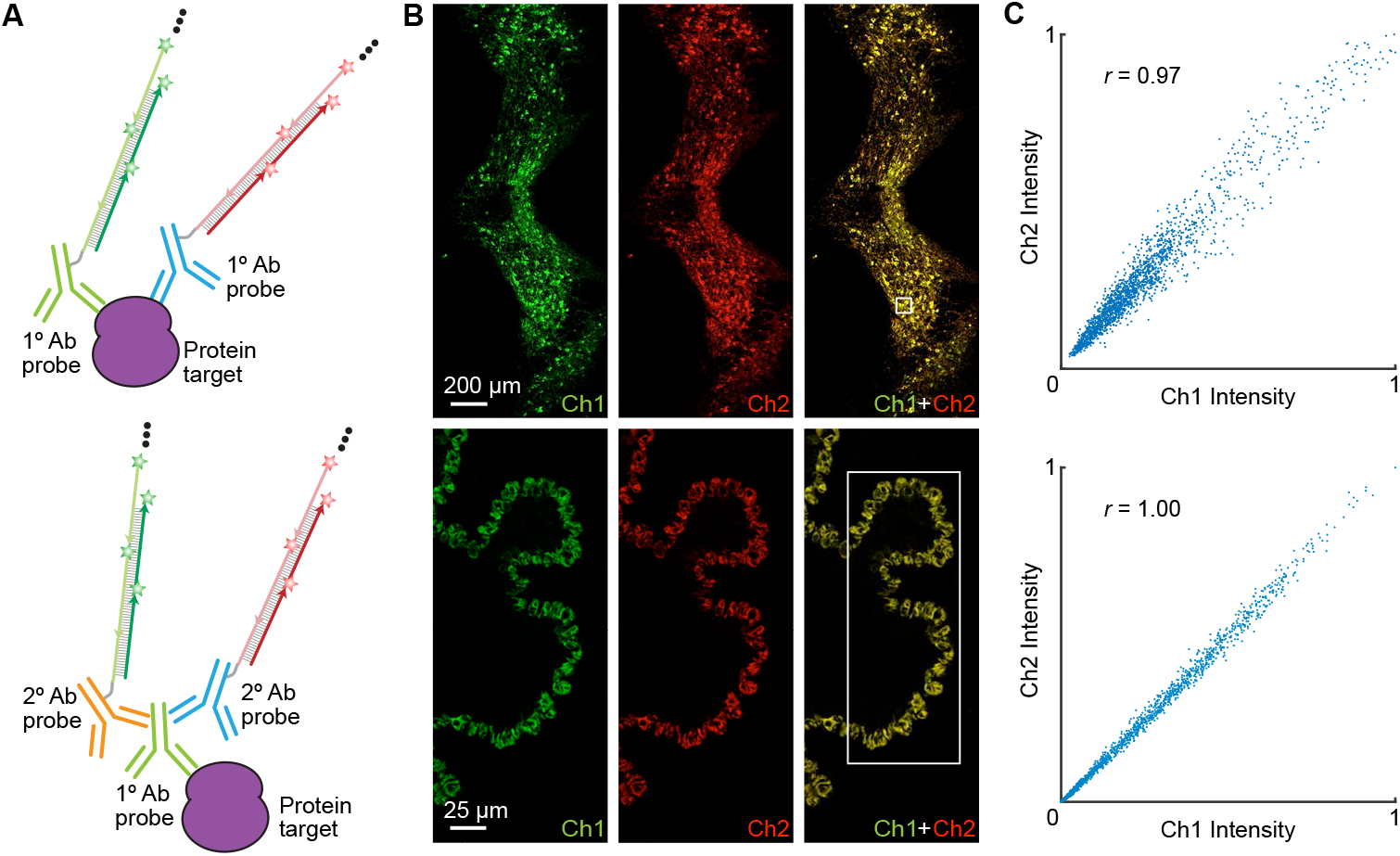
qHCR imaging: protein relative quantitation with subcellular resolution in an anatomical context using HCR 1°IHC or HCR 2°IHC. (A) 2-channel redundant detection of a target protein. Top: target protein detected using two primary antibody probes that bind different epitopes, each initiating an orthogonal spectrally distinct HCR amplifier (Ch1: Alexa647, Ch2: Alexa750). Bottom: target protein detected using unlabeled primary antibody probes and two batches of secondary antibody probes that initiate orthogonal spectrally distinct HCR amplifiers (Ch1: Alexa546, Ch2: Alexa647). (B) Top: Epifluorescence image of FFPE mouse brain section; 0.16×0.16 *µ*m pixels. Target protein: TH. Sample: FFPE C57BL/6 mouse brain section (coronal); thickness: 5 *µ*m. Bottom: Confocal image of FFPE human breast tissue; 0.3 × 0.3 *µ*m pixels; single optical section. Target protein: KRT17. Sample: FFPE human breast tissue section; thickness: 5 *µ*m. (C) High accuracy and precision for protein relative quantitation in an anatomical context. Highly correlated normalized signal (Pearson correlation coefficient, *r*) for subcellular voxels (2 × 2 *µ*m) in the depicted region of panel B. Accuracy: linearity with zero intercept. Precision: scatter around the line. See Section S5.6 for additional data.

### Simultaneous multiplexed protein and RNA imaging using HCR 1° IHC + HCR RNA-ISH or HCR 2°IHC + HCR RNA-ISH

It is important for biologists, drug developers, and pathologists to have the flexibility to image proteins and RNAs simultaneously so as to enable interrogation of both levels of gene expression in the same specimen. Here, we demonstrate that HCR 1°IHC and HCR 2°IHC are both compatible with HCR RNA-ISH, enabling multiplexed quantitative protein and RNA imaging with high signal-to-background. Figure 5 demonstrates HCR 1°IHC + HCR RNA-ISH (2-plex protein + 2-plex RNA) in mammalian cells and FFPE mouse brain sections using initiator-labeled primary antibody probes for protein targets, split-initiator DNA probes for RNA targets, and simultaneous HCR signal amplification for all targets. Figure 6 demonstrates HCR 2°IHC + HCR RNA-ISH (2-plex protein + 2-plex RNA) in mammalian cells and FFPE mouse brain sections using unlabeled primary antibody probes and initiator-labeled secondary antibody probes for protein targets, split-initiator DNA probes for RNA targets, and simultaneous HCR signal amplification for all targets. Across 16 protein and RNA imaging scenarios (8 in mammalian cells + 8 in FFPE mouse brain sections; 8 using 1°IHC HCR + HCR RNA-ISH + 8 using 2°IHC HCR + HCR RNA-ISH; 8 using confocal microscopy + 8 using epifluorescence microscopy), the estimated signal-to-background ratio for each target protein or RNA ranged from 14 to 280 with a median of 43 (see Tables S9 and S11 for additional details).

**Fig. 5:**
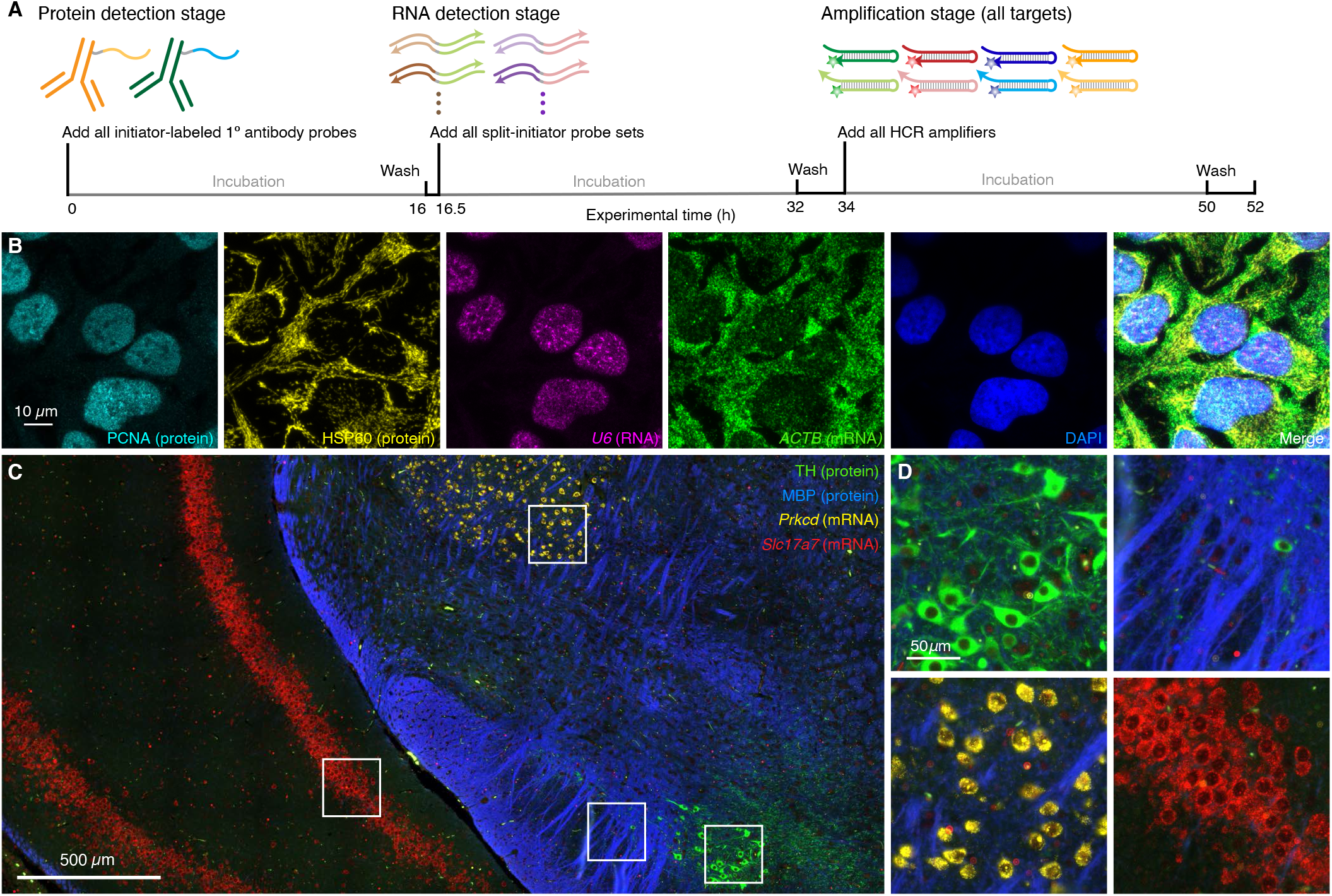
Simultaneous multiplexed protein and RNA imaging via HCR 1° IHC + HCR RNA-ISH using initiator-labeled primary antibody probes for protein targets, split-initiator DNA probes for RNA targets, and simultaneous HCR signal amplification for all targets. (A) 3-stage IHC + RNA-ISH protocol. Protein detection stage: initiator-labeled primary antibody probes bind to protein targets; wash. RNA detection stage: split-initiator DNA probes bind to RNA targets; wash. Amplification stage: initiators trigger self-assembly of fluorophore-labeled HCR hairpins into tethered fluorescent amplification polymers; wash. For multiplexed experiments, the same 3-stage protocol is used independent of the number of target proteins and RNAs. (B) Confocal image of 4-plex IHC + RNA-ISH in mammalian cells on a slide; 0.13 × 0.13 *µ*m pixels; maximum intensity z-projection. Targets: PCNA (protein; Alexa488), HSP60 (protein; Alexa546), *U6* (RNA,; Alexa594), *ACTB* (mRNA; Alexa647). Sample: HeLa cells. (C) Confocal image of 4-plex IHC + RNA-ISH in FFPE mouse brain sections; 0.16 × 0.16 *µ*m pixels. Targets: TH (protein; Alexa488), MBP (protein; Alexa546), *Prkcd* (mRNA; Alexa647), *Slc17a7* (mRNA; Alexa750). Sample: FFPE C57BL/6 mouse brain section (coronal); thickness: 5 *µ*m. (D) Zoom of depicted regions of panel C. See Section S5.7 for additional data.

**Fig. 6:**
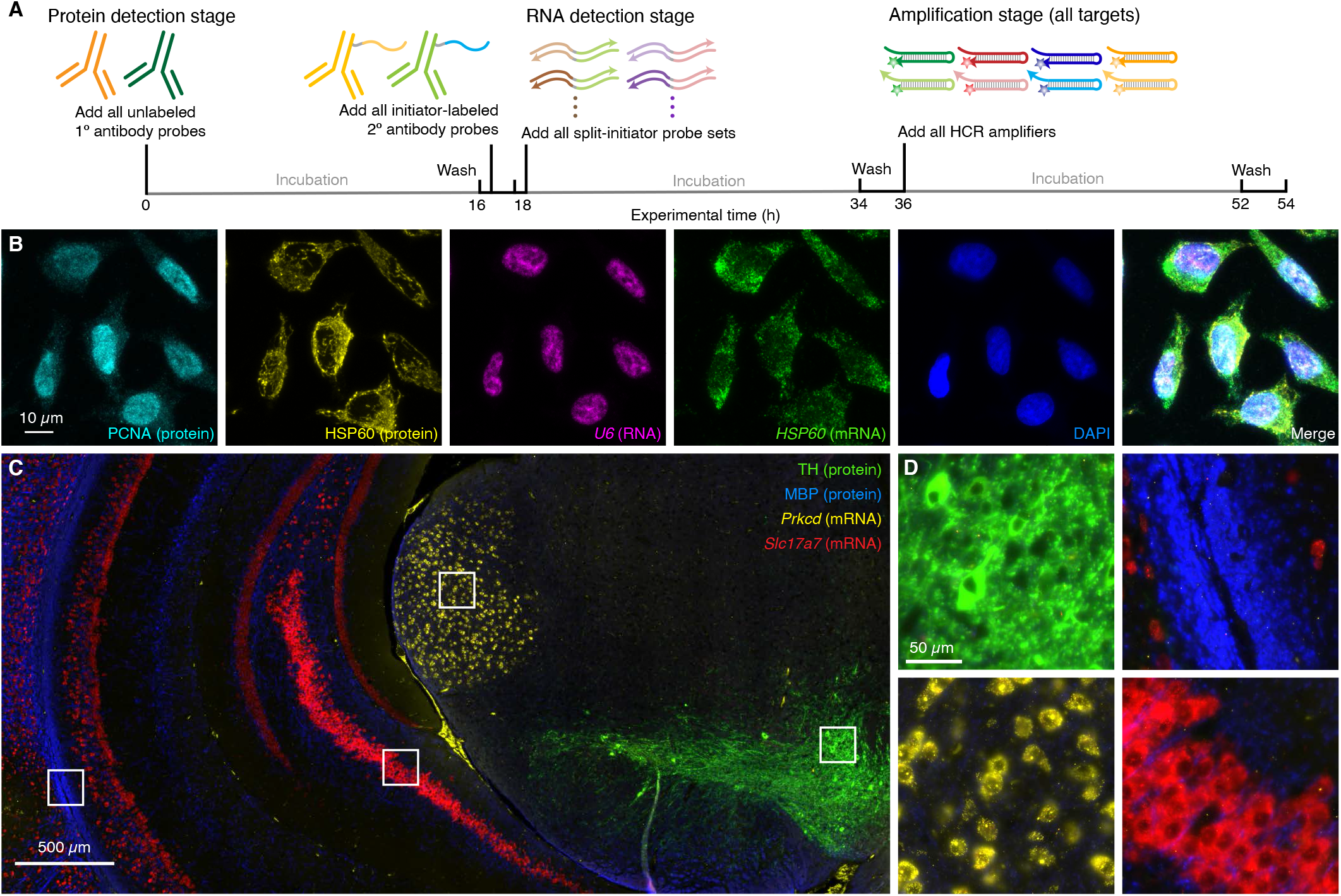
Simultaneous multiplexed protein and RNA imaging via HCR 2°IHC + HCR RNA-ISH using unlabeled primary antibody probes and initiator-labeled secondary antibody probes for protein targets, split-initiator DNA probes for RNA targets, and simultaneous HCR signal amplification for all targets. (A) 3-stage IHC + RNA-ISH protocol. Protein detection stage: unlabeled primary antibody probes bind to protein targets; wash; initiator-labeled secondary antibody probes bind to primary antibody probes; wash. RNA detection stage: split-initiator DNA probes bind to RNA targets; wash. Amplification stage: initiators trigger self-assembly of fluorophore-labeled HCR hairpins into tethered fluorescent amplification polymers; wash. For multiplexed experiments, the same 3-stage protocol is used independent of the number of target proteins and RNAs. (B) Confocal image of 4-plex IHC + RNA-ISH in mammalian cells on a slide; 0.13 × 0.13 *µ*m pixels; maximum intensity z-projection. Targets: PCNA (protein; Alexa488), HSP60 (protein; Alexa546), *U6* (RNA,; Alexa594), *HSP60* (mRNA; Alexa647). Sample: HeLa cells. (C) Confocal image of 4-plex IHC + RNA-ISH in FFPE mouse brain sections; 0.16 × 0.16 *µ*m pixels. Targets: TH (protein; Alexa488), MBP (protein; Alexa546), *Prkcd* (mRNA; Alexa647), *Slc17a7* (mRNA; Alexa750). Sample: FFPE C57BL/6 mouse brain section (coronal); thickness: 5 *µ*m. (D) Zoom of depicted regions of panel C. See Section S5.8 for additional data.

## DISCUSSION

qHCR imaging enables a unified approach to multiplexed quantitative IHC and RNA-ISH. A single experiment yields accurate and precise relative quantitation of both protein and RNA targets with subcellular resolution in the anatomical context of highly autofluorescent samples. Note that no extra work is necessary to perform quantitative imaging – it is a natural property of HCR signal amplification. Here, we validated two complementary approaches for HCR IHC. Using HCR 1°IHC (initiator-labeled primary antibody probes), each target protein in a multiplexed experiment can be detected with antibodies raised in the same host species, which is of-ten convenient based on available antibody libraries. However, antibody-initiator conjugation must be validated for each new primary antibody probe. Alternatively, using HCR 2°IHC (unlabeled primary antibody probes and initiator-labeled secondary antibody probes), each target protein in a multiplexed experiment must be detected with primary antibodies raised in different host species, thus enabling subsequent binding by initiator-labeled secondary antibodies that react with those different host species. This approach has the benefit that a small library of initiator-labeled secondary antibodies can be validated a priori and then used with large libraries of (unmodified) validated primary antibodies, enabling a plug-and-play approach using validated reagents. For simultaneous protein and RNA imaging: during the protein detection stage, M target proteins are detected in parallel; during the RNA detection stage, N target RNAs are detected in parallel; and during the amplification stage, 1-step quantitative HCR signal amplification is performed for all M+N protein and RNA targets simultaneously. In 4-plex experiments in FFPE tissue sections, protein and RNA targets are simultaneously imaged with high signal-to-background in all 4 channels using fluorophores that compete with varying degrees of autofluorescence. For protein imaging using HCR 1°IHC or HCR 2°IHC, we favor protocols with two overnights (Figures 2B and 3B), and for simultaneous protein and RNA imaging using HCR 1°IHC + RNA-ISH or HCR 2°IHC + RNA-ISH, we favor protocols with three overnights (Figures 5A and 6A), allowing researchers to maintain a normal sleep schedule.

HCR RNA-ISH provides automatic background suppression throughout the protocol, ensuring that reagents will not generate amplified background even if they bind non-specifically within the sample (Choi *et al*., 2018). During the detection stage, each RNA target is detected by a probe set comprising one or more pairs of split-initiator probes, each carrying a fraction of HCR initiator i1 (Figure 1B). For a given probe pair, probes that hybridize specifically to their adjacent binding sites on the target RNA colocalize full initiator i1, enabling cooperative initiation of HCR signal amplification. Meanwhile, any individual probes that bind non-specifically in the sample do not colocalize full initiator i1, do not trigger HCR, and thus suppress generation of amplified background. During the amplification stage, automatic background suppression is inherent to HCR hairpins because polymerization is conditional on the presence of the initiator i1; individual h1 or h2 hairpins that bind non-specifically in the sample do not trigger formation of an amplification polymer. For HCR IHC, during the detection stage, each target protein is detected using primary or secondary antibody probes carrying one or more full i1 initiators (Figures 1CD). Hence, if a probe binds non-specifically in the sample, initiator i1 will nonetheless trigger HCR, generating amplified background. As a result, it is important to use antibody probes that are highly selective for their targets, and to wash unused antibody probes from the sample. Nonetheless, during the amplification stage, kinetically trapped HCR hairpins provide automatic background suppression for protein targets just as they do for RNA targets, ensuring that any hairpins that bind non-specifically in the sample do not trigger growth of an HCR amplification polymer. For experiments using HCR IHC + RNA-ISH to image protein and RNA targets simultaneously, RNA targets enjoy automatic background suppression throughout the protocol, while protein targets rely on selective antibody binding to suppress background during the detection stage, combined with automatic background suppression during the amplification stage.

For RNA targets, we have previously shown that multiplexed qHCR imaging enables bi-directional quantitative discovery (Trivedi *et al*., 2018): *read-out* from anatomical space to expression space to discover co-expression relationships in selected regions of the sample; *read-in* from expression space to anatomical space to discover those anatomical locations in which selected gene co-expression relationships occur. Here, by validating high-accuracy, high-precision, high-resolution qHCR imaging for protein targets, read-out/read-in analyses can now be performed for RNA and protein targets simultaneously, offering biologists, drug developers, and pathologists a significantly expanded window for analyzing biological circuits in an anatomical context.

## MATERIALS AND METHODS

### Probes, amplifiers, and buffers

Details on the probes, amplifiers, and buffers for each experiment are displayed in Table S2 for HCR 1°IHC, Table S3 for HCR 2°IHC, and Table S4 for HCR RNA-ISH.

### HCR IHC with/without HCR RNA-ISH

HCR 1°IHC with or without HCR RNA-ISH was performed using the protocols detailed in Sections S3. HCR 2°IHC with or without HCR RNA-ISH was performed using the protocols detailed in Sections S4. Strictly speaking, the cultured cell studies represent immunocytochemistry (ICC) rather than IHC; for notational simplicity, we use the term IHC uniformly in the main text but denote protocols for cultured cells as ICC in the supplementary information. For 5-channel imaging of HeLa cells (Figures 5B, S33–S34, 6B, S37–S38) the above protocols were modified as follows to enable imaging on an upright confocal microscope: cells were grown on a chambered slide with removable chambers (Ibidi, Cat. #81201); prior to imaging, the silicone chambers were removed and cells were mounted with ProLong glass antifade mountant with NucBlue (Thermo Fisher Scientific Cat. #P36981) according to the manufacturer’s instructions.

Experiments were performed in HeLa cells (ATCC Cat. # CRM-CCL-2), human embryonic kidney (HEK) 293T cells (ATCC Cat. # CRL-3216), FFPE C57BL/6 mouse brain sections (coronal; thickness 5 *µ*m, Acepix Biosciences Cat. # 7011-0120), FFPE human breast tissue sections (thickness 5 *µ*m; Acepix Biosciences Cat. # 7310-0620), and whole-mount zebrafish embryos (fixed 27 hpf). Procedures for the care and use of zebrafish embryos were approved by the Caltech IACUC.

### Confocal microscopy

Confocal microscopy was performed using a Zeiss LSM 800 inverted confocal microscope or a Zeiss LSM 880 with Fast Airyscan upright confocal microscope. All confocal images are displayed without background subtraction. See Table S5 for details on the microscope, objective, excitation lasers, beam splitters, emission bandpass filters used for each experiment.

### Epifluorescence microscopy

Epifluorescence microscopy was performed using a Leica THUNDER Imager 3D cell culture epifluorescence microscope equipped with a Leica LED8 multi-LED light source and sCMOS camera (Leica DFC9000 GTC). All epifluorescence images are displayed with instrument noise subtracted but without background subtraction. See Table S6 for details on the objective, excitation wavelengths, and filters used for each experiment.

### Image analysis

Image analysis was performed as detailed in Section S2.6 of the supplementary material including: definition of raw pixel intensities, measurement of signal, background, and signal-to-background, measurement of background components, calculation of normalized subcellular voxel intensities for qHCR imaging.

## Acknowledgments

We thank F. Chen for discussions, M.E. Bronner for reading a draft of the manuscript, and the following resources within the Beckman Institute at Caltech: A. Collazo of the Biological Imaging Facility for assistance with imaging, G. Shin of Molecular Technologies for providing HCR reagents, and the Zebrafish Facility for providing zebrafish embryos.

## Author contributions

Conceptualization: N.A.P.; Methodology: M.S., M.C.L., S.J.S., N.H., H.M.T.C., N.A.P.; Validation: M.S., S.J.S., R.I.; Investigation: M.S., M.C.L, S.J.S., R.I., N.H.; Writing - original draft: N.A.P.; Writing - review & editing: all; Visualization: M.S., R.I., S.J.S.; Supervision: M.C.L., H.M.T.C., and N.A.P.; Project administration: H.M.T.C. and N.A.P.; Funding acquisition: H.M.T.C. and N.A.P.

## Supplementary information

Supplementary information available online.

