## Supplementary Information for "Hybridization chain reaction enables a unified approach to multiplexed, quantitative, high-resolution immunohistochemistry and in situ hybridization"

**References** **114**

### S1 HCR RNA-ISH in diverse organisms and sample types

| Organism | Sample types |
|---|---|
| Human | FFPE thin breast tissue sections (Choi *et al.*, 2016), FFPE thin brain tissue sections (Glineburg *et al.*, 2021), cultured cells on a slide (Choi *et al.*, 2018; Nandagopal *et al.*, 2019; Emert *et al.*, 2021; Glineburg *et al.*, 2021), cultured cell flow cytometry (Choi *et al.*, 2018; Gasperini *et al.*, 2019), thin brain tissue sections (Kamermans *et al.*, 2019), thin intestine tissue sections (May-Zhang *et al.*, 2021), expanded cultured cells on a slide (Alon *et al.*, 2021), thick brain tissue sections (Kumar *et al.*, 2021) |
| Marmoset | thick brain tissue sections (Krienen *et al.*, 2020) |
| Rat | thin rat brain tissue sections (Sui *et al.*, 2016), thick rat brain tissue sections (Chen *et al.*, 2021) |
| Mouse | whole-mount embryos (Huss *et al.*, 2015; Choi *et al.*, 2016), cleared thick brain tissue sections (Shah *et al.*, 2016b; Sylwestrak *et al.*, 2016; Park *et al.*, 2018; Kramer *et al.*, 2018; Kahan *et al.*, 2020; Kumar *et al.*, 2021; Mich *et al.*, 2021), thin brain tissue sections (Shah *et al.*, 2016a; Askary *et al.*, 2020; Ren *et al.*, 2019; Carriere *et al.*, 2020; Young & Song, 2020; Cleary *et al.*, 2021), thin nose tissue sections (Baxter *et al.*, 2021), cultured cells on a slide (Shah *et al.*, 2016b; Nandagopal *et al.*, 2019; Rodriguez *et al.*, 2020; Alon *et al.*, 2021; Glineburg *et al.*, 2021), expanded thick brain tissue sections (Chen *et al.*, 2016; Arshadi *et al.*, 2021; Alon *et al.*, 2021), thick brain tissue sections (Patriarchi *et al.*, 2018; Krienen *et al.*, 2020; Chen *et al.*, 2021; Michael *et al.*, 2020; Mu *et al.*, 2021), whole-mount retina (Anderson *et al.*, 2019), thin spinal cord tissue sections (Li *et al.*, 2020), thin intestine tissue sections (May-Zhang *et al.*, 2021), thin retina tissue sections (Zhuang *et al.*, 2020), thin whole-embryo tissue sections (Liu *et al.*, 2020), thin trigeminal ganglia tissue sections (von Buchholtz *et al.*, 2020), gastruloids (van den Brink *et al.*, 2020), expanded thin brain tissue sections (Alon *et al.*, 2021), thin FFPE liver tissue sections (Wells *et al.*, 2021) |
| Zebrafish | whole-mount embryo (Choi *et al.*, 2010; Choi *et al.*, 2014; Shah *et al.*, 2016b; Trivedi *et al.*, 2018; Ton *et al.*, 2018; Meinecke *et al.*, 2018; Gallagher *et al.*, 2017; Tsai *et al.*, 2020; Cayuso *et al.*, 2019; Wang *et al.*, 2019; Kinney *et al.*, 2020; Howard *et al.*, 2021; Pond *et al.*, 2021), whole-mount larvae (Choi *et al.*, 2016; Andalman *et al.*, 2019; Callahan *et al.*, 2019; Lovett-Barron *et al.*, 2020; O'Brown *et al.*, 2019; Weinberger *et al.*, 2020; Wurster *et al.*, 2021), thin brain tissue sections (O'Brown *et al.*, 2019), FFPE thin heart tissue sections (Simões *et al.*, 2020), |
| Chicken | whole-mount embryos (McLennan *et al.*, 2015; Choi *et al.*, 2016; Lignell *et al.*, 2017; Choi *et al.*, 2018; Ling & Sauka-Spengler, 2019; Williams *et al.*, 2019; Gandhi *et al.*, 2020; Anderson *et al.*, 2020; Gandhi *et al.*, 2021; Mantri *et al.*, 2021), thin whole-embryo tissue sections (Askary *et al.*, 2020; Mantri *et al.*, 2021), thin heart tissue sections (Mantri *et al.*, 2021), thick cochlea tissue sections (Benkafadar *et al.*, 2021; Janesick *et al.*, 2021) |
| Quail | whole-mount embryos (Huss *et al.*, 2019) |
| Xenopus | whole-mount tadpole tails and limbs (Aztekin *et al.*, 2021) |
| Sea urchin | whole-mount embryos (Choi *et al.*, 2016) |
| Axolotl | thin tail tissue sections (Freitas *et al.*, 2019), thin FFPE lung tissue sections (Jensen *et al.*, 2021), limb buds (Schloissnig *et al.*, 2021) |
| Little skate | thin FFPE tissue sections (Marconi *et al.*, 2020; Criswell & Gillis, 2020) |
| Sea lamprey | whole-mount embryos (Hockman *et al.*, 2019) |
| Fruit fly | whole-mount embryos (Choi *et al.*, 2016; Tu *et al.*, 2021; Domsch *et al.*, 2021), whole-mount larvae (Ali *et al.*, 2019), whole-mount brain (Lacin *et al.*, 2019; Michki *et al.*, 2021), whole-mount ovaries (Lacin *et al.*, 2019) |
| Beetle | whole-mount head (Crabtree *et al.*, 2020), whole-mount genitalia (Crabtree *et al.*, 2020), thin thoracic tissue sections (Hu *et al.*, 2019) |
| Deep-sea anemone | thin tissue sections (Goffredi *et al.*, 2021) |
| Basal chordate | whole-mount (Kourakis *et al.*, 2019) |
| Lancelet | whole-mount embryos (Herrera-Úbeda *et al.*, 2019) |
| Blood fluke | whole-mount (Diaz Soria *et al.*, 2020) |
| Nematode | whole-mount larvae (Choi *et al.*, 2016) |
| Bacteria | bacteria on termite gut protozoa (Rosenthal *et al.*, 2013), bacteria in environmental samples (Yamaguchi *et al.*, 2015), cultured bacteria on a slide (Choi *et al.*, 2016), bacteria in cleared sputum (DePas *et al.*, 2016), cultured bacterial flow cytometry (Choi *et al.*, 2018), aggregates (Jorth *et al.*, 2019), bacterial symbionts within whole-mount juvenile squid light organ (Bennett *et al.*, 2020; Moriano-Gutierrez *et al.*, 2020), bacteria on cleared plant roots (Dar *et al.*, 2020), bacteria in cleared whole-mount mouse intestine (Gallego-Hernandez *et al.*, 2020), symbionts of the giant tube worm (Hinzke *et al.*, 2021) |
| Viruses | SARS-CoV-2 in human airway epithelial cultures (Milewska *et al.*, 2020) |
| Multi-kingdom | whole-mount juvenile squid light organ and bacterial symbionts (Nikolakakis *et al.*, 2015), consortia of archaea and bacteria (Metcalfe *et al.*, 2021) |

**Table S1.** Examples of HCR RNA-ISH in diverse organisms and sample types. Tissue sections are classified as "thick" for thickness $\geq$ 50 $\mu$m and "thin" otherwise.

### S2 Additional materials and methods

#### S2.1 Probe and amplifier details for protein targets using HCR 1°IHC

| Species | Sample | Protein target | 1°Ab probe (initiator labeled) | Working concentration ($\mu$g/mL) | Supplier (catalog #) | HCR amplifier | Figures |
|---|---|---|---|---|---|---|---|
| *H. sapiens sapiens* | HeLa cells | HSP60 | 1°mAb rabbit IgG anti-HSP60 | 1 | Ab (ab224528) | B3-Alexa488 | 2C, S1, S2 |
| | HeLa cells | GM130 | 1°mAb rabbit IgG anti-GM130 | 1 | Ab (ab215966) | B2-Alexa647 | 2C, S1, S3 |
| | HeLa cells | SC35 | 1°mAb mouse IgG1 anti-SC35 | 1 | Ab (ab11826) | B4-Alexa546 | 2C, S1, S4 |
| | HeLa cells | HSP60 | 1°mAb rabbit IgG anti-HSP60 | 1 | Ab (ab224528) | B3-Alexa546 | 5B, S33, S34 |
| | HeLa cells | PCNA | 1°mAb mouse IgG2a anti-PCNA | 1 | Ab (ab264494) | B5-Alexa488 | 5B, S33, S34 |
| | HeLa cells | PCNA | 1°mAb mouse IgG2a anti-PCNA | 1 | Ab (ab264494) | B5-Alexa647 | S20 |
| *M. musculus* | brain section | TH | 1°mAb rabbit IgG anti-TH | 1.4 | Ab (ab219729) | B1-Alexa488 | 2DE, S5, S6 |
| | brain section | GFAP | 1°mAb rabbit IgG anti-GFAP | 1.9 | Ab (ab223127) | B3-Alexa546 | 2DE, S5, S7 |
| | brain section | MBP | 1°mAb rabbit IgG anti-MBP | 0.4 | Ab (ab230378) | B5-Alexa647 | 2DE, 5CD, S5, S8, S35, S36 |
| | brain section | MAP2 | 1°mAb rabbit IgG anti-MAP2 | 1 | Ab (ab236033) | B4-Alexa750 | 2DE, S5, S9 |
| | brain section | TH | 1°mAb rabbit IgG anti-TH | 0.2 | Ab (ab219729) | B1-Alexa647 | 4B, S29 |
| | brain section | TH | 1°mAb rabbit IgG anti-TH | 0.1 | Ab (ab220218) | B3-Alexa750 | 4B, S29 |
| | brain section | TH | 1°mAb rabbit IgG anti-TH | 0.2 | Ab (ab219729) | B1-Alexa488 | 5CD, S35, S36 |
| | brain section | TH | 1°mAb rabbit IgG anti-TH | 0.1 | Ab (ab220218) | B3-Alexa647 | S22 |

**Table S2. Organism, sample type, target protein, 1°Ab probe details, HCR amplifier details, and figure numbers for HCR 1°IHC.** For HCR 1°IHC, initiator-labeled primary antibody probes, HCR amplifiers, and buffers (antibody buffer, wash buffer) were obtained from Molecular Technologies (MT) within the Beckman Institute at Caltech (HeLa cells) or from Molecular Instruments (MI) (FFPE mouse brain sections). Ab: Abcam.

#### S2.2 Probe and amplifier details for protein targets using HCR 2°IHC

| Species | Sample | Protein target | 1°Ab probe (unlabeled)<br>2°Ab probe (initiator-labeled) | Working concentration ($\mu$g/mL) | Supplier (catalog #) | HCR amplifier | Figures |
|---|---|---|---|---|---|---|---|
| *H. sapiens sapiens* | HeLa cells | PCNA | 1°mAb mouse IgG2a anti-PCNA | 0.1 | Ab (ab265585) | | |
| | | | 2°pAb goat anti mouse IgG2a-B5 | 1 | MT (A9112-B5) | B5-Alexa647 | 3C, S10, S11, S21, S24, S25, S27, S28 |
| | HeLa cells | HSP60 | 1°mAb rabbit IgG anti-HSP60 | 2.4 | Ab (ab190828) | | |
| | | | 2°pAb donkey anti rabbit IgG-B3 | 1 | MT (A9230-B3) | B3-Alexa546 | 3C, S10, S12 |
| | HeLa cells | SC35 | 1°mAb mouse IgG1 anti-SC35 | 5 | Ab (ab11826) | | |
| | | | 2°pAb goat anti mouse IgG1-B2 | 1 | MT (A9111-B2) | B2-Alexa488 | 3C, S10, S13 |
| | HeLa cells | SC35 | 1°mAb mouse IgG1 anti-SC35 | 5 | Ab (ab11826) | | |
| | | | 2°pAb goat anti mouse IgG1-B2 | 1 | MT (A9111-B2) | B2-Alexa546 | S24–S26, S28 |
| | HeLa cells | PCNA | 1°mAb mouse IgG2a anti-PCNA | 1.26 | CST (2586) | | |
| | | | 2°pAb goat anti mouse IgG2a-B5 | 1 | MT (A9112-B5) | B5-Alexa488 | 6B, S37, S38 |
| | HeLa cells | HSP60 | 1°mAb rabbit IgG anti-HSP60 | 2.4 | Ab (ab190828) | | |
| | | | 2°pAb donkey anti rabbit IgG-B4 | 1 | MT (A9230-B4) | B4-Alexa546 | 6B, S37, S38 |
| | FFPE breast section | KRT17 | 1°pAb rabbit IgG anti-KRT17 | 20 | Ab (ab53707) | | |
| | | | 2°pAb donkey anti rabbit IgG-B4 | 1 | MT (A9230-B4) | B4-Alexa546 | 4B, S30, S32 |
| | FFPE breast section | KRT17 | 1°pAb rabbit IgG anti-KRT17 | 20 | Ab (ab53707) | | |
| | | | 2°pAb donkey anti rabbit IgG-B3 | 1 | MT (A9230-B3) | B3-Alexa647 | 4B, S30, S32 |
| | FFPE breast section | KRT19 | 1°mAb mouse IgG1 anti-KRT19 | 10 | Ab (ab9221) | | |
| | | | 2°pAb goat anti mouse IgG1-B2 | 1 | MT (A9111-B2) | B2-Alexa546 | S31, S32 |
| | FFPE breast section | KRT19 | 1°mAb mouse IgG1 anti-KRT19 | 10 | Ab (ab9221) | | |
| | | | 2°pAb goat anti mouse IgG1-B5 | 1 | MT (A9111-B5) | B5-Alexa647 | S31, S32 |
| *M. musculus* | brain section | TH | 1°pAb sheep IgG2 anti-TH | 0.15 | Ab (ab113) | | |
| | | | 2°pAb donkey anti sheep IgG-B4 | 4 | MI (12-017-01-B4) | B4-Alexa488 | 3DE, 6CD, S14, S15, S39,S40 |
| | brain section | TH | 1°mAb rabbit IgG anti-TH | 0.2 | Ab (ab220218) | | |
| | | | 2°pAb donkey anti rabbit IgG-B3 | 3.4 | MI (12-015-01-B3) | B3-Alexa647 | S23 |
| | brain section | GFAP | 1°pAb chicken IgY anti-GFAP | 0.1 | TFS (PA1-10004) | | |
| | | | 2°pAb donkey anti chicken IgG-B1 | 2.5 | MI (12-018-01-B1) | B1-Alexa546 | 3DE, S14, S16 |
| | brain section | PVALB | 1°mAb rabbit IgG anti-PVALB | 1.1 | Ab (ab243695) | | |
| | | | 2°pAb donkey anti rabbit IgG-B5 | 2.5 | MI (12-015-01-B5) | B5-Alexa647 | 3DE, S14, S17 |
| | brain section | MBP | 1°mAb rat IgG2a anti-MBP | 0.2 | Ab (ab7349) | | |
| | | | 2°pAb donkey anti rat IgG-B3 | 1.1 | MI (12-019-01-B3) | B3-Alexa750 | 3DE, S14, S18 |
| | | | 2°pAb donkey anti rat IgG-B3 | 1.1 | MI (12-019-01-B3) | B3-Alexa546 | 6CD, S39, S40 |
| *D. rerio* | whole-mount | Elavl3/Elavl4 | 1°mAb mouse IgG2b anti-Elavl3/Elavl4 | 5 | TFS (A-21271) | | |
| | 27 hpf embryo | | 2°pAb goat anti-mouse IgG2b-B1 | 1 | MT (A9113-B1) | B1-Alexa647 | S19 |

**Table S3. Organism, sample type, target protein, 1°Ab probe details, 2°Ab probe details, HCR amplifier details, and figure numbers for HCR 2°IHC.** For HCR 2°IHC, initiator-labeled secondary antibody probes, HCR amplifiers, and buffers (antibody buffer, wash buffer) were obtained from Molecular Technologies (MT) within the Beckman Institute at Caltech (HeLa cells) or from Molecular Instruments (MI) (FFPE mouse brain sections). Ab: Abcam. CST: Cell Signaling Technology. TFS: Thermo Fisher Scientific.

#### S2.3 Probe and amplifier details for RNA targets using HCR RNA-ISH

| Species | Sample | RNA target | Split-initiator probe pairs | Supplier (catalog #) | HCR amplifier | Figures |
|---|---|---|---|---|---|---|
| *H. sapiens sapiens* | HeLa cells | *ACTB* | 10 | MT (4226/A506) | B2-Alexa647 | 5B, S33, S34 |
| | HeLa cells | *U6* | 2 | MT (4138/E294) | B1-Alexa594 | 5B, 6B, S33, S34, S37, S38 |
| | HeLa cells | *HSP60* | 18 | MT (4069/E216) | B2-Alexa647 | 6B, S37, S38 |
| *M. musculus* | brain section | *Prkcd* | 31 | MI (PRH342) | B2-Alexa647 | 5CD, S35, S36 |
| | brain section | *Prkcd* | 31 | MI (PRB518) | B1-Alexa647 | 6CD, S39, S40 |
| | brain section | *Slc17a7* | 36 | MI (PRB315) | B4-Alexa750 | 5CD, S35, S36 |
| | brain section | *Slc17a7* | 36 | MI (PRF033) | B2-Alexa750 | 6CD, S39, S40 |

**Table S4. Organism, sample type, target RNA, probe set details, HCR amplifier details, and figure numbers for HCR RNA-ISH.** For HCR RNA-ISH, HCR probe sets, amplifiers, and buffers (probe hybridization buffer, probe wash buffer, amplification buffer) were obtained from Molecular Technologies (MT) within the Beckman Institute at Caltech (HeLa cells) or from Molecular Instruments (MI) (FFPE mouse brain sections).

#### S2.4 Confocal microscope settings

| Sample | Target | Microscope | Objective | Fluorophore | Laser (nm) | Beam splitter | Filter (nm) | Pixel size ($x \times y \times z$ $\mu$m) | Figures |
|---|---|---|---|---|---|---|---|---|---|
| HeLa cells | HSP60 | Zeiss LSM 800 | 63× | Alexa488 | 488 | MBS 405/488/561/640 (T10/R90) | 490–541 | $0.1981 \times 0.1981 \times 0.90$ | 2C, S1, S2 |
| | SC35 | | | Alexa546 | 561 | MBS 405/488/561/640 (T10/R90) | 565–600 | $0.1981 \times 0.1981 \times 0.90$ | 2C, S1, S4 |
| | GM130 | | | Alexa647 | 640 | MBS 405/488/561/640 (T10/R90) | 656–700 | $0.1981 \times 0.1981 \times 0.90$ | 2C, S1, S3 |
| | — | | | DAPI | 405 | MBS 405/488/561/640 (T10/R90) | 410–470 | $0.1981 \times 0.1981 \times 0.90$ | 2C, S1–S4 |
| | SC35 | Zeiss LSM 800 | 63× | Alexa488 | 488 | MBS 405/488/561/640 (T10/R90) | 490–541 | $0.1415 \times 0.1415 \times 0.90$ | 3C, S10, S13 |
| | HSP60 | | | Alexa546 | 561 | MBS 405/488/561/640 (T10/R90) | 565–600 | $0.1415 \times 0.1415 \times 0.90$ | 3C, S10, S12 |
| | PCNA | | | Alexa647 | 640 | MBS 405/488/561/640 (T10/R90) | 656–700 | $0.1415 \times 0.1415 \times 0.90$ | 3C, S10, S11 |
| | — | | | DAPI | 405 | MBS 405/488/561/640 (T10/R90) | 410–470 | $0.1415 \times 0.1415 \times 0.90$ | 3C, S10–S13 |
| | PCNA | Zeiss LSM 800 | 63× | Alexa647 | 640 | MBS 405/488/561/640 (T10/R90) | 656–700 | $0.1981 \times 0.1981 \times 0.90$ | S20, S21 |
| | — | | | DAPI | 405 | MBS 405/488/561/640 (T10/R90) | 410–470 | $0.1981 \times 0.1981 \times 0.90$ | S20, S21 |
| | SC35 | Zeiss LSM 800 | 20× | Alexa546 | 561 | MBS 405/488/561/640 (T10/R90) | 564–610 | $0.312 \times 0.312 \times 2.5$ | S24–S28 |
| | PCNA | | | Alexa647 | 640 | MBS 405/488/561/640 (T10/R90) | 656–700 | $0.312 \times 0.312 \times 2.5$ | S24–S28 |
| | — | | | DAPI | 405 | MBS 405/488/561/640 (T10/R90) | 410–470 | $0.312 \times 0.312 \times 2.5$ | S24–S28 |
| | PCNA | Zeiss LSM 880 | 63× | Alexa488 | 488 | MBS 488/561/633 | 499–554 | $0.1318 \times 0.1318 \times 0.90$ | 5B, S33, S34 |
| | HSP60 | | | Alexa546 | 561 | MBS 488/561/633 | 561–589 | $0.1318 \times 0.1318 \times 0.90$ | 5B, S33, S34 |
| | *U6* | | | Alexa594 | 594 | MBS 458/514/594 | 615–645 | $0.1318 \times 0.1318 \times 0.90$ | 5B, S33, S34 |
| | *ACTB* | | | Alexa647 | 633 | MBS 488/561/633 | 669–696 | $0.1318 \times 0.1318 \times 0.90$ | 5B, S33, S34 |
| | — | | | DAPI | 405 | MBS 405 | 410–471 | $0.1318 \times 0.1318 \times 0.90$ | 5B, S33, S34 |
| | PCNA | Zeiss LSM 880 | 63× | Alexa488 | 488 | MBS 488/561/633 | 499–554 | $0.1318 \times 0.1318 \times 1.0$ | 6B, S37, S38 |
| | HSP60 | | | Alexa546 | 561 | MBS 488/561/633 | 561–589 | $0.1318 \times 0.1318 \times 1.0$ | 6B, S37, S38 |
| | *U6* | | | Alexa594 | 594 | MBS 458/514/594 | 615–645 | $0.1318 \times 0.1318 \times 1.0$ | 6B, S37, S38 |
| | *HSP60* | | | Alexa647 | 633 | MBS 488/561/633 | 669–696 | $0.1318 \times 0.1318 \times 1.0$ | 6B, S37, S38 |
| | — | | | DAPI | 405 | MBS 405 | 410–471 | $0.1318 \times 0.1318 \times 1.0$ | 6B, S37, S38 |
| FFPE human breast section | KRT17 | Zeiss LSM 800 | 20× | Alexa546 | 561 | MBS 405/488/561/640 (T10/R90) | 568–615 | $0.312 \times 0.312 \times 2.5$ | 4B, S30, S32 |
| | KRT17 | | | Alexa647 | 640 | MBS 405/488/561/640 (T10/R90) | 656–700 | $0.312 \times 0.312 \times 2.5$ | 4B, S30, S32 |
| | KRT19 | | | Alexa546 | 561 | MBS 405/488/561/640 (T10/R90) | 568–615 | $0.312 \times 0.312 \times 2.5$ | S31, S32 |
| | KRT19 | | | Alexa647 | 640 | MBS 405/488/561/640 (T10/R90) | 656–700 | $0.312 \times 0.312 \times 2.5$ | S31, S32 |
| Zebrafish embryos | Elavl3/Elavl4 | Zeiss LSM 800 | 20× | Alexa647 | 640 | MBS 405/488/561/640 (T10/R90) | 656–700 | $0.312 \times 0.312 \times 2.5$ | S19 |

**Table S5. Confocal microscope settings.** Confocal microscopy was performed with a Zeiss LSM 800 inverted confocal microscope or a Zeiss LSM880 with Fast Airyscan upright confocal microscope. Objectives were as follows: Plan-Apochromat 63×/1.4 Oil DIC M27 (Zeiss LSM 800), Plan-Apochromat 20×/0.8 M27 (Zeiss LSM 800), Plan-Apochromat 63×/1.4 Oil DIC (Zeiss LSM 880).

#### S2.5 Epifluorescence microscope settings

| Sample | Target | Objective | Fluorophore | LED (nm) | Ex/Em Filter | External filter wheel: emission filter center wavelength/bandwidth (nm) | Pixel size ($x \times y$ $\mu$m) | LED intensity (%) | Exposure time (ms) | Figures |
|---|---|---|---|---|---|---|---|---|---|---|
| FFPE mouse brain section | TH | 20× | Alexa488 | 438 | DFT51011 | 510/40 | 0.3238 × 0.3238 | 60 | 150 | 2DE, S5, S6 |
| | GFAP | | Alexa546 | 555 | DFT51011 | 590/50 | 0.3238 × 0.3238 | 15 | 50 | 2DE, S5, S7 |
| | MBP | | Alexa647 | 640 | DFT51011 | 700/75 | 0.3238 × 0.3238 | 60 | 200 | 2DE, S5, S8 |
| | MAP2 | | Alexa750 | 730 | Y7 | NA | 0.3238 × 0.3238 | 60 | 650 | 2DE, S5, S9 |
| | — | | DAPI | 395 | DFT51011 | 440/40 | 0.3238 × 0.3238 | 60 | 50 | S6, S7, S8, S9 |
| | TH | 10× | Alexa488 | 438 | DFT51011 | 510/40 | 0.6451 × 0.6451 | 60 | 350 | 3DE, S14, S15 |
| | GFAP | | Alexa546 | 555 | DFT51011 | 590/50 | 0.6451 × 0.6451 | 50 | 100 | 3DE, S14, S16 |
| | PVALB | | Alexa647 | 640 | DFT51011 | 700/75 | 0.6451 × 0.6451 | 60 | 250 | 3DE, S14, S17 |
| | MBP | | Alexa750 | 730 | Y7 | NA | 0.6451 × 0.6451 | 80 | 750 | 3DE, S14, S18 |
| | — | | DAPI | 395 | DFT51011 | 440/40 | 0.6451 × 0.6451 | 60 | 50 | S15, S16, S17, S18 |
| | TH | 40× | Alexa647 | 640 | DFT51011 | 700/75 | 0.1612 × 0.1612 | 19 | 100 | 4B, S29 |
| | TH | | Alexa750 | 730 | Y7 | NA | 0.1612 × 0.1612 | 75 | 650 | 4B, S29 |
| | TH | 40× | Alexa488 | 438 | DFT51011 | 510/40 | 0.1619 × 0.1619 | 60 | 250 | 5CD, S35, S36 |
| | MBP | | Alexa546 | 555 | DFT51011 | 590/50 | 0.1619 × 0.1619 | 60 | 250 | 5CD, S35, S36 |
| | *Prkcd* | | Alexa647 | 640 | DFT51011 | 700/75 | 0.1619 × 0.1619 | 60 | 550 | 5CD, S35, S36 |
| | *Slc17a7* | | Alexa750 | 730 | Y7 | NA | 0.1619 × 0.1619 | 91 | 850 | 5CD, S35, S36 |
| | — | | DAPI | 395 | DFT51011 | 440/40 | 0.1619 × 0.1619 | 60 | 50 | S36 |
| | TH | 40× | Alexa488 | 438 | DFT51011 | 510/40 | 0.1619 × 0.1619 | 20 | 150 | 6CD, S39, S40 |
| | MBP | | Alexa546 | 555 | DFT51011 | 590/50 | 0.1619 × 0.1619 | 6 | 50 | 6CD, S39, S40 |
| | *Prkcd* | | Alexa647 | 640 | DFT51011 | 700/75 | 0.1619 × 0.1619 | 80 | 650 | 6CD, S39, S40 |
| | *Slc17a7* | | Alexa750 | 730 | Y7 | NA | 0.1619 × 0.1619 | 70 | 950 | 6CD, S39, S40 |
| | — | | DAPI | 395 | DFT51011 | 440/40 | 0.1619 × 0.1619 | 60 | 150 | S40 |
| | TH | 40× | Alexa647 | 640 | DFT51011 | 700/75 | 0.1619 × 0.1619 | 4 | 100 | S23 |
| | TH | | Alexa647 | 640 | DFT51011 | 700/75 | 0.1619 × 0.1619 | 60 | 100 | S22 |
| | — | | DAPI | 395 | DFT51011 | 440/40 | 0.1619 × 0.1619 | 60 | 50 | S22, S23 |

**Table S6. Epifluorescence microscope settings.** Epifluorescence microscopy was performed with a Leica THUNDER Imager 3D cell culture epifluorescence microscope equipped with a Leica LED8 multi-LED light source and an sCMOS camera (Leica DFC9000 GTC). Objectives were as follows: Leica HC PL APO 20×/0.80, Leica HC PL APO 10×/0.45, Leica HC PL APO 40x/1.30 oil. The DFT51011 filter set comprises: excitation filter with center wavelengths/bandwidths (nm): 391/32, 479/33, 554/24, 638/31; dichroic mirror with wavelengths (nm): 415, 500, 572, 660; emission filter with center wavelengths/bandwidths (nm): 435/30, 519/25, 594/32, 695/58. The DFT51011 filter set is used in conjunction with an external filter wheel as detailed above. The Y7 filter comprises: excitation filter with center wavelength/bandwidth (nm): 710/75; dichroic mirror with wavelength (nm): 750; emission filter with center wavelength/bandwidth (nm): 810/90.

##### S2.6 Image analysis

We build on an image analysis framework developed over a series of publications (Choi *et al.*, 2010; Choi *et al.*, 2014; Choi *et al.*, 2016; Choi *et al.*, 2018; Trivedi *et al.*, 2018). For convenience, here we provide a self-contained description of the details relevant to the present work.

###### S2.6.1 Raw pixel intensities

The total fluorescence within a pixel is a combination of signal, background, and instrument noise. Instrument noise (NOISE) in each channel corresponds to a non-zero reading in the absence of sample.* Fluorescent background (BACK) arises from three sources in each channel:

- autofluorescence (AF): fluorescence inherent to the sample.
- non-specific detection (NSD): probes that bind non-specifically in the sample and subsequently trigger HCR amplification. For experiments that use both primary antibody probes and secondary antibody probes, $\text{NSD}_{1^\circ}$ arises from non-specific binding of primary antibody probes and $\text{NSD}_{2^\circ}$ arises from non-specific binding of secondary antibody probes, with NSD = $\text{NSD}_{1^\circ}$ + $\text{NSD}_{2^\circ}$.
- non-specific amplification (NSA): HCR hairpins that bind non-specifically in the sample.

Fluorescent signal (SIG) in each channel corresponds to:

- signal (SIG): probes that bind specifically to the target and subsequently trigger HCR amplification.

For pixel $i$ of replicate sample $n$, we denote the background

$$X_{n,i}^{\text{BACK}} = X_{n,i}^{\text{NSD}} + X_{n,i}^{\text{NSA}} + X_{n,i}^{\text{AF}}, \tag{S1}$$

the signal:

$$X_{n,i}^{\text{SIG}}, \tag{S2}$$

and the total fluorescence (SIG+BACK+NOISE):

$$X_{n,i}^{\text{SIG+BACK+NOISE}} = X_{n,i}^{\text{SIG}} + X_{n,i}^{\text{BACK}} + X_{n,i}^{\text{NOISE}}. \tag{S3}$$

###### S2.6.2 Measurement of signal, background, noise, and signal-to-background for HCR 1°IHC, HCR 2°IHC, and HCR RNA-ISH

Noise, background, and signal are characterized differently depending on the sample type and microscope type:

- For mouse brain sections imaged using an epifluorescence microscope, noise (NOISE) is characterized for pixels in a representative rectangular region with no sample, background plus noise (BACK+NOISE) is characterized for pixels in a representative rectangular region of no- or low-expression and the combination of signal plus background plus noise (SIG+BACK+NOISE) is characterized for pixels in a representative rectangular region of high expression. All of these measurements can be made based on an experiment of Type 1 (using the terminology of Tables S7A and S8A) using the full protocol with probes and hairpins.
- For cells on a slide imaged using a confocal microscope, instrument noise is negligible so we use the approximation NOISE $\approx 0$. Signal plus background (SIG+BACK) is characterized for pixels in a representative rectangular region of high expression using an experiment of Type 1 (Tables S7A and S8A) employing the full protocol with probes and hairpins. For HCR 1°IHC experiments and HCR RNA-ISH experiments, background (BACK) is characterized for pixels in a representative rectangular region of maximum intensity using the standard protocol but omitting probes (experiment of Type 2 in Table S7B), yielding the partial background

*For experiments using an epifluorescence microscope (mouse brain sections), noise is non-negligible and we take it into account in our analyses. For experiments using a confocal microscope (cells on a slide and whole-mount zebrafish embryos), noise is negligible and we use the approximation NOISE $\approx 0$.

estimate BACK $\approx$ AF+NSA. For HCR 2°IHC experiments, background (BACK) is characterized for pixels in a representative rectangular region of maximum intensity using the standard protocol including secondary antibody probes but omitting primary antibody probes (experiment of Type 4 in Table S8B), yielding the partial background estimate BACK $\approx$ $NSD_{2^\circ}$+AF+NSA.

- For whole-mount zebrafish embryos using a confocal microscope, instrument noise is negligible so we use the approximation NOISE $\approx 0$. Background (BACK) is characterized for pixels in a representative rectangular region of no- or low-expression and the combination of signal plus background plus noise (SIG+BACK) is characterized for pixels in a representative rectangular region of high expression. All of these measurements can be made based on an experiment of Type 1 (using the terminology of Tables S7A and S8A) using the full protocol with probes and hairpins.

For the pixels in these regions, we characterize the distribution by plotting an intensity histogram and characterize average performance by calculating the mean pixel intensities

$$\bar{X}_n^{\text{NOISE}}, \qquad \bar{X}_n^{\text{BACK+NOISE}}, \qquad \bar{X}_n^{\text{SIG+BACK+NOISE}}$$

for replicate embryo $n$. Performance across replicate embryos is characterized by calculating the sample means

$$\bar{X}^{\text{NOISE}}, \qquad \bar{X}^{\text{BACK+NOISE}}, \qquad \bar{X}^{\text{SIG+BACK+NOISE}}$$

and standard error of the means

$$s_{\bar{X}^{\text{NOISE}}}, \qquad s_{\bar{X}^{\text{BACK+NOISE}}}, \qquad s_{\bar{X}^{\text{SIG+BACK+NOISE}}}.$$

The mean background is estimated as

$$\bar{X}^{\text{BACK}} = \bar{X}^{\text{BACK+NOISE}} - \bar{X}^{\text{NOISE}} \tag{S4}$$

with the standard error of the mean estimated via uncertainty propagation as

$$s_{\bar{X}^{\text{BACK}}} \leq \sqrt{(s_{\bar{X}^{\text{BACK+NOISE}}})^2 + (s_{\bar{X}^{\text{NOISE}}})^2}. \tag{S5}$$

The upper bound on estimated standard error holds under the assumption that the correlation between BACK+NOISE and NOISE is non-negative. The mean signal is estimated as

$$\bar{X}^{\text{SIG}} = \bar{X}^{\text{SIG+BACK+NOISE}} - \bar{X}^{\text{BACK+NOISE}} \tag{S6}$$

with the standard error of the mean estimated via uncertainty propagation as

$$s_{\bar{X}^{\text{SIG}}} \leq \sqrt{(s_{\bar{X}^{\text{SIG+BACK+NOISE}}})^2 + (s_{\bar{X}^{\text{BACK+NOISE}}})^2}. \tag{S7}$$

The upper bound on estimated standard error holds under the assumption that the correlation between SIG+BACK+NOISE and BACK+NOISE is non-negative. The signal-to-background ratio is estimated as:

$$\bar{X}^{\text{SIG/BACK}} = \bar{X}^{\text{SIG}} / \bar{X}^{\text{BACK}} \tag{S8}$$

with standard error estimated via uncertainty propagation as

$$s^{\text{SIG/BACK}} \leq \bar{X}^{\text{SIG/BACK}} \sqrt{\left(\frac{s_{\bar{X}^{\text{SIG}}}}{\bar{X}^{\text{SIG}}}\right)^2 + \left(\frac{s_{\bar{X}^{\text{BACK}}}}{\bar{X}^{\text{BACK}}}\right)^2}. \tag{S9}$$

The upper bound on estimated standard error holds under the assumption that the correlation between SIG and BACK is non-negative.

##### S2.6.3 Measurement of background components for HCR 1°IHC and HCR 2°IHC

Calculation of the signal-to-background ratio (Section S2.6.2) requires only a Type 1 experiment (using the terminology of Tables S7A and S8A), yielding the values $\bar{X}^{\text{SIG+BACK+NOISE}}$, $\bar{X}^{\text{BACK+NOISE}}$, and $\bar{X}^{\text{NOISE}}$ that are needed to calculate SIG/BACK. If desired, additional control experiments that omit certain reagents can be used to characterize the individual components of background (AF, NSA, NSD). A Type 2 experiment (no probes, hairpins only) yields $\bar{X}^{\text{NSA+AF+NOISE}}$ and a Type 3 experiment (no probes, no hairpins) yields $\bar{X}^{\text{AF+NOISE}}$ (using the terminology of Tables S7B and S8B). The background components can then be estimated via calculations analogous to (S6) and (S7). The estimated means are:

$$\bar{X}^{\text{NSD}} = \bar{X}^{\text{BACK+NOISE}} - \bar{X}^{\text{NSA+AF+NOISE}} \tag{S10}$$

$$\bar{X}^{\text{NSA}} = \bar{X}^{\text{NSA+AF+NOISE}} - \bar{X}^{\text{AF+NOISE}} \tag{S11}$$

$$\bar{X}^{\text{AF}} = \bar{X}^{\text{AF+NOISE}} - \bar{X}^{\text{NOISE}} \tag{S12}$$

with estimated standard error of the means are:

$$s_{\bar{X}^{\text{NSD}}} \leq \sqrt{(s_{\bar{X}^{\text{BACK+NOISE}}})^2 + (s_{\bar{X}^{\text{NSA+AF+NOISE}}})^2} \tag{S13}$$

$$s_{\bar{X}^{\text{NSA}}} \leq \sqrt{(s_{\bar{X}^{\text{NSA+AF+NOISE}}})^2 + (s_{\bar{X}^{\text{AF+NOISE}}})^2} \tag{S14}$$

$$s_{\bar{X}^{\text{AF}}} \leq \sqrt{(s_{\bar{X}^{\text{AF+NOISE}}})^2 + (s_{\bar{X}^{\text{NOISE}}})^2}. \tag{S15}$$

These upper bounds on estimated standard errors hold under the assumption that the correlations are non-negative for the components being subtracted in the calculation of the mean.

For HCR 1°IHC experiments that employ initiator-labeled primary antibody probes (and no secondary antibody probes), we have by construction $X^{\text{NSD}_{2^\circ}} = 0$ and $X^{\text{NSD}_{1^\circ}} = X^{\text{NSD}}$, with all of the NSD background attributable to primary antibody probes.

For HCR 2°IHC experiments that employ unlabeled primary antibodies and initiator-labeled secondary antibody probes, NSD background arises from both primary and secondary antibody probes:

$$\bar{X}^{\text{NSD}} = \bar{X}^{\text{NSD}_{1^\circ}} + \bar{X}^{\text{NSD}_{2^\circ}}. \tag{S16}$$

A Type 4 experiment (Table S8B; no primary antibody probes, with initiator-labeled secondary antibody probes, with hairpins) yields $\bar{X}^{\text{NSD}_{2^\circ}\text{+NSA+AF+NOISE}}$. The estimated mean for $\text{NSD}_{2^\circ}$ is:

$$\bar{X}^{\text{NSD}_{2^\circ}} = \bar{X}^{\text{NSD}_{2^\circ}\text{+NSA+AF+NOISE}} - \bar{X}^{\text{NSA+AF+NOISE}} \tag{S17}$$

with estimated standard error of the mean:

$$s_{\bar{X}^{\text{NSD}_{2^\circ}}} \leq \sqrt{(s_{\bar{X}^{\text{NSD}_{2^\circ}\text{+NSA+AF+NOISE}}})^2 + (s_{\bar{X}^{\text{NSA+AF+NOISE}}})^2} \tag{S18}$$

The estimated mean for $\text{NSD}_{1^\circ}$ is then:

$$\bar{X}^{\text{NSD}_{1^\circ}} = \bar{X}^{\text{NSD+NSA+AF+NOISE}} - \bar{X}^{\text{NSD}_{2^\circ}\text{NSA+AF+NOISE}} \tag{S19}$$

with estimated standard error of the mean:

$$s_{\bar{X}^{\text{NSD}_{1^\circ}}} \leq \sqrt{(s_{\bar{X}^{\text{NSD+NSA+AF+NOISE}}})^2 + (s_{\bar{X}^{\text{NSD}_{2^\circ}\text{+NSA+AF+NOISE}}})^2}. \tag{S20}$$

For a given quantity, if $\bar{X} < s_{\bar{X}}$, we instead report $\max(\bar{X}, 0) + s_{\bar{X}}$ as an estimated upper bound, and use this bound for uncertainty propagation.

If a Type 1 experiment demonstrates SIG $\gg$ BACK, as is typically the case using HCR imaging, then there is little motivation to perform the other experiment Types to characterize the individual background components (AF, NSA, NSD) as these are all bounded above by BACK.

##### S2.6.4 Measurement of HCR amplification gain (i.e., amplification polymer length)

To estimate HCR amplification gain (corresponding to the number of HCR hairpins per amplification polymer), an additional experiment type can be performed using h1 hairpins only (Type 4 in Table S7A; Type 5 in S8A) to yield $\bar{X}^{\text{SIG}_{\text{h1}}\text{+BACK+NOISE}}$. HCR polymerization cannot proceed without hairpin h2 so each HCR initiator can tether only a single fluorescent h1 hairpin, corresponding to unamplified signal $\text{SIG}_{\text{h1}}$. The mean $\text{SIG}_{\text{h1}}$ is estimated as:

$$\bar{X}^{\text{SIG}_{\text{h1}}} = \bar{X}^{\text{SIG}_{\text{h1}}+\text{BACK}+\text{NOISE}} - \bar{X}^{\text{BACK}+\text{NOISE}} \tag{S21}$$

with estimated standard error of the mean:

$$s_{\bar{X}^{\text{SIG}_{\text{h1}}}} \leq \sqrt{(s_{\bar{X}^{\text{SIG}_{\text{h1}}+\text{BACK}+\text{NOISE}}})^2 + (s_{\bar{X}^{\text{BACK}+\text{NOISE}}})^2} \tag{S22}$$

The upper bound on estimated standard error holds under the assumption that the correlation between $\text{SIG}_{\text{h1}}$+BACK+NOISE and BACK+NOISE is non-negative. The ratio of amplified to unamplified signal provides an estimate of mean HCR polymer length:

$$\bar{X}^{\text{SIG}/\text{SIG}_{\text{h1}}} = \bar{X}^{\text{SIG}}/\bar{X}^{\text{SIG}_{\text{h1}}} \tag{S23}$$

with standard error estimated via uncertainty propagation as

$$s^{\text{SIG}/\text{SIG}_{\text{h1}}} \leq \bar{X}^{\text{SIG}/\text{SIG}_{\text{h1}}} \sqrt{\left(\frac{s_{\bar{X}^{\text{SIG}}}}{\bar{X}^{\text{SIG}}}\right)^2 + \left(\frac{s_{\bar{X}^{\text{SIG}_{\text{h1}}}}}{\bar{X}^{\text{SIG}_{\text{h1}}}}\right)^2}. \tag{S24}$$

The upper bound on estimated standard error holds under the assumption that the correlation between SIG and $\text{SIG}_{\text{h1}}$ is non-negative.

| | Experiment type | Quantity | Reagents: 1° Ab-init | Reagents: Hairpins | Exression region in tissue |
|---|---|---|---|---|---|
| **A** | 1 | SIG+NSD+NSA+AF+NOISE = SIG+BACK+NOISE | ✓ | ✓ | high |
| | 1 | NSD+NSA+AF+NOISE = BACK+NOISE | ✓ | ✓ | no/low |
| | 1 | NOISE | | | no sample |
| **B** | 2 | NSA+AF+NOISE | | ✓ | high |
| | 3 | AF+NOISE | | | high |
| **C** | 4 | $SIG_{h1}$+NSD+NSA+AF+NOISE = $SIG_{h1}$+BACK+NOISE | ✓ | h1 only | high |

**Table S7. Experiment types for HCR IHC using initiator-labeled primary antibody probes.** (A) Characterize signal, background, noise, and signal-to-background. (B) Characterize components of background (AF, NSA, NSD). (C) Characterize unamplified signal and polymer legnth.

| | Experiment type | Quantity | Reagents: 1° Ab | Reagents: 2° Ab-init | Reagents: Hairpins | Expression region in tissue |
|---|---|---|---|---|---|---|
| **A** | 1 | SIG+NSD+NSA+AF+NOISE = SIG+BACK+NOISE | ✓ | ✓ | ✓ | high |
| | 1 | NSD+NSA+AF+NOISE = BACK+NOISE | ✓ | ✓ | ✓ | no/low |
| | 1 | NOISE | | ✓ | ✓ | no sample |
| **B** | 2 | NSA+AF+NOISE | | | ✓ | high |
| | 3 | AF+NOISE | | | | high |
| | 4 | $NSD_{2^\circ}$+NSA+AF+NOISE | | ✓ | ✓ | high |
| **C** | 5 | $SIG_{h1}$+NSD+NSA+AF+NOISE = $SIG_{h1}$+BACK+NOISE | ✓ | ✓ | h1 only | high |

**Table S8. Experiment types for HCR IHC using unlabeled primary antibody probes and initiator-labeled secondary antibody probes.** (A) Characterize signal, background, noise, and signal-to-background. (B) Characterize components of background (AF, NSA, $NSD_{1^\circ}$, $NSD_{2^\circ}$, NSD). (C) Characterize unamplified signal and polymer legnth.

##### S2.6.5 Normalized voxel intensities for qHCR imaging: protein relative quantitation with subcellular resolution in an anatomical context

For quantitative imaging using in situ HCR, precision increases with voxel size as long as the imaging voxels remain smaller than the features in the expression pattern (see Section S2.2 of (Trivedi *et al.*, 2018)). To increase precision, we calculate raw voxel intensities by averaging neighboring pixel intensities while still maintaining a subcellcular voxel size. To facilitate relative quantitation between voxels, we estimate the normalized HCR signal of voxel $j$ in replicate $n$ as:

$$x_{n,j} \equiv \frac{X_{n,j}^{\mathrm{SIG+BACK+NOISE}} - X^{\mathrm{BOT}}}{X^{\mathrm{TOP}} - X^{\mathrm{BOT}}}, \tag{S25}$$

which translates and rescales the data so that the voxel intensities in each channel fall in the interval [0,1]. Here,

$$X^{\mathrm{BOT}} \equiv \bar{X}^{\mathrm{BACK+NOISE}} \tag{S26}$$

is the mean background plus noise across replicates (see Section S2.6.2) and

$$X^{\mathrm{TOP}} \equiv \max_{n,j} X_{n,j}^{\mathrm{SIG+BACK+NOISE}} \tag{S27}$$

is the maximum total fluorescence for a voxel across replicates.

Pairwise expression scatter plots that each display normalized voxel intensities for two channels (e.g., Figures 4 and 5 of (Trivedi *et al.*, 2018)) provide a powerful quantitative framework for performing multidimensional read-out/read-in analyses (Figure 6 of (Trivedi *et al.*, 2018)). Read-out from anatomical space to expression space enables discovery of expression clusters of voxels with quantitatively related expression levels and ratios (amplitudes and slopes in the expression scatter plots), while read-in from expression space to anatomical space enables discovery of the corresponding anatomical locations of these expression clusters within the embryo. The simple and practical normalization approach of (S25)–(S27) translates and rescales all voxels identically within a given channel (enabling comparison of amplitudes and slopes in scatter plots between replicates), and does not attempt to remove scatter in the normalized signal estimate that is caused by scatter in background or noise.

To validate relative protein quantitation with subcellular resolution ($2\times2$ $\mu$m voxels) in FFPE mouse brain sections and FFPE human breast tissue sections, Figures 4C, S29C, S30C, and S31C display highly correlated normalized voxel intensities for 2-channel redundant detection of different protein targets. In this setting, accuracy corresponds to linearity with zero intercept, and precision corresponds to scatter around the line (Trivedi *et al.*, 2018).

### S3 Protocols for HCR 1°IHC with/without HCR RNA-ISH

#### S3.1 Protocols for mammalian cells on a chambered slide

##### S3.1.1 Preparation of fixed mammalian cells on a chambered slide

1. Coat bottom of each chamber by applying 300 $\mu$L of 0.01% poly-D-lysine prepared in cell culture grade $H_2O$.
   NOTE: *A volume of 300 $\mu$L is sufficient per chamber on an 8-chamber slide. Scale volume accordingly if using a different slide format.*
2. Incubate for at least 30 min at room temperature.
3. Aspirate the coating solution and wash each chamber twice with molecular biology grade $H_2O$.
4. Plate desired number of cells in each chamber.
5. Grow cells to desired confluency for 24–48 h.
6. Aspirate growth media and wash each chamber with 300 $\mu$L of DPBS.
   NOTE: *avoid using calcium chloride and magnesium chloride in DPBS as this leads to increased autofluorescence.*
7. Add 300 $\mu$L of 4% formaldehyde to each chamber.
   CAUTION: *use formaldehyde with extreme care as it is a hazardous material.*
8. Incubate for 10 min at room temperature.
9. Remove fixative and wash each chamber 2 $\times$ 300 $\mu$L of DPBS.
10. Aspirate DPBS and add 300 $\mu$L of ice-cold 70% ethanol (EtOH).
11. Permeabilize cells overnight (or longer) at -20 °C.
12. Proceed to HCR assay.

##### S3.1.2 Multiplexed HCR 1°ICC with/without HCR RNA-ISH using initiator-labeled primary antibody probes for protein targets, spit-initiator DNA probes for RNA targets, and simultaneous HCR signal amplification for all targets

**Protein detection stage**

1. Aspirate EtOH and wash samples 2 × 5 min with 300 $\mu$L of 1× PBS.

2. Apply 300 $\mu$L antibody buffer to each chamber. Incubate at room temperature for 1 h with gentle agitation.

3. Prepare working concentration of primary antibodies in antibody buffer. Prepare 300 $\mu$L per chamber.
NOTE: *follow manufacturer's guidelines for primary antibody working concentration.*

4. Replace antibody solution with primary antibody solution and incubate overnight (>12 h) at 4 °C with gentle agitation.
NOTE: *Incubation may be optimized (e.g., 1–2 h at room temperature) depending on sample type and thickness.*

5. Remove excess antibodies by washing 3 × 5 min with PBST at room temperature with gentle agitation.

6. Proceed to **RNA detection stage** for co-detection of protein and RNA. Otherwise, proceed to **Amplification stage**.

**RNA detection stage**

1. Post-fix sample with 300 $\mu$L of 4% formaldehyde.
CAUTION: *use formaldehyde with extreme care as it is a hazardous material.*

2. Incubate for 10 min at room temperature.

3. Remove fixative and wash each chamber 2 × 300 $\mu$L of PBS.

4. Wash sample with 300 $\mu$L of 2× SSC.

5. Pre-hybridize samples in 300 $\mu$L of probe hybridization buffer for 30 min at 37 °C.
CAUTION: *Probe hybridization buffer contains formamide, a hazardous material.*
NOTE: *pre-heat probe hybridization buffer to 37 °C before use.*

6. Prepare a 16 nM probe solution by adding 4.8 pmol of each probe mixture (e.g. 4.8 $\mu$L of 1 $\mu$M stock) to 300 $\mu$L of probe hybridization buffer at 37 °C.
NOTE: *This is the amount of probe set needed for each target on a single chamber of an 8-well chambered slide using 300 $\mu$L of incubation volume.*

7. Remove the pre-hybridization solution and add the probe solution.

8. Incubate samples overnight (>12 h) at 37 °C.

9. Remove excess probes by washing 4 × 5 min with 300 $\mu$L of probe wash buffer at 37 °C.
CAUTION: *Probe wash buffer contains formamide, a hazardous material.*
NOTE: *pre-heat probe wash buffer to 37 °C before use.*

10. Wash samples 5 × 5 min with 5× SSCT at room temperature.

11. Proceed to **Amplification stage**.

**Amplification stage**

1. Wash once with 300 $\mu$L 5$\times$ SSCT at room temperature for 5 min.

2. Pre-amplify samples in 300 $\mu$L of amplification buffer for 30 min at room temperature.
   NOTE: *equilibrate amplification buffer to room temperature before use.*

3. Separately prepare 18 pmol of hairpin h1 and 18 pmol of hairpin h2 by snap cooling 6 $\mu$L of 3 $\mu$M stock (heat at 95 °C for 90 seconds and cool to room temperature in a dark drawer for 30 min).
   NOTE: *HCR hairpins h1 and h2 are provided in hairpin storage buffer ready for snap cooling. h1 and h2 should be snap cooled in separate tubes. This is the amount of hairpins needed for each target in a single sample using 300 $\mu$L of incubation volume.*

4. Prepare a 60 nM hairpin solution by adding all snap-cooled h1 hairpins and snap-cooled h2 hairpins to 300 $\mu$L of amplification buffer at room temperature per sample.

5. Remove the pre-amplification solution and add the hairpin solution.

6. Incubate the slide overnight (>12 h) in the dark at room temperature.

7. Remove excess hairpins by washing 5 $\times$ 5 min with 300 $\mu$L of 5$\times$ SSCT at room temperature.

**Sample mounting for microscopy**

1. Remove final wash and add 150 $\mu$L of mounting medium (e.g., Fluoromount-G with DAPI).

2. Slides can be stored at 4 °C protected from light prior to imaging.
   NOTE: *see Section S2.4 for details of confocal microscopes used to image mammalian cells on a chambered slide.*

##### S3.1.3 Buffers for HCR 1°IHC with/without HCR RNA-ISH

HCR probes (initiator-labeled antibody probes, split-initiator DNA probes), amplifiers, and buffers (antibody buffer, probe hybridization buffer, probe wash buffer, amplification buffer) are available from Molecular Instruments (`www.molecularinstruments.com`). Probe hybridization buffer, and probe wash buffer should be stored at -20 °C. Antibody buffer and amplification buffer should be stored at 4 °C. Make sure all solutions are well mixed before use.

| | |
|---|---|
| **1× PBST** | For 40 mL of solution |
| 1× phosphate buffered solution (PBS) | 4 mL of 10× PBS |
| 0.1% Tween 20 | 400 $\mu$L of 10% Tween 20 |
| | Fill up to 40 mL with ultrapure $H_2O$ |

| | |
|---|---|
| **5× SSCT** | For 40 mL of solution |
| 5× sodium chloride sodium citrate (SSC) | 10 mL of 20× SSC |
| 0.1% Tween 20 | 400 $\mu$L of 10% Tween 20 |
| | Fill up to 40 mL with ultrapure $H_2O$ |

##### S3.1.4 Reagents and supplies

ibidi $\mu$-slide ibitreat (ibidi Cat. # 80826)
Poly-D-lysine hydrobromide (Sigma-Aldrich Cat. # P7280)
Molecular biology grade $H_2O$ (Corning Cat. # 46-000-CV)
DPBS, no calcium, no magnesium (Life Technologies Cat. # 14190144)
Image-iT Fixative Solution 4% (Thermo Fisher Scientific Cat. # FB002)
10× PBS (Ambion Cat. # AM9624)
10% Tween 20 (Teknova Cat. # T0710)
20× sodium chloride sodium citrate (SSC) (Life Technologies Cat. # 15557-044)
DAPI Fluoromount-G (SouthernBiotech Cat. # 0100-20)

#### S3.2 Protocols for FFPE mouse brain tissue sections

##### S3.2.1 Preparation of formalin-fixed paraffin-embedded (FFPE) mouse brain tissue sections

1. Bake slides in a dry oven for 1 h at 60 °C to improve sample adhesion to the slide.

2. In a fume hood, deparaffinize FFPE tissue by immersing in Pro-Par Clearant for 2 × 5 min. Move slides up and down occasionally.
   CAUTION: *use Pro-Par Clearant with care as it is a hazardous material.*
   NOTE: *Xylene can be used in place of Pro-Par Clearant.*
   NOTE: *Each 50 mL tube can fit two outward-facing slides. A volume of 30 mL is sufficient to immerse sections in the tube. If desired, a larger number of slides can be processed together using a Coplin jar.*

3. Incubate slides in 100% ethanol (EtOH) for 2 × 2 min at room temperature. Move slides up and down occasionally.

4. Rehydrate with a series of graded EtOH washes at room temperature.
   (a) 95% EtOH for 3 min
   (b) 70% EtOH for 3 min
   (c) 50% EtOH for 3 min
   (d) Nanopure water for 3 min

5. Bring 500 mL of 1× citrate buffer (pH 6.0) in a beaker to boil in a microwave.
   NOTE: *1× Tris-EDTA buffer (pH 9.0) can be used in place of citrate buffer (pH 6.0). Optimal antigen retrieval method may differ depending on the antigen/antibody used.*

6. Maintain citrate buffer at 90–95 °C on a hot plate.

7. Immerse slides for 15 min.
   NOTE: *Alternatively, slides may be immersed at 95–99 °C for 15 min in a steamer.*

8. Remove beaker from hot plate and add 100 mL of nanopure water every 5 min to allow temperature to decrease to 45 °C in 20 min.

9. Immerse slides in 400 mL of nanopure water in a separate container for 10 min at room temperature.

10. Immerse slides in 1× PBST for 2 × 2 min at room temperature.
    NOTE: *avoid using calcium chloride and magnesium chloride in PBS as this leads to increased autofluorescence in the tissue.*

11. Drain slide by blotting edges on a Kimwipe.

12. Wipe around the section with a Kimwipe and circle tissue with a hydrophobic pen.

13. Proceed to HCR assay.

###### S3.2.2 Buffer recipes for sample preparation

| | |
|---|---|
| **1× citrate buffer** | For 500 mL of solution |
| 1× citrate buffer | 5 mL of 100× citrate buffer (pH 6.0) |
| | Fill up to 500 mL with water |
| **1× Tris-EDTA buffer** | For 500 mL of solution |
| 1× Tris-EDTA buffer | 5 mL of 100× Tris-EDTA buffer (pH 9.0) |
| | Fill up to 500 mL with water |
| **PBST** | For 50 mL of solution |
| 1× PBS | 5 mL of 10× PBS |
| 0.1% Tween 20 | 500 $\mu$L of 10% Tween 20 |
| | Fill up to 50 mL with ultrapure $H_2O$ |

##### S3.2.3 Multiplexed HCR 1°IHC with/without HCR RNA-ISH using initiator-labeled primary antibody probes for protein targets, spit-initiator DNA probes for RNA targets, and simultaneous HCR signal amplification for all targets

**Protein detection stage**

1. Block tissue by applying 200 $\mu$L of antibody buffer on top of the sample. Incubate at room temperature for 1 h in a humidified chamber.
2. Prepare working concentration of initiator-labeled primary antibodies in antibody buffer. Prepare 100 $\mu$L per section.
   NOTE: *follow manufacturer's guidelines for primary antibody working concentration.*
3. Drain slide by blotting edges on a Kimwipe and wipe around the section with another Kimwipe.
4. Add primary antibody solution to each section and incubate overnight (>12 h) at 4 °C in a humidified chamber.
   NOTE: *Incubation may be optimized (e.g., 1–2 h at room temperature) depending on sample type and thickness.*
5. Remove excess antibodies by immersing slide in 1× PBST at room temperature for 3 × 5 min.
6. Proceed to **RNA detection stage** for co-detection of protein and RNA. Otherwise, proceed to **Amplification stage**.

**RNA detection stage**

1. Drain slide by blotting edges on a Kimwipe and wipe around the section with another Kimwipe.
2. Post-fix sample with 200 $\mu$L of 4% formaldehyde on the tissue.
   CAUTION: *use formaldehyde with extreme care as it is a hazardous material.*
3. Incubate slides for 10 min at room temperature.
4. Immerse slides for $2 \times 5$ min in PBST.
5. Immerse slides for 5 min in $5\times$ SSCT.
6. Pre-warm a humidified chamber to 37 °C.
7. Drain slide by blotting edges on a Kimwipe and wipe around the section with another Kimwipe.
8. Add 200 $\mu$L of probe hybridization buffer on top of the tissue sample.
   CAUTION: *Probe hybridization buffer contains formamide, a hazardous material.*
   NOTE: *pre-heat probe hybridization buffer to 37 °C before use.*
9. Pre-hybridize for 10 min inside the humidified chamber.
10. Prepare a 16 nM probe solution by adding 1.6 pmol of each probe set (e.g. 1.6 $\mu$L of 1 $\mu$M stock) to 100 $\mu$L of probe hybridization buffer at 37 °C.
    NOTE: *This is the amount of probe set needed for each target on a single slide using 100 $\mu$L of incubation volume.*
11. Remove the pre-hybridization solution and drain excess buffer on slide by blotting edges on a Kimwipe.
12. Add 100 $\mu$L of the probe solution on top of the tissue sample.
13. Place a coverslip on the sample and incubate overnight (>12 h) in the 37 °C humidified chamber.
14. Immerse slide in probe wash buffer at 37 °C to float off coverslip.
    CAUTION: *Probe wash buffer contains formamide, a hazardous material.*
15. Remove excess probes by incubating slide at 37 °C in:
    (a) 75% of probe wash buffer / 25% $5\times$ SSCT for 15 min
    (b) 50% of probe wash buffer / 50% $5\times$ SSCT for 15 min
    (c) 25% of probe wash buffer / 75% $5\times$ SSCT for 15 min
    (d) 100% $5\times$ SSCT for 15 min

    NOTE: *Wash solutions should be pre-heated to 37 °C before use.*
16. Proceed to amplification stage.

**Amplification stage**

1. Immerse slide in 5× SSCT at room temperature for 5 min.

2. Drain slide by blotting edges on a Kimwipe and wipe around the section with another Kimwipe.

3. Add 200 $\mu$L of amplification buffer on top of the tissue sample and pre-amplify in a humidified chamber for 30 min at room temperature.
   NOTE: *equilibrate amplification buffer to room temperature before use.*

4. Separately prepare 6 pmol of hairpin h1 and 6 pmol of hairpin h2 by snap cooling 2 $\mu$L of 3 $\mu$M stock (heat at 95 °C for 90 seconds and cool to room temperature in a dark drawer for 30 min).
   NOTE: *HCR hairpins h1 and h2 are provided in hairpin storage buffer ready for snap cooling. h1 and h2 should be snap cooled in separate tubes. This is the amount of hairpins needed for each target on a single slide using 100 $\mu$L of incubation volume.*

5. Prepare hairpin solution by adding all snap-cooled h1 hairpins and snap-cooled h2 hairpins to 100 $\mu$L of amplification buffer at room temperature per section.

6. Remove the pre-amplification solution and drain excess buffer on slide by blotting edges on a Kimwipe.

7. Add 100 $\mu$L of the hairpin solution on top of the tissue sample.

8. Incubate overnight (>12 h) in a dark humidified chamber at room temperature.

9. Remove excess hairpins by immersing slide in 5× SSCT at room temperature for:

   (a) 2 × 5 min
   (b) 2 × 15 min
   (c) 1 × 5 min

**Sample mounting for microscopy**

1. Drain slide by blotting edges on a Kimwipe and dry around the section with another Kimwipe.

2. Apply 35 $\mu$L of Slowfade Diamond antifade mountant with DAPI on top of the tissue.

3. Place a 22 × 30 mm No. 1 coverslip on top carefully to prevent air bubbles.

4. Slides can be stored at 4 °C protected from light prior to imaging.
   NOTE: *see Section S2.5 for details of epifluorescence microscope used to image FFPE mouse brain tissue section.*

##### S3.2.4 Buffer for HCR 1°IHC with/without HCR RNA-ISH

HCR probes (initiator-labeled antibody probes, split-initiator DNA probes), amplifiers, and buffers (antibody buffer, probe hybridization buffer, probe wash buffer, amplification buffer) are available from Molecular Instruments (`www.molecularinstruments.com`). Probe hybridization buffer, and probe wash buffer should be stored at -20 °C. Antibody buffer and amplification buffer should be stored at 4 °C. Make sure all solutions are well mixed before use.

| **5× SSCT** | For 40 mL of solution |
|---|---|
| 5× sodium chloride sodium citrate (SSC) | 10 mL of 20× SSC |
| 0.1% Tween 20 | 400 $\mu$L of 10% Tween 20 |
| | Fill up to 40 mL with ultrapure $H_2O$ |

##### S3.2.5 Reagents and supplies

Pro-Par Clearant (ANATECH LTD Cat. # 510)
100% Ethanol (EtOH) (VWR Cat. # 89125-172)
100× Citrate buffer pH 6.0 (Abcam Cat. #ab93678)
100× Tris-EDTA buffer pH 9.0 (Abcam Cat. #ab93684)
10× Phosphate buffered saline (PBS)
20× sodium chloride sodium citrate (SSC) (Life Technologies Cat. # 15557-044)
10% Tween 20 (Teknova Cat. # T0710)
SlowFade Diamond Antifade Mountant with DAPI (Invitrogen Cat. # S36973)
22 mm × 30 mm No. 1 coverslip (VWR Cat. # 48393-026)

### S4 Protocols for HCR 2°IHC with/without HCR RNA-ISH

#### S4.1 Protocols for mammalian cells on a chambered slide

##### S4.1.1 Preparation of fixed mammalian cells on a chambered slide

1. Coat bottom of each chamber by applying 300 $\mu$L of 0.01% poly-D-lysine prepared in cell culture grade $H_2O$.
   NOTE: *A volume of 300 $\mu$L is sufficient per chamber on an 8-chamber slide. Scale volume accordingly if using a different slide format.*
2. Incubate for at least 30 min at room temperature.
3. Aspirate the coating solution and wash each chamber twice with molecular biology grade $H_2O$.
4. Plate desired number of cells in each chamber.
5. Grow cells to desired confluency for 24–48 h.
6. Aspirate growth media and wash each chamber with 300 $\mu$L of DPBS.
   NOTE: *avoid using calcium chloride and magnesium chloride in DPBS as this leads to increased autofluorescence.*
7. Add 300 $\mu$L of 4% formaldehyde to each chamber.
   CAUTION: *use formaldehyde with extreme care as it is a hazardous material.*
8. Incubate for 10 min at room temperature.
9. Remove fixative and wash each chamber 2 $\times$ 300 $\mu$L of DPBS.
10. Aspirate DPBS and add 300 $\mu$L of ice-cold 70% ethanol (EtOH).
11. Permeabilize cells overnight (or longer) at -20 °C.
12. Proceed to HCR assay.

##### S4.1.2 HCR 2°ICC with/without HCR RNA-ISH using unlabeled primary antibody probes and initiator-labeled secondary antibody probes for protein targets, spit-initiator DNA probes for RNA targets, and simultaneous HCR signal amplification for all targets

**Protein detection stage**

1. Aspirate EtOH from sample and wash samples 2 × 5 min with 300 $\mu$L of 1× PBS.

2. Apply 300 $\mu$L antibody buffer to each chamber. Incubate at room temperature for 1 hr with gentle agitation.

3. Prepare working concentration of primary antibodies in antibody buffer. Prepare 300 $\mu$L per chamber.
   NOTE: *follow manufacturer's guidelines for primary antibody working concentration.*

4. Replace antibody buffer with primary antibody solution and incubate overnight (>12 h) at 4 °C with gentle agitation.
   NOTE: *Incubation may be optimized (e.g., 1–2 h at room temperature) depending on sample type and thickness.*

5. Remove excess antibodies by washing 3 × 5 min with 1× PBST at room temperature with gentle agitation.

6. Prepare working concentration of initiator-labeled secondary antibodies in antibody buffer. Prepare 300 $\mu$L per chamber.
   NOTE: *We recommend starting with a 1 µg/mL working concentration.*

7. Add secondary antibody solution to each chamber and incubate 1 h at room temperature with gentle agitation.

8. Remove excess antibodies by washing 3 × 5 min with 1× PBST at room temperature with gentle agitation.

9. Proceed to **RNA detection stage** for co-detection of protein and RNA. Otherwise, proceed to **Amplification stage**.

**RNA detection stage**

1. Post-fix sample with 300 $\mu$L of 4% formaldehyde.
   CAUTION: *use formaldehyde with extreme care as it is a hazardous material.*

2. Incubate for 10 min at room temperature.

3. Remove fixative and wash each chamber 2 × 300 $\mu$L of PBS.

4. Wash sample with 300 $\mu$L of 2× SSC.

5. Pre-hybridize samples in 300 $\mu$L of probe hybridization buffer for 30 min at 37 °C.
   CAUTION: *Probe hybridization buffer contains formamide, a hazardous material.*
   NOTE: *pre-heat probe hybridization buffer to 37 °C before use.*

6. Prepare a 16 nM probe solution by adding 4.8 pmol of each probe mixture (e.g. 4.8 $\mu$L of 1 $\mu$M stock) to 300 $\mu$L of probe hybridization buffer at 37 °C.
   NOTE: *This is the amount of probe set needed for each target on a single chamber of an 8-well chambered slide using 300 µL of incubation volume.*

7. Remove the pre-hybridization solution and add the probe solution.

8. Incubate samples overnight (>12 h) at 37 °C.

9. Remove excess probes by washing 4 × 5 min with 300 $\mu$L of probe wash buffer at 37 °C.
   CAUTION: *Probe wash buffer contains formamide, a hazardous material.*
   NOTE: *pre-heat probe wash buffer to 37 °C before use.*

10. Wash samples 4 × 5 min with 5× SSCT at room temperature.
11. Wash once with 300 $\mu$L 5× SSCT at room temperature for 5 min.
12. Proceed to **Amplification stage**.

**Amplification stage**

1. Wash once with 300 $\mu$L 5× SSCT at room temperature for 5 min.
2. Pre-amplify samples in 300 $\mu$L of amplification buffer for 30 min at room temperature.
   NOTE: *Equilibrate amplification buffer to room temperature before use.*
3. Separately prepare 18 pmol of hairpin h1 and 18 pmol of hairpin h2 by snap cooling 6 $\mu$L of 3 $\mu$M stock (heat at 95 °C for 90 seconds and cool to room temperature in a dark drawer for 30 min).
   NOTE: *HCR hairpins h1 and h2 are provided in hairpin storage buffer ready for snap cooling. h1 and h2 should be snap cooled in separate tubes. This is the amount of hairpins needed for each target in a single sample using 300 $\mu$L of incubation volume.*
4. Prepare a 60 nM hairpin solution by adding all snap-cooled h1 hairpins and snap-cooled h2 hairpins to 300 $\mu$L of amplification buffer at room temperature per sample.
5. Remove the pre-amplification solution and add the hairpin solution.
6. Incubate the slide overnight (>12 h) protected from light at room temperature.
7. Remove excess hairpins by washing 5 × 5 min with 300 $\mu$L of 5× SSCT at room temperature.

**Sample mounting for microscopy**

1. Remove final wash and add 150 $\mu$L of mounting medium (e.g., Fluoromount-G with DAPI).
2. Slides can be stored at 4 °C protected from light prior to imaging.
   NOTE: *see Section S2.4 for details of confocal microscopes used to image mammalian cells on a chambered slide.*

##### S4.1.3 Buffers for HCR 2°IHC with/without HCR RNA-ISH

HCR probes (initiator-labeled antibody probes, split-initiator DNA probes), amplifiers, and buffers (antibody buffer, probe hybridization buffer, probe wash buffer, amplification buffer) are available from Molecular Instruments (`www.molecularinstruments.com`). Probe hybridization buffer, and probe wash buffer should be stored at -20 °C. Antibody buffer and amplification buffer should be stored at 4 °C. Make sure all solutions are well mixed before use.

| | |
|---|---|
| **1× PBST** | For 40 mL of solution |
| 1× phosphate buffered solution (PBS) | 4 mL of 10× PBS |
| 0.1% Tween 20 | 400 $\mu$L of 10% Tween 20 |
| | Fill up to 40 mL with ultrapure $H_2O$ |

| | |
|---|---|
| **5× SSCT** | For 40 mL of solution |
| 5× sodium chloride sodium citrate (SSC) | 10 mL of 20× SSC |
| 0.1% Tween 20 | 400 $\mu$L of 10% Tween 20 |
| | Fill up to 40 mL with ultrapure $H_2O$ |

##### S4.1.4 Reagents and supplies

ibidi $\mu$-slide ibitreat (ibidi Cat. # 80826)
Poly-D-lysine hydrobromide (Sigma-Aldrich Cat. # P7280)
Molecular biology grade $H_2O$ (Corning Cat. # 46-000-CV)
DPBS, no calcium, no magnesium (Life Technologies Cat. # 14190144)
Image-iT Fixative Solution 4% (Thermo Fisher Scientific Cat. # FB002)
10× PBS (Ambion Cat. # AM9624)
10% Tween 20 (Teknova Cat. # T0710)
20× sodium chloride sodium citrate (SSC) (Life Technologies Cat. # 15557-044)
DAPI Fluoromount-G (SouthernBiotech Cat. # 0100-20)

#### S4.2 Protocols for FFPE mouse brain tissue sections

##### S4.2.1 Preparation of formalin-fixed paraffin-embedded (FFPE) mouse brain tissue sections

1. Bake slides in a dry oven for 1 h at 60 °C to improve sample adhesion to the slide.
2. In a fume hood, deparaffinize FFPE tissue by immersing in Pro-Par Clearant for 2 × 5 min. Move slides up and down occasionally.
   CAUTION: *use Pro-Par Clearant with care as it is a hazardous material.*
   NOTE: *Xylene can be used in place of Pro-Par Clearant.*
   NOTE: *Each 50 mL tube can fit two outward-facing slides. A volume of 30mL is sufficient to immerse sections in the tube. If desired, a larger number of slides can be processed together using a Coplin jar.*
3. Incubate slides in 100% ethanol (EtOH) for 2 × 2 min at room temperature. Move slides up and down occasionally.
4. Rehydrate with a series of graded EtOH washes at room temperature.
   (a) 95% EtOH for 3 min
   (b) 70% EtOH for 3 min
   (c) 50% EtOH for 3 min
   (d) Nanopure water for 3 min
5. Bring 500 mL of 1× citrate buffer (pH 6.0) in a beaker to boil in a microwave.
   NOTE: *1× Tris-EDTA buffer (pH 9.0) can be used in place of citrate buffer (pH 6.0). Optimal antigen retrieval method may differ depending on the antigen/antibody used.*
6. Maintain citrate buffer at 90–95 °C on a hot plate.
7. Immerse slides for 15 min.
   NOTE: *Alternatively, slides may be immersed at 95–99 °C for 15 min in a steamer.*
8. Remove beaker from hot plate and add 100 mL of nanopure water every 5 min to allow temperature to decrease to 45 °C in 20 min.
9. Immerse slides in 400 mL of nanopure water in a separate container for 10 min at room temperature.
10. Immerse slides in 1× PBST for 2 × 2 min at room temperature.
    NOTE: *avoid using calcium chloride and magnesium chloride in PBS as this leads to increased autofluorescence in the tissue.*
11. Drain slide by blotting edges on a Kimwipe.
12. Wipe around the section with a Kimwipe and circle tissue with a hydrophobic pen.
13. Proceed to HCR assay.

##### S4.2.2 Buffer recipes for sample preparation

| **1× citrate buffer** | For 500 mL of solution |
| --- | --- |
| 1× citrate buffer | 5 mL of 100× citrate buffer (pH 6.0) |
| | Fill up to 500 mL with water |

| **1× Tris-EDTA buffer** | For 500 mL of solution |
| --- | --- |
| 1× Tris-EDTA buffer | 5 mL of 100× Tris-EDTA buffer (pH 9.0) |
| | Fill up to 500 mL with water |

| **PBST** | For 50 mL of solution |
| --- | --- |
| 1× PBS | 5 mL of 10× PBS |
| 0.1% Tween 20 | 500 $\mu$L of 10% Tween 20 |
| | Fill up to 50 mL with ultrapure $H_2O$ |

##### S4.2.3 Multiplexed HCR 2°IHC with/without HCR RNA-ISH using unlabeled primary antibody probes and initiator-labeled secondary antibody probes for protein targets, spit-initiator DNA probes for RNA targets, and simultaneous HCR signal amplification for all targets

**Protein detection stage**

1. Block tissue by applying 200 $\mu$L of antibody buffer on top of the sample. Incubate at room temperature for 1 h in a humidified chamber.
2. Prepare working concentration of primary antibodies in antibody buffer. Prepare 100 $\mu$L per section.
   NOTE: *follow manufacturer's guidelines for primary antibody working concentration.*
3. Drain slide by blotting edges on a Kimwipe and wipe around the section with another Kimwipe.
4. Add primary antibody solution to each section and incubate overnight (>12 h) at 4 °C in a humidified chamber.
   NOTE: *Incubation may be optimized (e.g., 1–2 h at room temperature) depending on sample type and thickness.*
5. Remove excess antibodies by immersing slide in 1× PBST at room temperature for 3 × 5 min.
6. Prepare working concentration of initiator-labeled secondary antibodies in antibody buffer. Prepare 100 $\mu$L per section.
7. Drain slide by blotting edges on a Kimwipe and wipe around the section with another Kimwipe.
8. Add secondary antibody solution to each section and incubate for 1 h at room temperature in a humidified chamber.
9. Remove excess antibodies by immersing slide in 1× PBST at room temperature for 3 × 5 min.
10. Proceed to **RNA detection stage** for co-detection of protein and RNA. Otherwise, proceed to **Amplification stage**.

**RNA detection stage**

1. Drain slide by blotting edges on a Kimwipe and wipe around the section with another Kimwipe.
2. Post-fix sample by adding 200 $\mu$L of 4% formaldehyde on the tissue.
   CAUTION: *use formaldehyde with extreme care as it is a hazardous material.*
3. Incubate slides for 10 min at room temperature.
4. Immerse slides for 2 × 5 min in PBST.
5. Immerse slides for 5 min in 5× SSCT.
6. Pre-warm a humidified chamber to 37 °C.
7. Drain slide by blotting edges on a Kimwipe and wipe around the section with another Kimwipe.
8. Add 200 $\mu$L of probe hybridization buffer on top of the tissue sample.
   CAUTION: *Probe hybridization buffer contains formamide, a hazardous material.*
   NOTE: *pre-heat probe hybridization buffer to 37 °C before use.*
9. Pre-hybridize for 10 min inside the humidified chamber.

10. Prepare a 16 nM probe solution by adding 1.6 pmol of each probe set (e.g. 1.6 $\mu$L of 1 $\mu$M stock) to 100 $\mu$L of probe hybridization buffer at 37 °C.
    NOTE: *This is the amount of probe set needed for each target on a single slide using 100 $\mu$L of incubation volume.*

11. Remove the pre-hybridization solution and drain excess buffer on slide by blotting edges on a Kimwipe.

12. Add 100 $\mu$L of the probe solution on top of the tissue sample.

13. Place a coverslip on the sample and incubate overnight ($>$12 h) in the 37 °C humidified chamber.

14. Immerse slide in probe wash buffer at 37 °C to float off coverslip.
    CAUTION: *Probe wash buffer contains formamide, a hazardous material.*

15. Remove excess probes by incubating slide at 37 °C in:

    (a) 75% of probe wash buffer / 25% 5$\times$ SSCT for 15 min
    (b) 50% of probe wash buffer / 50% 5$\times$ SSCT for 15 min
    (c) 25% of probe wash buffer / 75% 5$\times$ SSCT for 15 min
    (d) 100% 5$\times$ SSCT for 15 min

    NOTE: *Wash solutions should be pre-heated to 37 °C before use.*

16. Proceed to **Amplification stage**.

**Amplification stage**

1. Immerse slide in 5× SSCT at room temperature for 5 min.
2. Drain slide by blotting edges on a Kimwipe and wipe around the section with another Kimwipe.
3. Add 200 $\mu$L of amplification buffer on top of the tissue sample and pre-amplify in a humidified chamber for 30 min at room temperature.
   NOTE: *equilibrate amplification buffer to room temperature before use.*
4. Separately prepare 6 pmol of hairpin h1 and 6 pmol of hairpin h2 by snap cooling 2 $\mu$L of 3 $\mu$M stock (heat at 95 °C for 90 seconds and cool to room temperature in a dark drawer for 30 min).
   NOTE: *HCR hairpins h1 and h2 are provided in hairpin storage buffer ready for snap cooling. h1 and h2 should be snap cooled in separate tubes. This is the amount of hairpins needed for each target on a single slide using 100 $\mu$L of incubation volume.*
5. Prepare a 60 nM hairpin solution by adding all snap-cooled h1 hairpins and snap-cooled h2 hairpins to 100 $\mu$L of amplification buffer at room temperature per section.
6. Remove the pre-amplification solution and drain excess buffer on slide by blotting edges on a Kimwipe.
7. Add 100 $\mu$L of the hairpin solution on top of the tissue sample.
8. Incubate overnight (>12 h) in a dark humidified chamber at room temperature.
9. Remove excess hairpins by immersing slide in 5× SSCT at room temperature for:
   (a) 2 × 5 min
   (b) 2 × 15 min
   (c) 1 × 5 min

**Sample mounting for microscopy**

1. Drain slide by blotting edges on a Kimwipe and dry around the section with another Kimwipe.
2. Apply 35 $\mu$L of Slowfade Diamond antifade mountant with DAPI on top of the tissue.
3. Place a 22 × 30 mm No. 1 coverslip on top carefully to prevent air bubbles.
4. Slides can be stored at 4 °C protected from light prior to imaging.
   NOTE: *see Section S2.5 for details of epifluorescence microscope used to image FFPE mouse brain tissue section.*

##### S4.2.4 Buffers for HCR 2°IHC with/without HCR RNA-ISH

HCR probes (initiator-labeled antibody probes, split-initiator DNA probes), amplifiers, and buffers (antibody buffer, probe hybridization buffer, probe wash buffer, amplification buffer) are available from Molecular Instruments (`www.molecularinstruments.com`). Probe hybridization buffer, and probe wash buffer should be stored at -20 °C. Antibody buffer and amplification buffer should be stored at 4 °C. Make sure all solutions are well mixed before use.

| **5× SSCT** | For 40 mL of solution |
|---|---|
| 5× sodium chloride sodium citrate (SSC) | 10 mL of 20× SSC |
| 0.1% Tween 20 | 400 $\mu$L of 10% Tween 20 |
| | Fill up to 40 mL with ultrapure $H_2O$ |

##### S4.2.5 Reagents and supplies

Pro-Par Clearant (ANATECH LTD Cat. # 510)
100% Ethanol (EtOH) (VWR Cat. # 89125-172)
100× Citrate buffer pH 6.0 (Abcam Cat. # ab93678)
100× Tris-EDTA buffer pH 9.0 (Abcam Cat. # ab93684)
10× Phosphate buffered saline (PBS) (Invitrogen Cat. # AM9624)
20× sodium chloride sodium citrate (SSC) (Life Technologies Cat. # 15557-044)
10% Tween 20 (Teknova Cat. # T0710)
SlowFade Diamond Antifade Mountant with DAPI (Invitrogen Cat. # S36973)
22 mm × 30 mm No. 1 coverslip (VWR Cat. # 48393-026)

##### S4.3 Protocols for FFPE human breast tissue sections

###### S4.3.1 Preparation of formalin-fixed paraffin-embedded (FFPE) human breast tissue sections

1. Bake slides in a dry oven for 1 h at 60 °C to improve sample adhesion to the slide.
2. In a fume hood, deparaffinize FFPE tissue by immersing in xylene for 2 × 5 min. Move slides up and down occasionally.
   CAUTION: *use xylene with care as it is a hazardous material.*
   NOTE: *Each 50 mL tube can fit two outward-facing slides. A volume of 30mL is sufficient to immerse sections in the tube. If desired, a larger number of slides can be processed together using a Coplin jar.*
3. Incubate slides in 100% ethanol (EtOH) for 2 × 3 min at room temperature. Move slides up and down occasionally.
4. Rehydrate with a series of graded EtOH washes at room temperature.
   (a) 95% EtOH for 3 min
   (b) 70% EtOH for 3 min
   (c) Ultrapure water for 3 min
5. Remove slides from ultrapure water and gently tap off water.
6. Carefully dry around the tissue with a Kimwipe.
7. Draw a hydrophobic barrier around the tissue with a hydrophobic pen.
8. Apply 200 $\mu$L of 4 U/$\mu$L proteinase K solution for 7 min at room temperature.
   NOTE: *Proteolytic-Induced Epitope Retrieval (PIER) is used in place of Heat-Induced Epitope Retrieval (HIER). Optimal antigen retrieval method may differ depending on the antigen/antibody used.*
9. Gently tap off proteinase K solution and immerse slides in a Coplin jar with ultrapure water for 1 min.
10. Remove slides from ultrapure water and gently tap off water.
11. Carefully dry around the tissue with a Kimwipe.
12. Proceed to HCR assay.

###### S4.3.2 Buffer recipes for sample preparation

| **Proteinase K solution** | For 800 $\mu$L of solution |
|---|---|
| 4 U/$\mu$L proteinase K | 4 $\mu$L of 800 U/$\mu$L proteinase K |
| | 796 $\mu$L of 1× phosphate-buffered saline (PBS) |

##### S4.3.3 Multiplexed HCR 2°IHC using unlabeled primary antibody probes and initiator-labeled secondary probes with simultaneous HCR signal amplification for all targets antibodies

**Protein detection stage**

1. Block tissue by adding 200 $\mu$L of antibody buffer on top of the sample. Incubate at room temperature for 1 h in a humidified chamber.
2. Prepare working concentration of primary antibodies in antibody buffer. Prepare 100 $\mu$L per section.
   NOTE: *follow manufacturer's guidelines for primary antibody working concentration.*
3. Drain slide by blotting edges on a Kimwipe and wipe around the section with another Kimwipe.
4. Add primary antibody solution to each section and incubate overnight (>12 h) at 4 °C in a humidified chamber.
   NOTE: *Incubation may be optimized (e.g., 1–2 h at room temperature) depending on sample type and thickness.*
5. Remove excess antibodies by washing 3 × 5 min with 100 $\mu$L of 1× PBST at room temperature.
6. Prepare working concentration of initiator-labeled secondary antibodies in antibody buffer. Prepare 100 $\mu$L per section.
7. Drain slide by blotting edges on a Kimwipe and wipe around the section with another Kimwipe.
8. Add secondary antibody solution to each section and incubate for 1 h at room temperature in a humidified chamber.
9. Remove excess antibodies by washing 3 × 5 min with 100 $\mu$L of 1× PBST at room temperature.

**Amplification stage**

1. Drain slide by blotting edges on a Kimwipe and wipe around the section with another Kimwipe.
2. Add 200 $\mu$L of amplification buffer on top of the tissue sample and pre-amplify in a humidified chamber for 30 min at room temperature.
   NOTE: *equilibrate amplification buffer to room temperature before use.*
3. Separately prepare 12 pmol of hairpin h1 and 12 pmol of hairpin h2 by snap cooling 4 $\mu$L of 3 $\mu$M stock (heat at 95 °C for 90 seconds and cool to room temperature in a dark drawer for 30 min).
   NOTE: *HCR hairpins h1 and h2 are provided in hairpin storage buffer ready for snap cooling. h1 and h2 should be snap cooled in separate tubes. This is the amount of hairpins needed for each target on a single slide using 200 $\mu$L of incubation volume.*
4. Prepare a 60 nM hairpin solution by adding all snap-cooled h1 hairpins and snap-cooled h2 hairpins to 200 $\mu$L of amplification buffer at room temperature per section.
5. Remove the pre-amplification solution and drain excess buffer on slide by blotting edges on a Kimwipe.
6. Add 200 $\mu$L of the hairpin solution on top of the tissue sample.
7. Incubate overnight (>12 h) in a dark humidified chamber at room temperature.
8. Remove excess hairpins by washing with 100 $\mu$L 5× SSCT at room temperature:
   (a) 2 × 5 min
   (b) 2 × 15 min
   (c) 1 × 5 min

**Sample mounting for microscopy**

1. Drain slide by blotting edges on a Kimwipe and dry around the section with another Kimwipe.
2. Apply 20 $\mu$L of Fluoromount-G with DAPI on top of the tissue.
3. Place a 22 $\times$ 30 mm No. 1 coverslip on top carefully to prevent air bubbles.
4. Seal the edges of the coverslip by applying nail polish hardener and allow it to dry for 30 min.
5. Slides can be stored at 4 °C protected from light prior to imaging.
   NOTE: *see Section S2.4 for details of confocal microscopes used to image FFPE human breast tissue section.*

##### S4.3.4 Buffers for HCR 2°IHC

HCR probes (initiator-labeled antibody probes), amplifiers, and buffers (antibody buffer, amplification buffer) are available from Molecular Instruments (`www.molecularinstruments.com`). Antibody buffer and amplification buffer should be stored at 4 °C. Make sure all solutions are well mixed before use.

| **PBST** | For 50 mL of solution |
| --- | --- |
| 1× PBS | 5 mL of 10× PBS |
| 0.1% Tween 20 | 500 $\mu$L of 10% Tween 20 |
| | Fill up to 50 mL with ultrapure $H_2O$ |

| **5× SSCT** | For 40 mL of solution |
| --- | --- |
| 5× sodium chloride sodium citrate (SSC) | 10 mL of 20× SSC |
| 0.1% Tween 20 | 400 $\mu$L of 10% Tween 20 |
| | Fill up to 40 mL with ultrapure $H_2O$ |

##### S4.3.5 Reagents and supplies

Pro-Par Clearant (ANATECH LTD Cat. # 510)
100% Ethanol (EtOH) (VWR Cat. # 89125-172)
Proteinase K, molecular biology grade (NEB Cat. # P8107S)
10× Phosphate buffered saline (PBS) (Invitrogen Cat. # AM9624)
20× sodium chloride sodium citrate (SSC) (Life Technologies Cat. # 15557-044)
10% Tween 20 (Teknova Cat. # T0710)
Fluoromount-G with DAPI (SouthernBiotech Cat. # 0100-20)
22 mm × 30 mm No. 1 coverslip (VWR Cat. # 48393-026)

#### S4.4 Protocols for whole-mount zebrafish embryos

This protocol has been optimized for embryos at 27 hpf. Other developmental stages may require additional optimization.

##### S4.4.1 Preparation of whole-mount zebrafish embryos

1. Collect zebrafish embryos and incubate at 28 °C in a petri dish with egg $H_2O$.
2. Dechorionate embryos at 27 hpf and wash with fresh egg $H_2O$.
3. Transfer 40 embryos to a 2 mL eppendorf tube and remove excess egg $H_2O$.
4. Fix embryos in 2 mL of 4% paraformaldehyde (PFA) for 24 h at 4 °C.
   CAUTION: *use PFA with extreme care as it is a hazardous material.*
   NOTE: *use fresh PFA and cool to 4 °C before use to avoid increased autofluorescence.*
5. Wash embryos 3 × 5 min with 1 mL of 1× phosphate-buffered saline (PBS) to stop the fixation.
   NOTE: *avoid using calcium chloride and magnesium chloride in PBS as this leads to increased autofluorescence in the samples.*
6. Dehydrate and permeabilize with a series of methanol (MeOH) washes at room temperature (1 mL each):
   (a) 100% MeOH for 4 × 10 min
   (b) 100% MeOH for 1 × 50 min
7. Store embryos at -20 °C overnight before use.
   NOTE: *Embryos can be stored for six months at -20 °C.*
8. Rehydrate with a series of graded MeOH/PBST washes for 5 min each at room temperature (1 mL each):
   (a) 75% MeOH / 25% PBST
   (b) 50% MeOH / 50% PBST
   (c) 25% MeOH / 75% PBST
   (d) 5 × 100% PBST

##### S4.4.2 Buffer recipes for sample preparation

| **4% Paraformaldehyde (PFA)** | For 25 mL of solution |
|---|---|
| 4% PFA | 1 g of PFA powder |
| 1× PBS | 25 mL of 1× PBS |
| | Heat to 50–60 °C to dissolve powder |

| **PBST** | For 50 mL of solution |
|---|---|
| 1× PBS | 5 mL of 10× PBS |
| 0.1% Tween 20 | 500 $\mu$L of 10% Tween 20 |
| | Fill up to 50 mL with ultrapure $H_2O$ |

##### S4.4.3 Multiplexed HCR 2°IHC using unlabeled primary antibody probes and initiator-labeled secondary antibody probes with simultaneous HCR signal amplification for all targets

**Protein detection stage**

1. Block embryos with 500 $\mu$L of zebrafish blocking buffer for 4 h at 4 °C.
2. Transfer 8 embryos to a 1.5 mL Eppendorf tube for each sample.
3. Prepare working concentration of unlabeled primary antibodies in zebrafish antibody buffer. Prepare 250 $\mu$L per sample.
   NOTE: *follow manufacturer's guidelines for primary antibody working concentration.*
4. Remove zebrafish blocking buffer and add primary antibody solution to embryos.
5. Incubate embryos overnight (>12 h) at 4 °C with gentle rotation (50 RPM).
6. Remove excess antibodies by washing 4 × 30 min with 500 $\mu$L of PBT at room temperature.
7. Prepare working concentration of initiator-labeled secondary antibodies in zebrafish antibody buffer. Prepare 250 $\mu$L per sample.
8. Remove PBST and add secondary antibody solution to embryos.
9. Incubate embryos for 3 h at room temperature with gentle rotation (50 RPM).
10. Remove excess antibodies by washing 5 × 5 min with 500 $\mu$L of PBT at room temperature.
11. Wash 1 × 5 min with 500 $\mu$L of 5× SSCT at room temperature.

**Amplification stage**

1. Pre-amplify embryos with 350 $\mu$L of amplification buffer for 30 min at room temperature.
   NOTE: *equilibrate amplification buffer to room temperature before use.*
2. Separately prepare 30 pmol of hairpin h1 and 30 pmol of hairpin h2 by snap cooling 10 $\mu$L of 3 $\mu$M stock (heat at 95 °C for 90 seconds and cool to room temperature in a dark drawer for 30 min).
   NOTE: *HCR hairpins h1 and h2 are provided in hairpin storage buffer ready for snap cooling. h1 and h2 should be snap cooled in separate tubes. This is the amount of hairpins needed for each target on a single slide using 500 $\mu$L of incubation volume.*
3. Prepare a 60 nM hairpin solution by adding all snap-cooled h1 hairpins and snap-cooled h2 hairpins to 500 $\mu$L of amplification buffer at room temperature per sample.
4. Remove the pre-amplification solution and add the hairpin solution.
5. Incubate the samples overnight (>12 h) in the dark at room temperature.
6. Remove excess hairpins by washing with 500 $\mu$L of 5× SSCT at room temperature:
   (a) 2 × 5 min
   (b) 2 × 30 min
   (c) 1 × 5 min

###### S4.4.4 Sample mounting for microscopy

1. Make a chamber for mounting the embryos by aligning two stacks of Scotch tape (6 pieces per stack) 2 cm apart on a 25 mm $\times$ 75 mm SuperFrost Plus glass slide.
2. Pipet embryos onto glass slide with a cut P1000 pipet tip. Use a P200 pipet to remove excess 5$\times$ SSCT.
3. Gently add 50 $\mu$L Fluoromount-G onto the embryos.
4. Use an eyelash tool to gently position the embryos onto their side for lateral imaging.
5. Use fine forceps to gradually lower a 22 $\times$ 30 mm No. 1 coverslip onto the tape stacks.
6. Gently add Fluoromount-G via the open sides of the chamber until the chamber is full (approximately 100 $\mu$L).
7. Seal the edges of the coverslip by applying nail polish hardener.
8. Let nail polish hardener dry for 30 min.
9. Slides can be stored at 4 °C protected from light prior to imaging.
   NOTE: *see Section S2.4 for details of confocal microscopes used to image whole-mount zebrafish embryos.*

##### S4.4.5 Buffers for HCR 2°IHC

HCR probes (initiator-labeled antibody probes), amplifiers, and buffers (zebrafish blocking buffer, zebrafish antibody buffer, amplification buffer) are available from Molecular Instruments (`www.molecularinstruments.com`). Zebrafish blocking buffer, zebrafish antibody buffer, and amplification buffer should be stored at 4 °C. Make sure all solutions are well mixed before use.

| **0.1 M phosphate buffer with Triton X-100 (PBT)** | For 40 mL of solution |
|---|---|
| 0.8% Triton X-100 | 320 $\mu$L of Triton X-100 |
| 0.061 M $K_2HPO_4$ | 0.555 g $K_2HPO_4$ $3H_2O$ |
| 0.039 M $KH_2PO_4$ | 0.214 g $KH_2PO_4$ |
| | Fill up to 40 mL with ultrapure $H_2O$ |

| **5× SSCT** | For 40 mL of solution |
|---|---|
| 5× sodium chloride sodium citrate (SSC) | 10 mL of 20× SSC |
| 0.1% Tween 20 | 400 $\mu$L of 10% Tween 20 |
| | Fill up to 40 mL with ultrapure $H_2O$ |

##### S4.4.6 Reagents and supplies

Paraformaldehyde (Sigma Cat. # P6148)
10× Phosphate buffered saline (PBS) (Invitrogen Cat. # AM9624)
Methanol (MeOH) (Mallinckrodt Chemicals Cat. # 3016-16)
$K_2HPO_4$ $3H_2O$ (Sigma Cat. # P9666)
$KH_2PO_4$ (Sigma Cat. # P9791)
Triton X-100 (Sigma Cat. # X100)
20× sodium chloride sodium citrate (SSC) (Life Technologies Cat. # 15557-044)
10% Tween 20 (Teknova Cat. # T0710)
Fluoromount-G (SouthernBiotech Cat. # 0100-01)
25 mm × 75 mm SuperFrost Plus glass slide (VWR Cat. # 48311-703)
22 mm × 30 mm No. 1 coverslip (VWR Cat. # 48393-026)

### S5 Additional studies

#### S5.1 Summary of signal-to-background estimates for HCR 1°IHC, HCR 2°IHC, and/or HCR RNA-ISH

| Method | Sample | Microscopy | Target | Type | Probes | Amplifier | SIG/BACK | Plex | Figures | Table |
|---|---|---|---|---|---|---|---|---|---|---|
| HCR 1°IHC | mammalian cells on a slide | Confocal | HSP60 | protein | 1°mAb-init | B3-Alexa488 | 609 ± 18 | 3 | 2C, S1 | S12 |
| HCR 1°IHC | mammalian cells on a slide | Confocal | GM130 | protein | 1°mAb-init | B2-Alexa647 | 211 ± 15 | 3 | 2C, S1 | S12 |
| HCR 1°IHC | mammalian cells on a slide | Confocal | SC35 | protein | 1°mAb-init | B4-Alexa546 | 41 ± 2 | 3 | 2C, S1 | S12 |
| HCR 1°IHC | FFPE mouse brain section | Epifluorescence | TH | protein | 1°mAb-init | B1-Alexa488 | 30 ± 5 | 4 | 2DE, S5 | S13 |
| HCR 1°IHC | FFPE mouse brain section | Epifluorescence | GFAP | protein | 1°mAb-init | B3-Alexa546 | 290 ± 60 | 4 | 2DE, S5 | S13 |
| HCR 1°IHC | FFPE mouse brain section | Epifluorescence | MBP | protein | 1°mAb-init | B5-Alexa647 | 58 ± 14 | 4 | 2DE, S5 | S13 |
| HCR 1°IHC | FFPE mouse brain section | Epifluorescence | MAP2 | protein | 1°mAb-init | B4-Alexa750 | 30 ± 8 | 4 | 2DE, S5 | S13 |
| HCR 2°IHC | mammalian cells on a slide | Confocal | PCNA | protein | 1°mAb + 2°pAb-init | B5-Alexa647 | 87 ± 6 | 3 | 3C, S10 | S14 |
| HCR 2°IHC | mammalian cells on a slide | Confocal | HSP60 | protein | 1°mAb + 2°pAb-init | B3-Alexa546 | 106 ± 7 | 3 | 3C, S10 | S14 |
| HCR 2°IHC | mammalian cells on a slide | Confocal | SC35 | protein | 1°mAb + 2°pAb-init | B2-Alexa488 | 69 ± 6 | 3 | 3C, S10 | S14 |
| HCR 2°IHC | FFPE mouse brain section | Epifluorescence | TH | protein | 1°pAb + 2°pAb-init | B4-Alexa488 | 72 ± 14 | 4 | 3DE, S14 | S15 |
| HCR 2°IHC | FFPE mouse brain section | Epifluorescence | GFAP | protein | 1°pAb + 2°pAb-init | B1-Alexa546 | 23 ± 11 | 4 | 3DE, S14 | S15 |
| HCR 2°IHC | FFPE mouse brain section | Epifluorescence | PVALB | protein | 1°mAb + 2°pAb-init | B5-Alexa647 | 170 ± 60 | 4 | 3DE, S14 | S15 |
| HCR 2°IHC | FFPE mouse brain section | Epifluorescence | MBP | protein | 1°mAb + 2°pAb-init | B3-Alexa750 | 260 ± 40 | 4 | 3DE, S14 | S15 |
| HCR 2°IHC | whole-mount zebrafish embryo | Confocal | Elavl3/Elavl4 | protein | 1°mAb + 2°pAb-init | B1-Alexa647 | 20 ± 4 | 1 | S19 | S16 |
| HCR 1°IHC | FFPE mouse brain section | Epifluorescence | TH | protein | 1°mAb-init | B1-Alexa647 | 45 ± 9 | 2 | 4B, S29 | S22 |
| HCR 1°IHC | FFPE mouse brain section | Epifluorescence | TH | protein | 1°mAb-init | B3-Alexa750 | 41 ± 9 | 2 | 4B, S29 | S22 |
| HCR 2°IHC | FFPE human breast section | Confocal | Krt17 | protein | 1°pAb + 2°pAb-init | B4-Alexa546 | 100 ± 30 | 2 | 4B, S30, S32 | S24 |
| HCR 2°IHC | FFPE human breast section | Confocal | Krt17 | protein | 1°pAb + 2°pAb-init | B3-Alexa647 | 110 ± 30 | 2 | 4B, S30, S32 | S24 |
| HCR 2°IHC | FFPE human breast section | Confocal | Krt19 | protein | 1°mAb + 2°pAb-init | B2-Alexa546 | 140 ± 10 | 2 | S31, S32 | S24 |
| HCR 2°IHC | FFPE human breast section | Confocal | Krt19 | protein | 1°mAb + 2°pAb-init | B5-Alexa647 | 170 ± 10 | 2 | S31, S32 | S24 |
| HCR 1°IHC + HCR RNA-ISH | mammalian cells on a slide | Confocal | PCNA | protein | 1°mAb-init | B5-Alexa488 | 95 ± 6 | 4 | 5B, S33 | S25 |
| HCR 1°IHC + HCR RNA-ISH | mammalian cells on a slide | Confocal | HSP60 | protein | 1°mAb-init | B3-Alexa546 | 280 ± 10 | 4 | 5B, S33 | S25 |
| HCR 1°IHC + HCR RNA-ISH | mammalian cells on a slide | Confocal | *U6* | RNA | 2 split-initiator pairs | B1-Alexa594 | 20.3 ± 1.3 | 4 | 5B, S33 | S25 |
| HCR 1°IHC + HCR RNA-ISH | mammalian cells on a slide | Confocal | *ACTB* | mRNA | 10 split-initiator pairs | B2-Alexa647 | 107 ± 12 | 4 | 5B, S33 | S25 |
| HCR 1°IHC + HCR RNA-ISH | FFPE mouse brain section | Epifluorescence | TH | protein | 1°mAb-init | B1-Alexa488 | 16.2 ± 0.9 | 4 | 5CD, S35 | S26 |
| HCR 1°IHC + HCR RNA-ISH | FFPE mouse brain section | Epifluorescence | MBP | protein | 1°mAb-init | B5-Alexa546 | 33 ± 5 | 4 | 5CD, S35 | S26 |
| HCR 1°IHC + HCR RNA-ISH | FFPE mouse brain section | Epifluorescence | *Prkcd* | mRNA | 31 split-initiator pairs | B2-Alexa647 | 14 ± 3 | 4 | 5CD, S35 | S26 |
| HCR 1°IHC + HCR RNA-ISH | FFPE mouse brain section | Epifluorescence | *Slc17a7* | mRNA | 36 split-initiator pairs | B4-Alexa750 | 41 ± 14 | 4 | 5CD, S35 | S26 |
| HCR 2°IHC + HCR RNA-ISH | mammalian cells on a slide | Confocal | PCNA | protein | 1°mAb + 2°pAb-init | B5-Alexa488 | 27 ± 2 | 4 | 6B, S37 | S27 |
| HCR 2°IHC + HCR RNA-ISH | mammalian cells on a slide | Confocal | HSP60 | protein | 1°mAb + 2°pAb-init | B4-Alexa546 | 240 ± 20 | 4 | 6B, S37 | S27 |
| HCR 2°IHC + HCR RNA-ISH | mammalian cells on a slide | Confocal | *U6* | RNA | 2 split-initiator pairs | B1-Alexa594 | 279 ± 17 | 4 | 6B, S37 | S27 |
| HCR 2°IHC + HCR RNA-ISH | mammalian cells on a slide | Confocal | *HSP60* | mRNA | 18 split-initiator pairs | B2-Alexa647 | 45 ± 6 | 4 | 6B , S37 | S27 |
| HCR 2°IHC + HCR RNA-ISH | FFPE mouse brain section | Epifluorescence | TH | protein | 1°mAb + 2°pAb-init | B4-Alexa488 | 65 ± 17 | 4 | 6CD, S39 | S28 |
| HCR 2°IHC + HCR RNA-ISH | FFPE mouse brain section | Epifluorescence | MBP | protein | 1°mAb + 2°pAb-init | B3-Alexa546 | 40 ± 16 | 4 | 6CD, S39 | S28 |
| HCR 2°IHC + HCR RNA-ISH | FFPE mouse brain section | Epifluorescence | *Prkcd* | mRNA | 31 split-initiator pairs | B1-Alexa647 | 22 ± 8 | 4 | 6CD, S39 | S28 |
| HCR 2°IHC + HCR RNA-ISH | FFPE mouse brain section | Epifluorescence | *Slc17a7* | mRNA | 36 split-initiator pairs | B2-Alexa750 | 50 ± 30 | 4 | 6CD, S39 | S28 |

**Table S9. Signal-to-background summary for protein imaging using HCR 1°IHC or HCR 2°IHC and for simultaneous protein and RNA imaging using HCR 1°IHC + HCR RNA-ISH or HCR 2°IHC + HCR RNA-ISH.** Mean ± standard error of the mean. For mammalian cells on a slide, estimates are based on $N = 15$

**Table S9 continued.** representative rectangular regions (one rectangle in each of 5 individual cells in each of 3 replicate wells on a multi-well slide). For FFPE mouse brain sections and FFPE human breast sections, estimates are based on representative rectangular regions of $N = 3$ replicate sections. For whole-mount zebrafish embryos, estimates are based on representative rectangular regions of $N = 3$ replicate embryos.

| | HCR 1°IHC | HCR 2°IHC | Overall |
|---|---|---|---|
| Mammalian cells on a slide | $210 \pm 170$ | $87 \pm 18$ | $100 \pm 40$ |
| FFPE mouse brain section | $43 \pm 13$ | $120 \pm 70$ | $50 \pm 20$ |
| FFPE human breast section | — | $130 \pm 20$ | $130 \pm 20$ |
| Whole-mount zebrafish embryo | — | 20 | 20 |
| Overall | $45 \pm 15$ | $100 \pm 40$ | $90 \pm 50$ |

**Table S10. Signal-to-background summary for protein imaging using HCR 1°IHC or HCR 2°IHC in mammalian cells on a slide, FFPE mouse brain sections, FFPE human breast sections, and whole-mount zebrafish embryos.** Median $\pm$ median absolute deviation. The number of imaging scenarios for each combination of sample and method is as follows: (mammalian cells on a slide, HCR 1°IHC, $N = 3$), (FFPE mouse brain section, HCR 1°IHC, $N = 6$), (mammalian cells on a slide, HCR 2°IHC, $N = 3$), (FFPE mouse brain section, HCR 2°IHC, $N = 4$), (FFPE human breast section, HCR 2°IHC, $N = 4$), (whole-mount zebrafish embryo, HCR 2°IHC, $N = 1$). The total number of imaging scenarios is $N = 21$. See Table S9 for details.

| | Target proteins | Target RNAs | Overall |
|---|---|---|---|
| Mammalian cells on a slide | $170 \pm 90$ | $80 \pm 40$ | $100 \pm 80$ |
| FFPE mouse brain section | $37 \pm 12$ | $32 \pm 14$ | $37 \pm 14$ |
| Overall | $50 \pm 30$ | $40 \pm 20$ | $40 \pm 20$ |

**Table S11. Signal-to-background summary for simultaneous protein and RNA imaging using HCR 1°IHC + HCR RNA-ISH or HCR 2°IHC + HCR RNA-ISH in mammalian cells on a slide or FFPE mouse brain sections.** Median $\pm$ median absolute deviation. $N = 4$ imaging scenarios for each combination of sample and target types (two targets for each of two methods). The total number of imaging scenarios is $N = 16$. See Table S9 for details.

#### S5.2 Replicates, signal, background, background components, and noise for multiplexed HCR 1°IHC (cf. Figure 2)

##### S5.2.1 Mammalian cells on a slide

For 3-plex protein imaging using HCR 1°IHC in mammalian cells on a slide, the 4 channels are (3 proteins + DAPI):

- **Ch1:** Target protein HSP60, probe 1°mAb rabbit IgG anti-HSP60 labeled with B3 initiator, amplifier B3-Alexa488.
- **Ch2:** Target protein GM130, probe 1°mAb rabbit IgG anti-GM130 labeled with B2 initiator, amplifier B2-Alexa647.
- **Ch3:** Target protein SC35, probe 1°mAb mouse IgG1 anti-SC35 labeled with B4 initiator, amplifier B4-Alexa546.
- **Ch4:** DAPI.

Additional studies are presented as follows:

- Figure S1 displays 3-plex images for $N = 3$ replicate wells on a multi-well slide (cf. Figure 2C).
- Figures S2–S4 displays representative regions of individual channels used for measurement of signal and background for each target.
- Table S12 displays estimated values for signal, background, and signal-to-background for each target.

**Protocol:** HCR 1°IHC (Section S3.1) using initiator-labeled primary antibody probes with HCR signal amplification for all targets simultaneously.
**Sample:** HeLa cells.
**Microscopy:** Confocal.

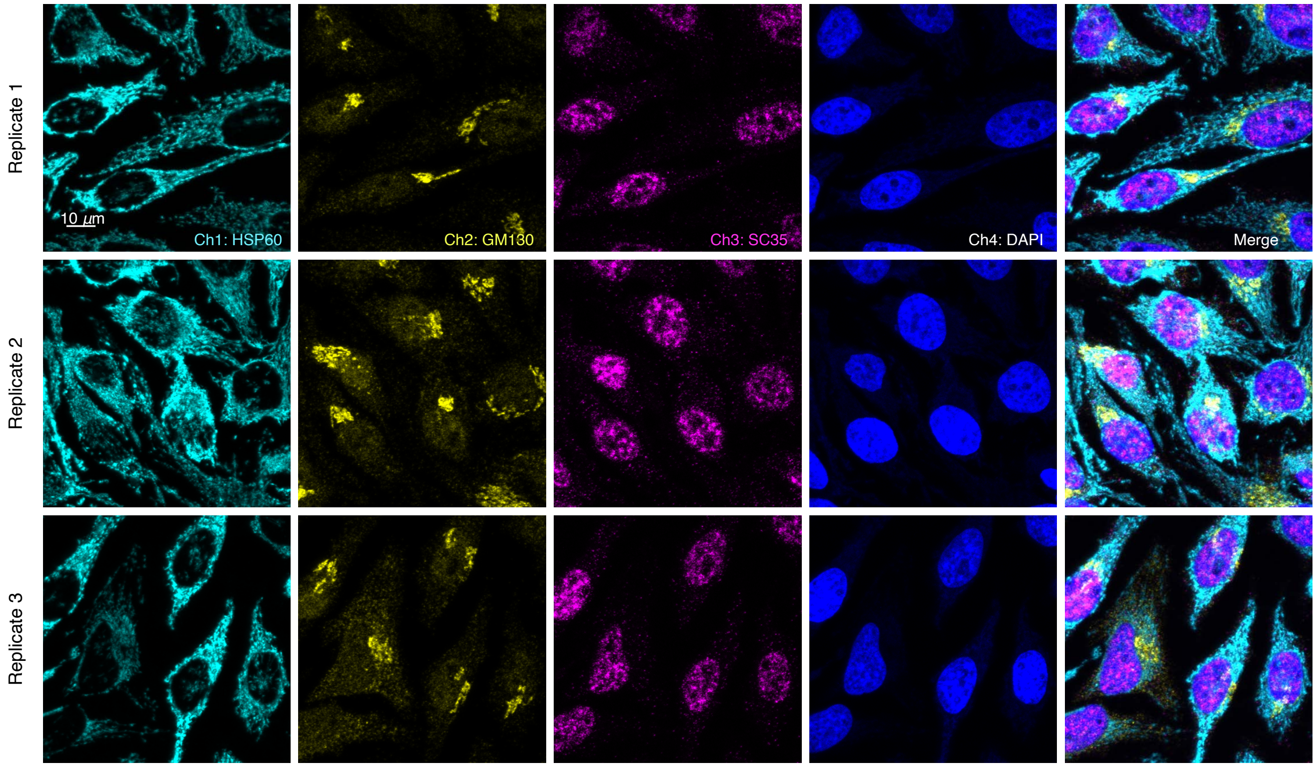

**Figure S1. Replicates for 3-plex protein imaging using HCR 1°IHC in mammalian cells on a slide (cf. Figures 2C).** 4-channel confocal images for 3 replicate wells on a multi-well slide; maximum intensity z-projection. Ch1: target protein HSP60 (Alexa488). Ch2: target protein GM130 (Alexa647). Ch3: target protein SC35 (Alexa546). Ch4: DAPI. Sample: HeLa cells.

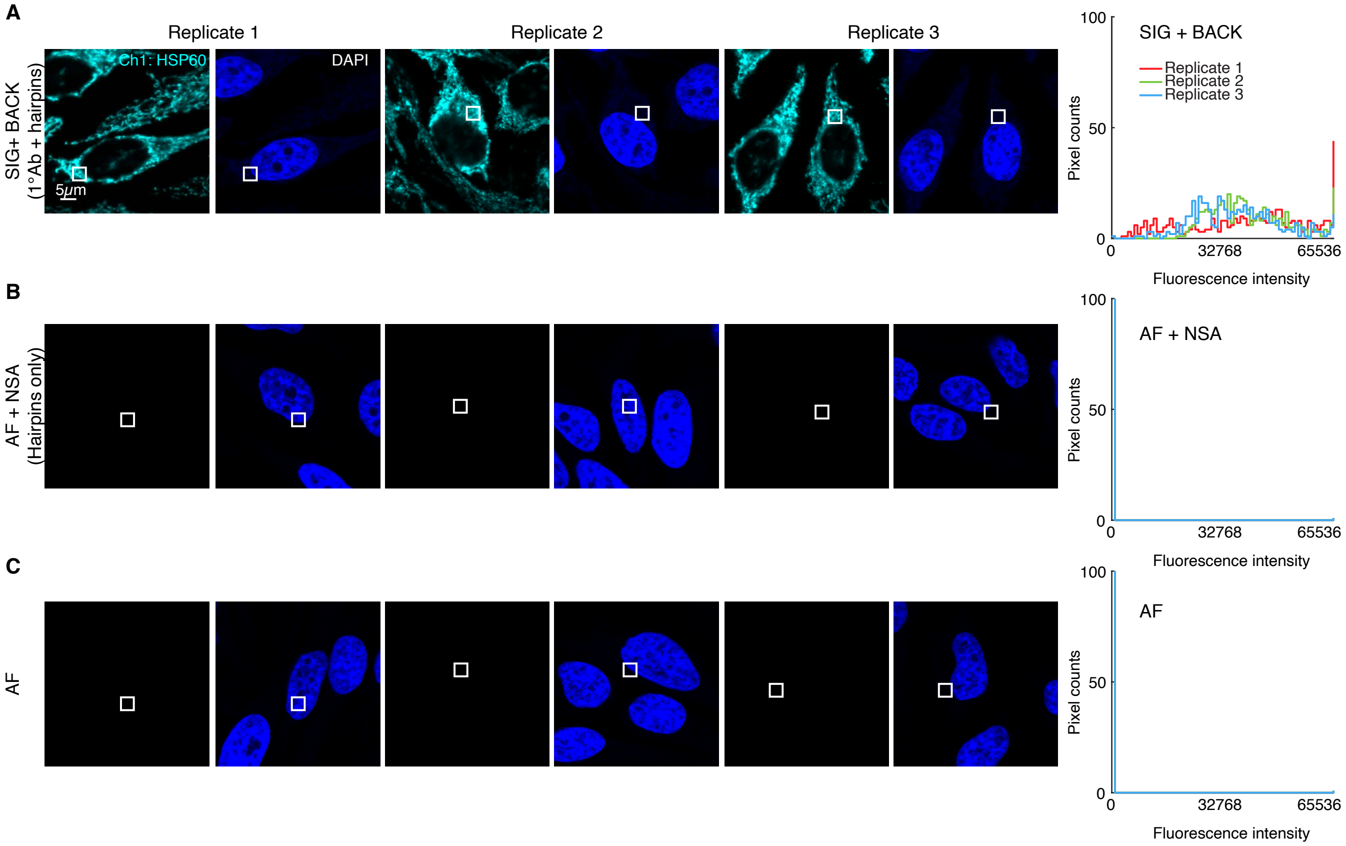

**Figure S2. Measurement of signal, background, and background components for target protein HSP60 using HCR 1°IHC in mammalian cells on a slide (cf. Figure 2C).** (A) Use experiment of Type 1 in Table S7A (1°Ab probe + hairpins) to measure SIG+BACK in a region of high expression. (B) Use experiment of Type 2 in Table S7B (no probes, hairpins only) to measure NSA+AF in a region of maximum background. (C) Use experiment of Type 3 in Table S7B (no probes, no hairpins) to measure AF in a region of maximum background. Left: confocal image collected with the microscope gain optimized to avoid saturating SIG+BACK pixels; DAPI channel facilitates placement of rectangles; single optical section. Right: pixel intensity histograms for representative regions (one rectangle in each of 5 individual cells in each of 3 replicate wells on a multi-well slide). Ch1: target protein HSP60 (Alexa488). Ch4: DAPI. Sample: HeLa cells.

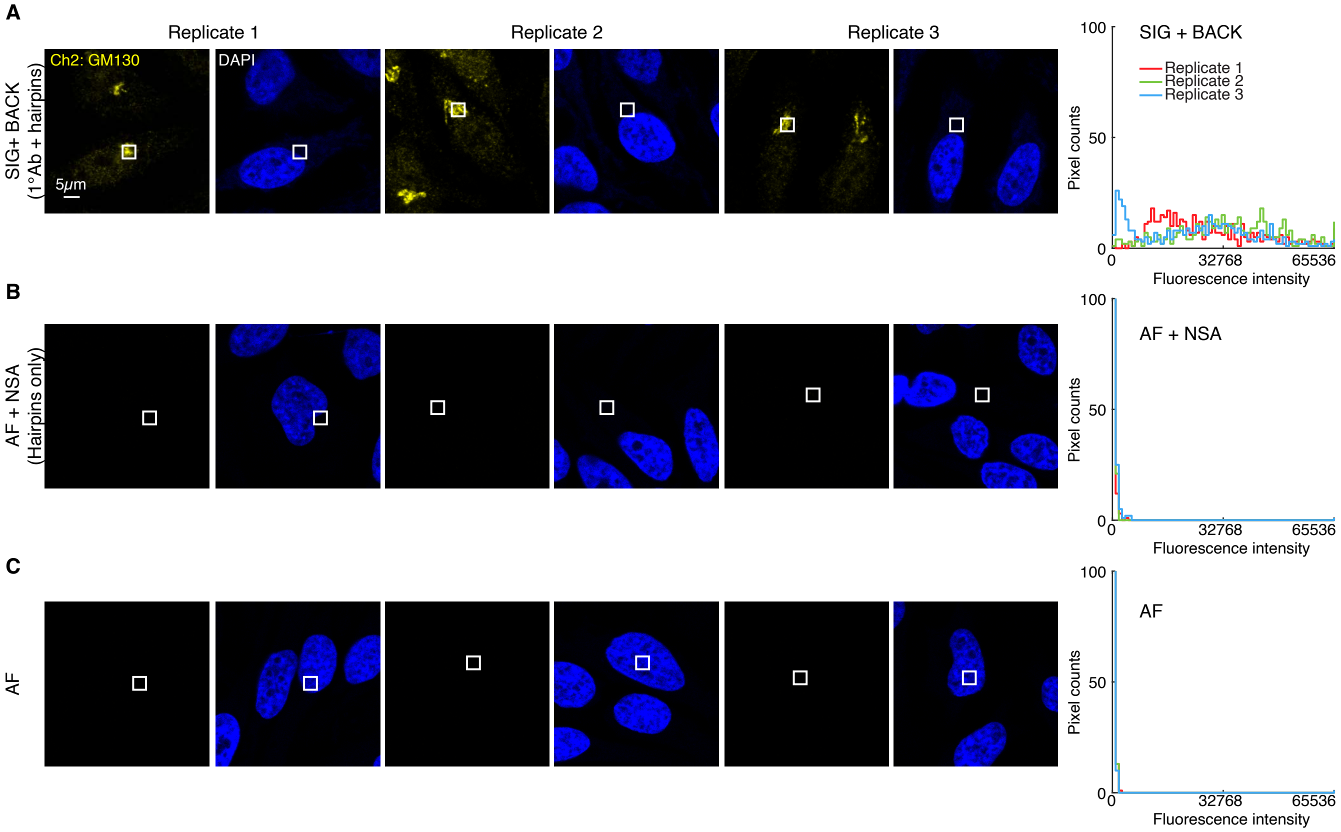

**Figure S3. Measurement of signal, background, and background components for target protein GM130 using HCR 1°IHC in mammalian cells on a slide (cf. Figure 2C).** (A) Use experiment of Type 1 in Table S7A (1°Ab probe + hairpins) to measure SIG+BACK in a region of high expression. (B) Use experiment of Type 2 in Table S7B (no probes, hairpins only) to measure NSA+AF in a region of maximum background. (C) Use experiment of Type 3 in Table S7B (no probes, no hairpins) to measure AF in a region of maximum background. Left: confocal image collected with the microscope gain optimized to avoid saturating SIG+BACK pixels; DAPI channel facilitates placement of rectangles; single optical section. Right: pixel intensity histograms for representative regions (one rectangle in each of 5 individual cells in each of 3 replicate wells on a multi-well slide). Ch2: target protein GM130 (Alexa 647). Ch4: DAPI. Sample: HeLa cells.

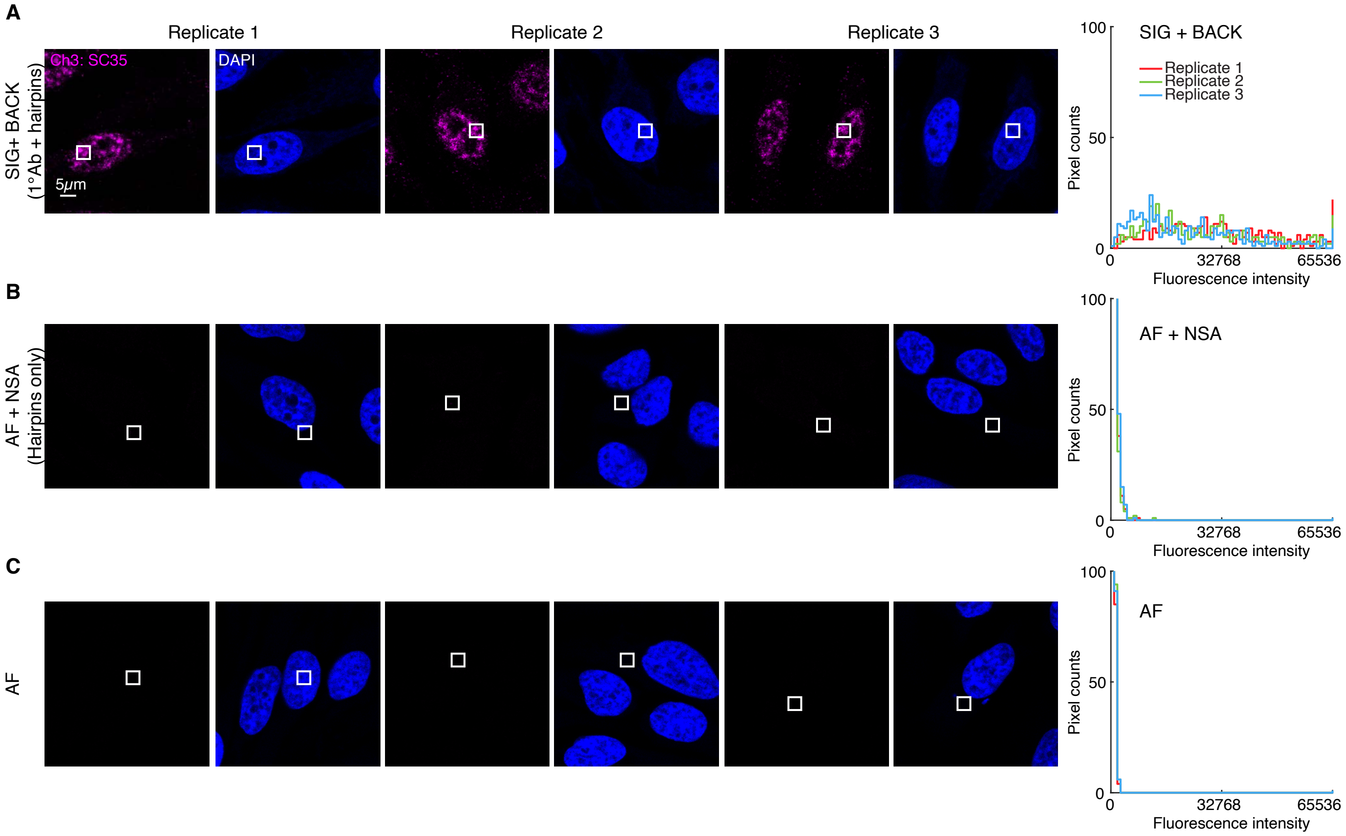

**Figure S4. Measurement of signal, background, and background components for target protein SC35 using HCR 1°IHC in mammalian cells on a slide (cf. Figure 2C).** (A) Use experiment of Type 1 in Table S7A (1°Ab probe + hairpins) to measure SIG+BACK in a region of high expression. (B) Use experiment of Type 2 in Table S7B (no probes, hairpins only) to measure NSA+AF in a region of maximum background. (C) Use experiment of Type 3 in Table S7B (no probes, no hairpins) to measure AF in a region of maximum background. Left: confocal image collected with the microscope gain optimized to avoid saturating SIG+BACK pixels; DAPI channel facilitates placement of rectangles; single optical section. Right: pixel intensity histograms for representative regions (one rectangle in each of 5 individual cells in each of 3 replicate wells on a multi-well slide). Ch3: target protein SC35 (Alexa546). Ch4: DAPI. Sample: HeLa cells.

| | Quantity | Ch1: HSP60<br>B3-Alexa488 | Ch2: GM130<br>B2-Alexa647 | Ch3: SC35<br>B4-Alexa546 | Reagents<br>1°Ab-init | <br>Hairpins | Figure<br>panel |
|---|---|---|---|---|---|---|---|
| **A** | SIG+BACK | 39 400 ± 1100 | 29 400 ± 1500 | 27 300 ± 900 | ✓ | ✓ | A |
| | SIG | 39 400 ± 1100 | 29 300 ± 1500 | 26 700 ± 900 | | | |
| | SIG/BACK | 609 ± 18 | 211 ± 15 | 41 ± 2 | | | |
| **B** | NSA+AF | 64.6 ± 0.4 | 139 ± 6 | 650 ± 30 | | ✓ | B |
| | AF | 66.3 ± 0.7 | 109.5 ± 1.8 | 257 ± 5 | | | C |
| | NSA | $< 0.8$ | 29 ± 7 | 390 ± 30 | | | |

**Table S12. Estimated signal-to-background and background components for 3-plex protein imaging using HCR 1°IHC in mammalian cells on a slide (cf. Figure 2C).** (A) Estimated signal-to-background (SIG/BACK) based on methods of Section S2.6.2. The signal estimate SIG is calculated using the background approximation BACK $\approx$ NSA+AF. (B) Estimated background components (AF, NSA) based on methods of Section S2.6.3. Instrument noise is negligible using confocal microscopy so calculations use the approximation NOISE $\approx 0$. Mean $\pm$ standard error of the mean, $N = 15$ representative rectangular regions (one rectangle in each of 5 individual cells in each of 3 replicate wells on a multi-well slide). Analysis based on representative rectangular regions (examples depicted in Figures S2–S4).

##### S5.2.2 FFPE mouse brain sections

For 4-plex protein imaging using HCR 1°IHC in FFPE mouse brain sections, the 5 channels are (4 proteins + DAPI):

- **Ch1:** Target protein TH, probe 1°mAb rabbit IgG anti-TH labeled with B1 initiator, amplifier B1-Alexa488.
- **Ch2:** Target protein GFAP, probe 1°mAb rabbit IgG anti-GFAP labeled with B3 initiator, amplifier B3-Alexa546.
- **Ch3:** Target protein MBP probe 1°mAb rabbit IgG anti-MBP labeled with B5 initiator, amplifier B5-Alexa647.
- **Ch4:** Target protein MAP2, probe 1°mAb rabbit IgG anti-MAP2 labeled with B4 initiator, amplifier B4-Alexa750.
- **Ch5:** DAPI.

Additional studies are presented as follows:

- Figure S5 displays 4-plex images for $N = 3$ replicate FFPE mouse brain sections (cf. Figure 2C).
- Figures S6–S9 display representative regions of individual channels used for measurement of signal and background for each target.
- Table S13 displays estimated values for signal, background, and signal-to-background for each target.

**Protocol:** HCR 1°IHC (Section S3.2) using initiator-labeled primary antibody probes with HCR signal amplification for all targets simultaneously.
**Sample:** FFPE C57BL/6 mouse brain section (coronal); thickness: 5 $\mu$m.
**Microscopy:** Epifluorescence.

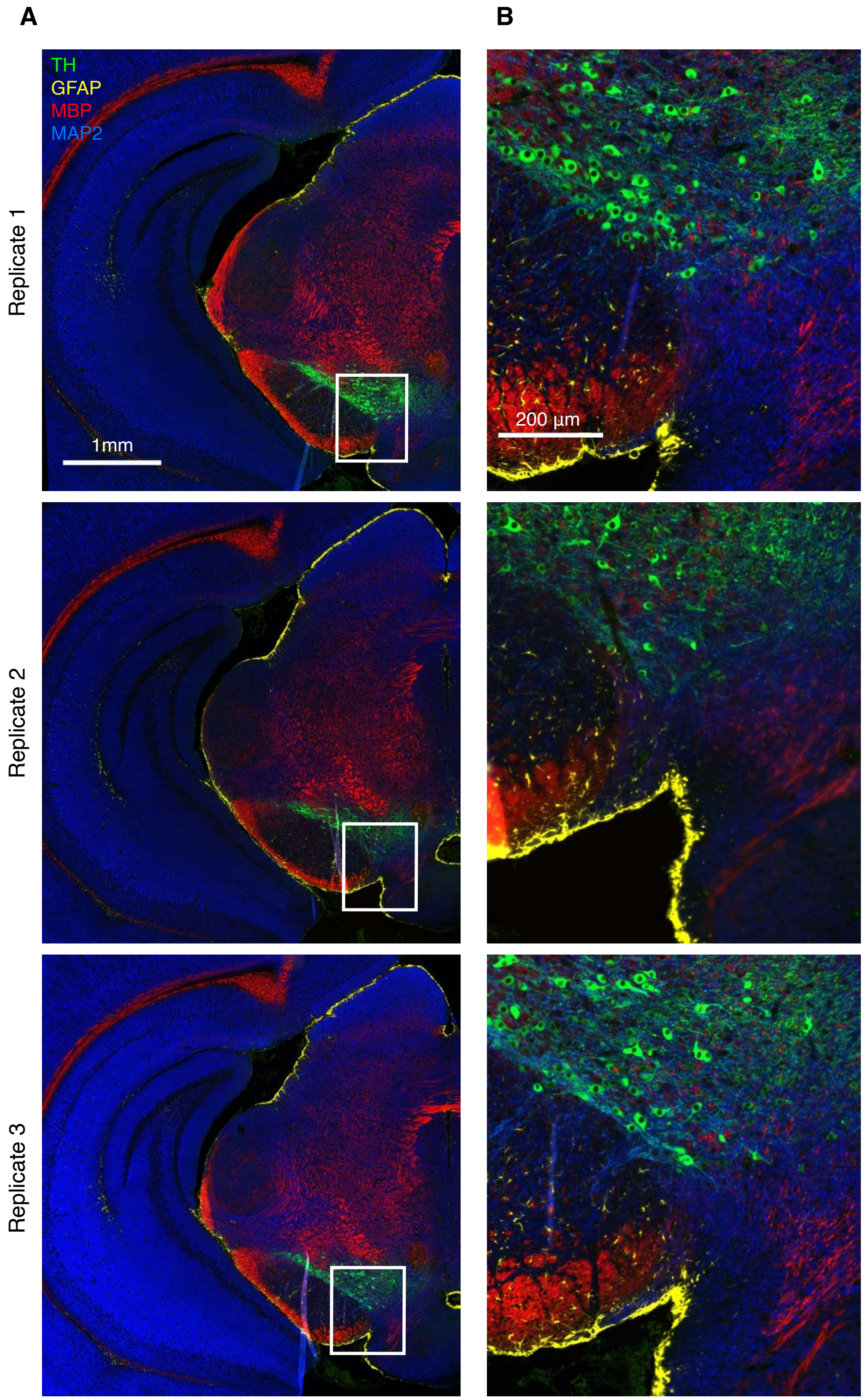

**Figure S5. Replicates for 4-plex protein imaging using HCR 1°IHC in FFPE mouse brain sections (cf. Figures 2DE).** (A) 4-channel epifluorescence images for 3 replicate FFPE mouse brain sections. (B) Zoom of the depicted region. Ch1: target protein TH (Alexa488). Ch2: target protein GFAP (Alexa546). Ch3: target protein MBP (Alexa647). Ch4: target protein MAP2 (Alexa750). Sample: FFPE C57BL/6 mouse brain section (coronal); thickness: 5 $\mu$m.

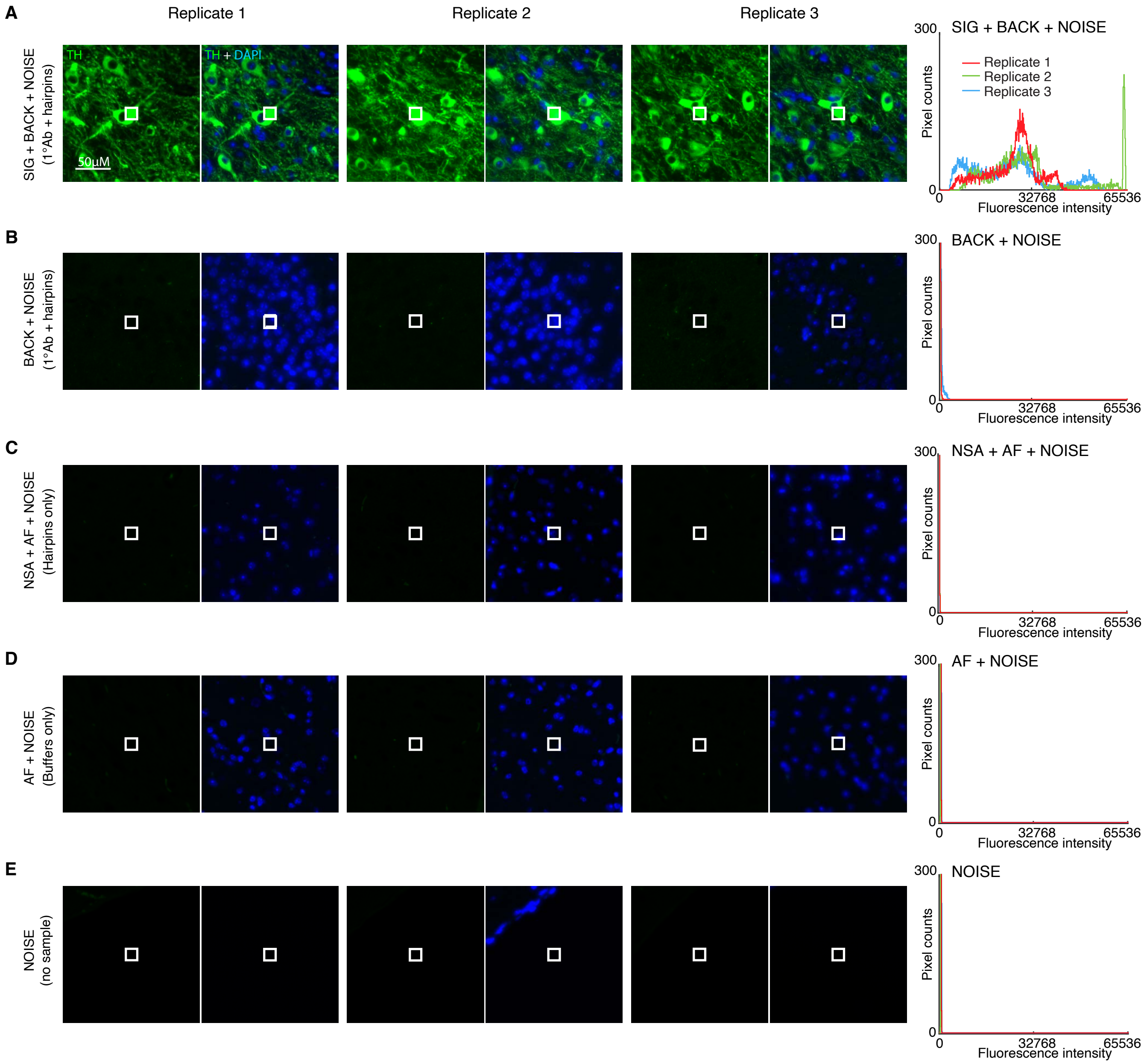

**Figure S6. Measurement of signal, background, background components, and noise for target protein TH using HCR 1°IHC in FFPE mouse brain sections (cf. Figures 2DE).** Use experiment of Type 1 in Table S7A (1°Ab probe + hairpins) to measure (A) SIG+BACK+NOISE in a region of high expression and (B) BACK+NOISE in a region of no/low expression. (C) Use experiment of Type 2 in Table S7B (no probes, hairpins only) to measure NSA+AF+NOISE in a region of high expression. Use experiment of Type 3 in Table S7B (no probes, no hairpins) to measure (D) AF+NOISE in a region of high expression and (E) NOISE in a region with no sample. Left: epifluorescence image collected with the microscope exposure time optimized to avoid saturating SIG+BACK+NOISE pixels; DAPI channel facilitates placement of rectangles. Right: pixel intensity histograms for representative regions (three rectangles per experiment type for each of three replicate FFPE mouse brain sections). Ch1: target protein TH (Alexa 488). Ch5: DAPI. Sample: FFPE C57BL/6 mouse brain section (coronal); thickness: 5 $\mu$m.

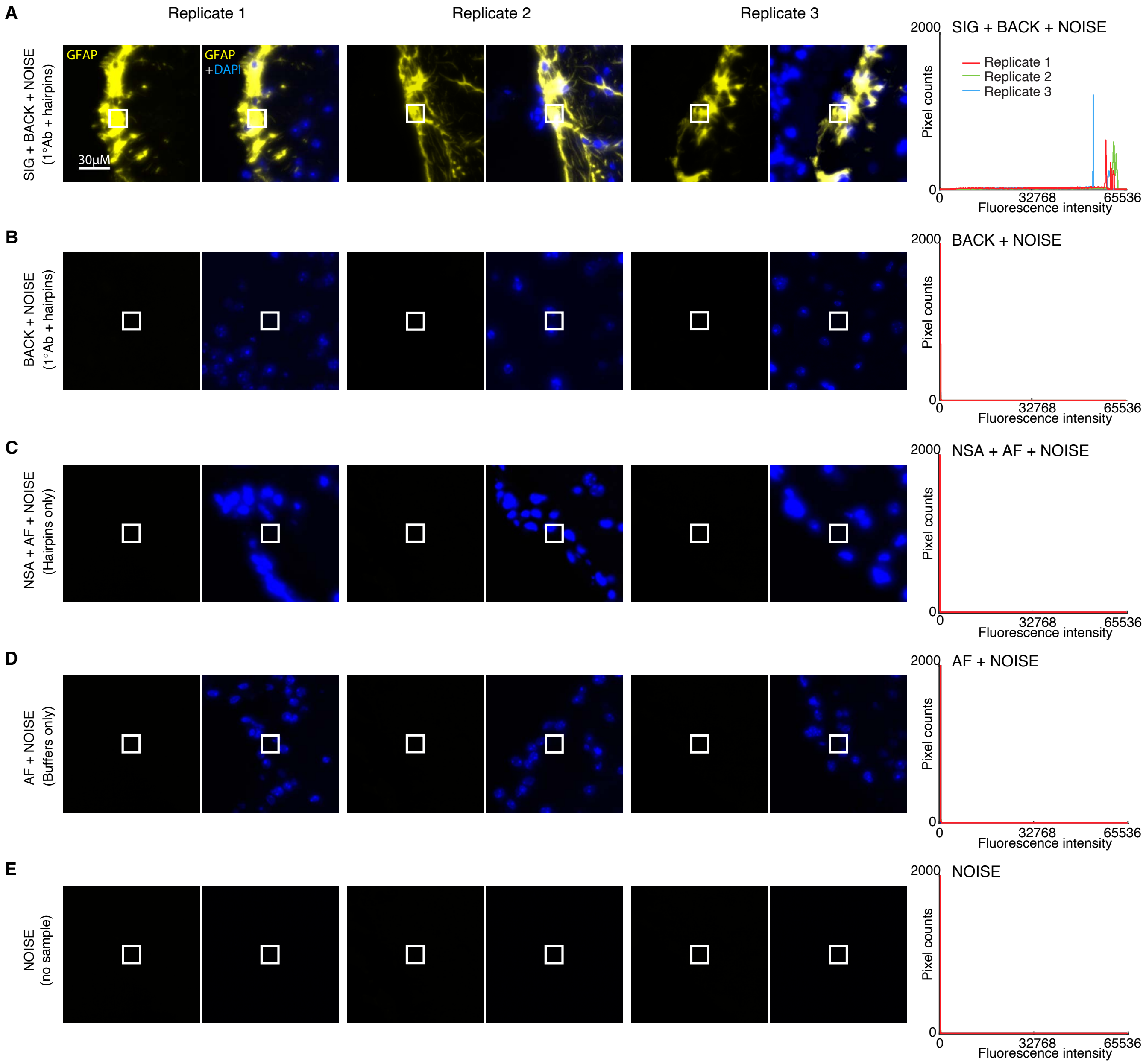

**Figure S7. Measurement of signal, background, background components, and noise for target protein GFAP using HCR 1°IHC in FFPE mouse brain sections (cf. Figures 2DE).** Use experiment of Type 1 in Table S7A (1°Ab probe + hairpins) to measure (A) SIG+BACK+NOISE in a region of high expression and (B) BACK+NOISE in a region of no/low expression. (C) Use experiment of Type 2 in Table S7B (no probes, hairpins only) to measure NSA+AF+NOISE in a region of high expression. Use experiment of Type 3 in Table S7B (no probes, no hairpins) to measure (D) AF+NOISE in a region of high expression and (E) NOISE in a region with no sample. Left: epifluorescence image collected with the microscope exposure time optimized to avoid saturating SIG+BACK+NOISE pixels; DAPI channel facilitates placement of rectangles. Right: pixel intensity histograms for representative regions (three rectangles per experiment type for each of three replicate FFPE mouse brain sections). Ch2: target protein GFAP (Alexa546). Ch5: DAPI. Sample: FFPE C57BL/6 mouse brain section (coronal); thickness: 5 $\mu$m.

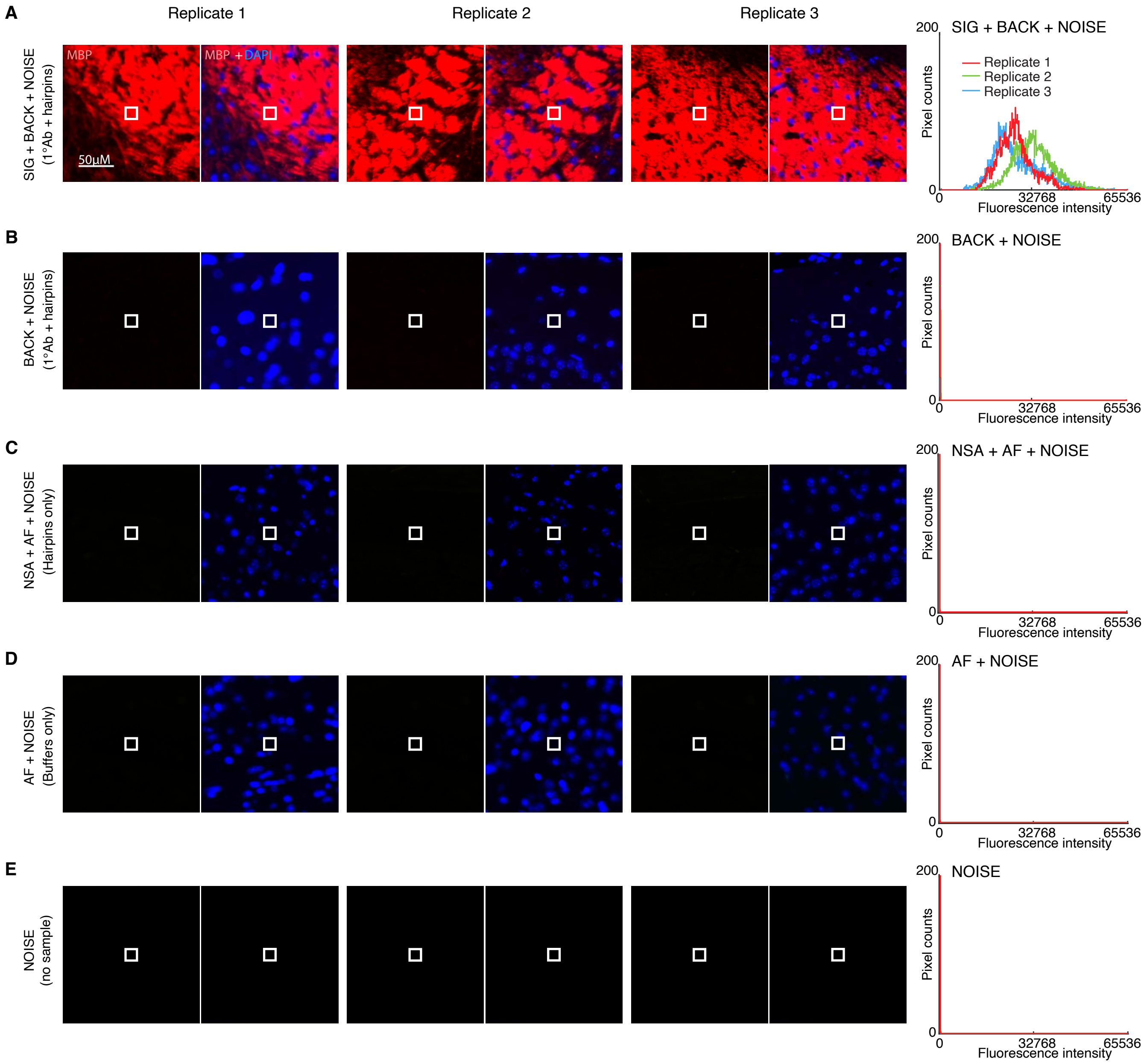

**Figure S8. Measurement of signal, background, background components, and noise for target protein MBP using HCR 1°IHC in FFPE mouse brain sections (cf. Figures 2DE).** Use experiment of Type 1 in Table S7A (1°Ab probe + hairpins) to measure (A) SIG+BACK+NOISE in a region of high expression and (B) BACK+NOISE in a region of no/low expression. (C) Use experiment of Type 2 in Table S7B (no probes, hairpins only) to measure NSA+AF+NOISE in a region of high expression. Use experiment of Type 3 in Table S7B (no probes, no hairpins) to measure (D) AF+NOISE in a region of high expression and (E) NOISE in a region with no sample. Left: epifluorescence image collected with the microscope exposure time optimized to avoid saturating SIG+BACK+NOISE pixels; DAPI channel facilitates placement of rectangles. Right: pixel intensity histograms for representative regions (three rectangles per experiment type for each of three replicate FFPE mouse brain sections). Ch3: target protein MBP (Alexa647). Ch5: DAPI. Sample: FFPE C57BL/6 mouse brain section (coronal); thickness: 5 $\mu$m.

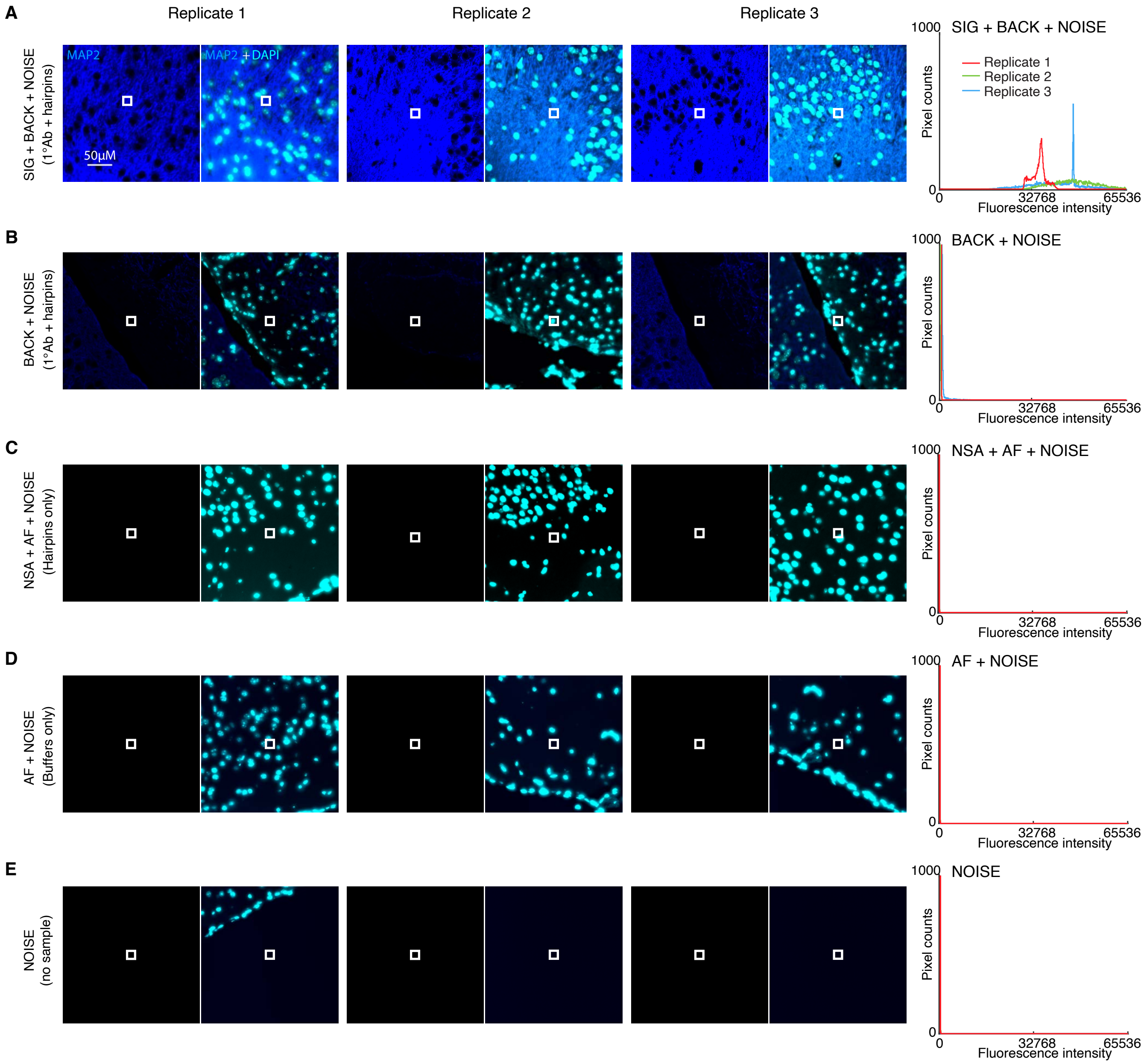

**Figure S9. Measurement of signal, background, background components, and noise for target protein MAP2 using HCR 1°IHC in FFPE mouse brain sections (cf. Figures 2DE).** Use experiment of Type 1 in Table S7A (1°Ab probe + hairpins) to measure (A) SIG+BACK+NOISE in a region of high expression and (B) BACK+NOISE in a region of no/low expression. (C) Use experiment of Type 2 in Table S7B (no probes, hairpins only) to measure NSA+AF+NOISE in a region of high expression. Use experiment of Type 3 in Table S7B (no probes, no hairpins) to measure (D) AF+NOISE in a region of high expression and (E) NOISE in a region with no sample. Left: epifluorescence image collected with the microscope exposure time optimized to avoid saturating SIG+BACK+NOISE pixels; DAPI channel facilitates placement of rectangles. Right: pixel intensity histograms for representative regions (three rectangles per experiment type for each of three replicate FFPE mouse brain sections). Ch4: target protein MAP2 (Alexa750). Ch5: DAPI. Sample: FFPE C57BL/6 mouse brain section (coronal); thickness: 5 $\mu$m.

| | Quantity | Ch1: TH B1-Alexa488 | Ch2: GFAP B3-Alexa546 | Ch3: MBP B5-Alexa647 | Ch4: MAP2 B4-Alexa750 | Reagents 1° Ab-init | Reagents Hairpins | Expression region | Figure panel |
|---|---|---|---|---|---|---|---|---|---|
| **A** | SIG+BACK+NOISE | 27 000 ± 4000 | 48 000 ± 5000 | 28 000 ± 4000 | 41 000 ± 6000 | ✓ | ✓ | high | A |
| | BACK+NOISE | 1060 ± 90 | 260 ± 30 | 720 ± 90 | 1500 ± 300 | ✓ | ✓ | low/no | B |
| | NOISE | 190 ± 16 | 94 ± 5 | 249 ± 18 | 119 ± 3 | ✓ | ✓ | no sample | E |
| | SIG | 26 000 ± 4000 | 48 000 ± 5000 | 28 000 ± 4000 | 40 000 ± 6000 | | | | |
| | BACK | 870 ± 90 | 170 ± 30 | 500 ± 100 | 1300 ± 300 | | | | |
| | SIG/BACK | 30 ± 5 | 290 ± 60 | 58 ± 14 | 30 ± 8 | | | | |
| **B** | NSA+AF+NOISE | 650 ± 20 | 145 ± 13 | 380 ± 30 | 147 ± 8 | | ✓ | high | C |
| | AF+NOISE | 600 ± 20 | 128 ± 13 | 303 ± 3 | 115 ± 2 | | | high | D |
| | NSD | 420 ± 90 | 110 ± 30 | 300 ± 100 | 1300 ± 300 | | | | |
| | NSA | 50 ± 30 | 18 ± 18 | 80 ± 30 | 30 ± 8 | | | | |
| | AF | 410 ± 30 | 34 ± 14 | 54 ± 19 | $< 4$ | | | | |

**Table S13. Estimated signal-to-background, background components, and noise for 4-plex protein imaging using HCR 1°IHC in FFPE mouse brain sections (cf. Figures 2DE).** (A) Estimated signal-to-background (SIG/BACK) based on methods of Section S2.6.2. (B) Estimated background components (AF, NSA, NSD) based on methods of Section S2.6.3. Mean ± standard error of the mean, $N = 3$ replicate FFPE mouse brain sections. Analysis based on representative rectangular regions (examples depicted in Figures S6–S9).

#### S5.3 Replicates, signal, background, background components, and noise for multiplexed HCR 2°IHC (cf. Figure 3)

##### S5.3.1 Mammalian cells on a slide

For 3-plex protein imaging using HCR 2°IHC in mammalian cells on a slide, the 4 channels are (3 proteins + DAPI):

- **Ch1:** Target protein PCNA, probe 1°mAb mouse IgG2a anti-PCNA, probe 2°pAb goat anti-mouse Fc$\gamma$ subclass 2a specific labeled with B5 initiator, amplifier B5-Alexa647.
- **Ch2:** Target protein HSP60, probe 1°mAb probe rabbit anti-HSP60, probe 2°pAb donkey anti-rabbit labeled with B3 initiator, amplifier B3-Alexa546.
- **Ch3:** Target protein SC35, probe 1°mAb mouse IgG1 anti-SC35, probe 2°pAb goat anti-mouse Fc$\gamma$ subclass 1 specific labeled with B2 initiator, amplifier B2-Alexa488.
- **Ch4:** DAPI.

Additional studies are presented as follows:

- Figure S10 displays 3-plex images for $N = 3$ replicate wells on a multi-well slide (cf. Figure 3C).
- Figures S11–S13 displays representative regions of individual channels used for measurement of signal and background for each target.
- Table S14 displays estimated values for signal, background, and signal-to-background for each target.

**Protocol:** HCR 2°IHC (Section S4.1) using unlabeled primary antibody probes and initiator-labeled secondary antibody probes with HCR signal amplification for all targets simultaneously.
**Sample:** HeLa cells.
**Microscopy:** Confocal.

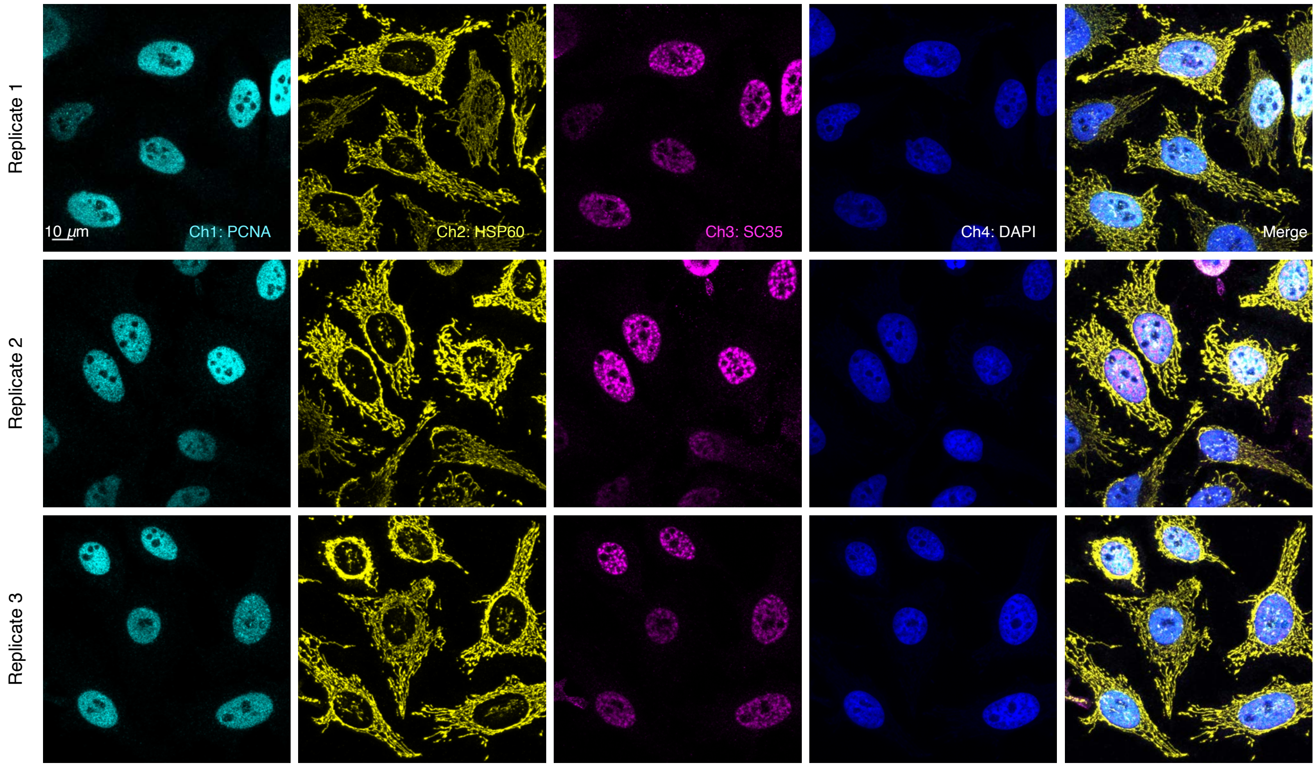

**Figure S10. Replicates for 3-plex protein imaging using HCR 2°IHC in mammalian cells on a slide (cf. Figures 3C).** 4-channel confocal images for 3 replicate wells on a multi-well slide; maximum intensity z-projection. Ch1: target protein PCNA (Alexa647). Ch2: target protein HSP60 (Alexa546). Ch3: target protein SC35 (Alexa488). Ch4: DAPI. Sample: HeLa cells.

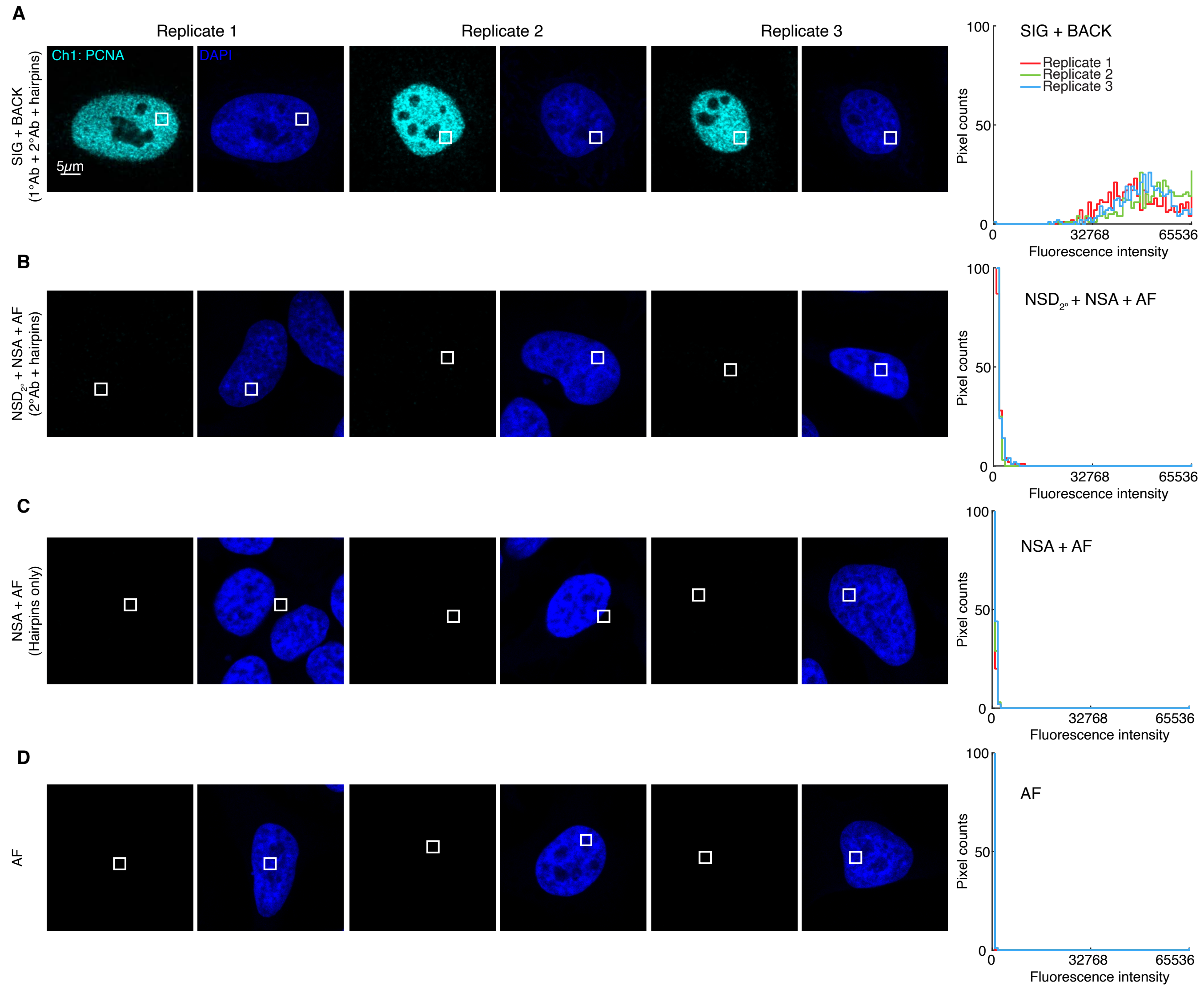

**Figure S11. Measurement of signal, background, and background components for target protein PCNA using HCR 2°IHC in mammalian cells on a slide (cf. Figure 3C).** (A) Use experiment of Type 1 in Table S8A (1°Ab probe + 2°Ab probe + hairpins) to measure SIG+BACK in a region of high expression. (B) Use experiment of Type 4 in Table S8B (2°Ab probes + hairpins) to measure $NSD_{2^\circ}$+NSA+AF in a region of maximum background. (C) Use experiment of Type 2 in Table S8B (no probes, hairpins only) to measure NSA+AF in a region of maximum background. (D) Use experiment of Type 3 in Table S8B (no probes, no hairpins) to measure AF in a region of maximum background. Left: confocal image collected with the microscope gain optimized to avoid saturating SIG+BACK pixels; DAPI channel facilitates placement of rectangles; single optical section. Right: pixel intensity histograms for representative regions (one rectangle in each of 5 individual cells in each of 3 replicate wells on a multi-well slide). Ch1: target protein PCNA (Alexa647). Ch4: DAPI. Sample: HeLa cells.

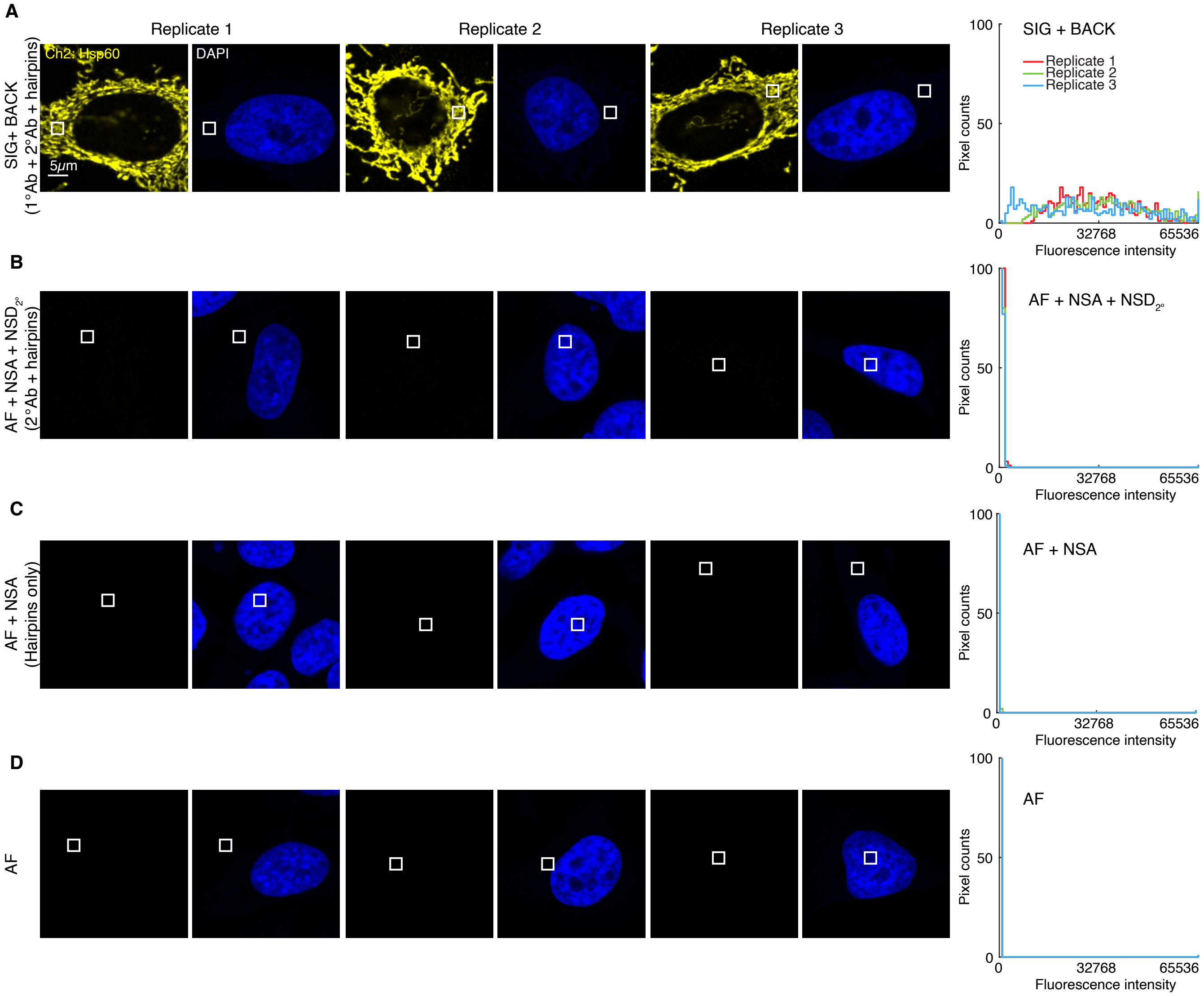

**Figure S12. Measurement of signal, background, and background components for target protein HSP60 using HCR 2°IHC in mammalian cells on a slide (cf. Figure 3C).** (A) Use experiment of Type 1 in Table S8A (1°Ab probe + 2°Ab probe + hairpins) to measure SIG+BACK in a region of high expression. (B) Use experiment of Type 4 in Table S8B (2°Ab probes + hairpins) to measure $NSD_{2^\circ}$+NSA+AF in a region of maximum background. (C) Use experiment of Type 2 in Table S8B (no probes, hairpins only) to measure NSA+AF in a region of maximum background. (D) Use experiment of Type 3 in Table S8B (no probes, no hairpins) to measure AF in a region of maximum background. Left: confocal image collected with the microscope gain optimized to avoid saturating SIG+BACK pixels; DAPI channel facilitates placement of rectangles; single optical section. Right: pixel intensity histograms for representative regions (one rectangle in each of 5 individual cells in each of 3 replicate wells on a multi-well slide). Ch2: target protein HSP60 (Alexa546). Ch4: DAPI. Sample: HeLa cells.

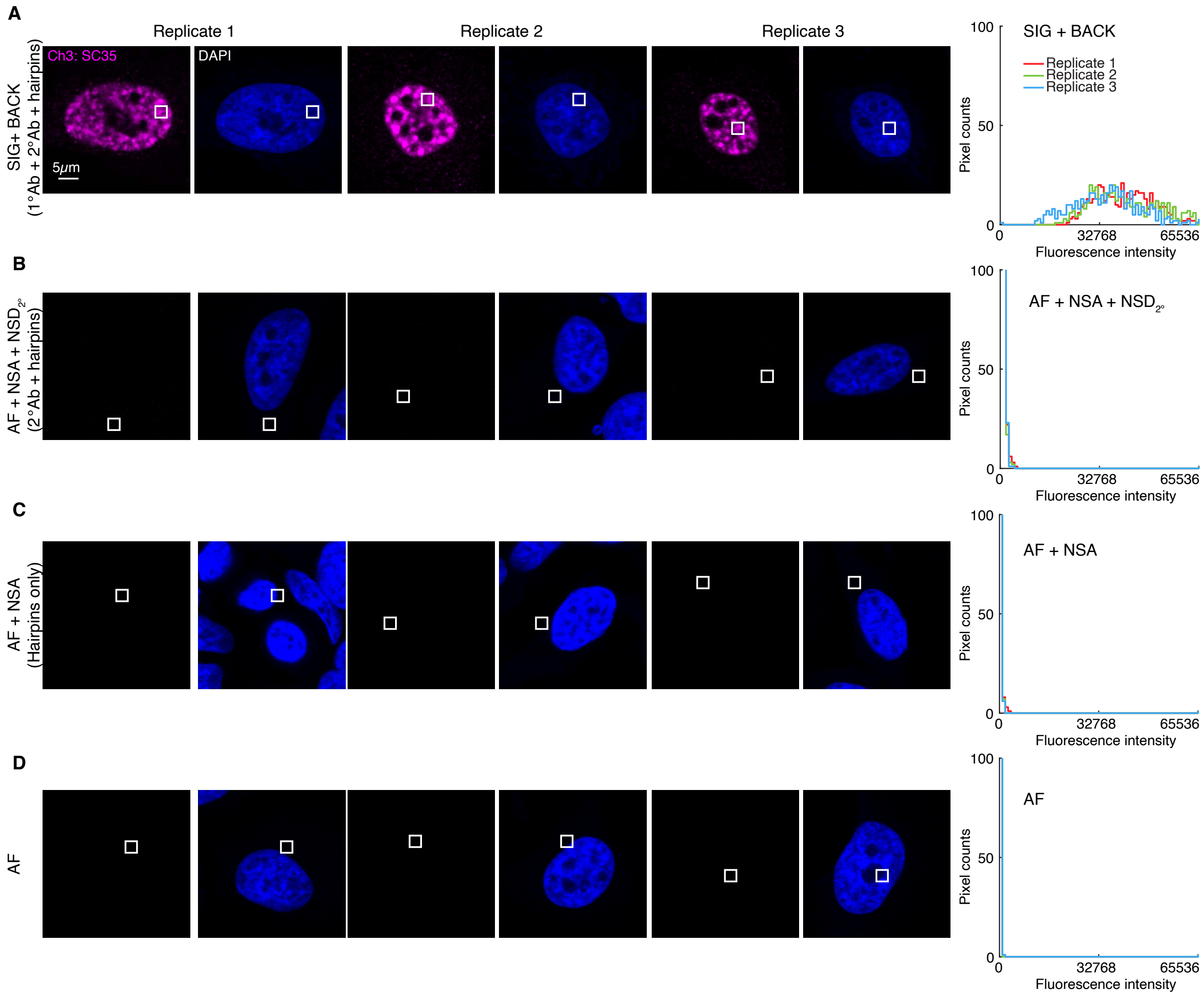

**Figure S13. Measurement of signal, background, and background components for target protein SC35 using HCR 2°IHC in mammalian cells on a slide (cf. Figure 3C).** (A) Use experiment of Type 1 in Table S8A (1°Ab probe + 2°Ab probe + hairpins) to measure SIG+BACK in a region of high expression. (B) Use experiment of Type 4 in Table S8B (2°Ab probes + hairpins) to measure $NSD_{2^\circ}$+NSA+AF in a region of maximum background. (C) Use experiment of Type 2 in Table S8B (no probes, hairpins only) to measure NSA+AF in a region of maximum background. (D) Use experiment of Type 3 in Table S8B (no probes, no hairpins) to measure AF in a region of maximum background. Left: confocal image collected with the microscope gain optimized to avoid saturating SIG+BACK pixels; DAPI channel facilitates placement of rectangles; single optical section. Right: pixel intensity histograms for representative regions (one rectangle in each of 5 individual cells in each of 3 replicate wells on a multi-well slide). Ch3: target protein SC35 (Alexa488). Ch4: DAPI. Sample: HeLa cells.

| | | Ch1: PCNA | Ch2: Hsp60 | Ch3: SC35 | Reagents | | | Figure |
|---|---|---|---|---|---|---|---|---|
| | Quantity | B5-Alexa647 | B3-Alexa546 | B2-Alexa488 | 1°Ab | 2°Ab-init | Hairpins | panel |
| **A** | SIG+BACK | 43 300 ± 1700 | 31 500 ± 1200 | 33 000 ± 2000 | ✓ | ✓ | ✓ | A |
| | SIG | 42 800 ± 1700 | 31 200 ± 1200 | 33 000 ± 2000 | | | | |
| | SIG/BACK | 87 ± 6 | 106 ± 7 | 69 ± 6 | | | | |
| **B** | $NSD_{2^\circ}$+NSA+AF | 490 ± 30 | 293 ± 17 | 470 ± 30 | | ✓ | ✓ | B |
| | NSA+AF | 138 ± 6 | 79 ± 3 | 113 ± 3 | | | ✓ | C |
| | AF | 64.5 ± 0.5 | 51.5 ± 0.3 | 100 ± 3 | | | | D |
| | $NSD_{2^\circ}$ | 360 ± 30 | 215 ± 17 | 360 ± 30 | | | | |
| | NSA | 74 ± 6 | 27 ± 3 | 13 ± 5 | | | | |

**Table S14. Estimated signal-to-background and background components for 3-plex protein imaging using HCR 2°IHC in mammalian cells on a slide (cf. Figure 3C).** (A) Estimated signal-to-background (SIG/BACK) based on methods of Section S2.6.2. The signal estimate SIG is calculated using the background approximation BACK ≈ $NSD_{2^\circ}$+NSA+AF. (B) Estimated background components (AF, NSA, $NSD_{2^\circ}$) based on methods of Section S2.6.3. Instrument noise is negligible using confocal microscopy so calculations use the approximation NOISE ≈ 0. Mean ± standard error of the mean, $N = 15$ representative rectangular regions (one rectangle in each of 5 individual cells in each of 3 replicate wells on a multi-well slide). Analysis based on rectangular regions depicted in Figures S11–S13.

##### S5.3.2 FFPE mouse brain sections

For 4-plex protein imaging using HCR 2°IHC in FFPE mouse brain sections, the 5 channels are (4 proteins + DAPI):

- **Ch1:** Target protein TH, probe 1°pAb sheep IgG anti-TH, probe 2°pAb donkey anti-sheep IgG labeled with B4 initiator, amplifier B4-Alexa488.
- **Ch2:** Target protein GFAP, probe 1°pAb chicken IgY anti-GFAP, probe 2°pAb donkey anti-chicken IgG labeled with B1 initiator, amplifier B1-Alexa546.
- **Ch3:** Target protein PVALB, probe 1°mAb rabbit IgG anti-PVALB, probe 2°pAb donkey anti-rabbit IgG labeled with B5 initiator, amplifier B5-Alexa647.
- **Ch4:** Target protein MBP, probe 1°mAb rat IgG2a anti-MBP, probe 2°pAb donkey anti-rat IgG labeled with B3 initiator, amplifier B3-Alexa750.
- **Ch5:** DAPI.

Additional studies are presented as follows:

- Figure S14 displays 4-plex images for $N = 3$ replicate FFPE mouse brain sections (cf. Figure 2C).
- Figures S15–S18 display representative regions of individual channels used for measurement of signal and background for each target.
- Table S15 displays estimated values for signal, background, and signal-to-background for each target.

**Protocol:** HCR 2°IHC (Section S4.2) using unlabeled primary antibody probes and initiator-labeled secondary antibody probes with HCR signal amplification for all targets simultaneously.
**Sample:** FFPE C57BL/6 mouse brain section (coronal); thickness: 5 $\mu$m.
**Microscopy:** Epifluorescence.

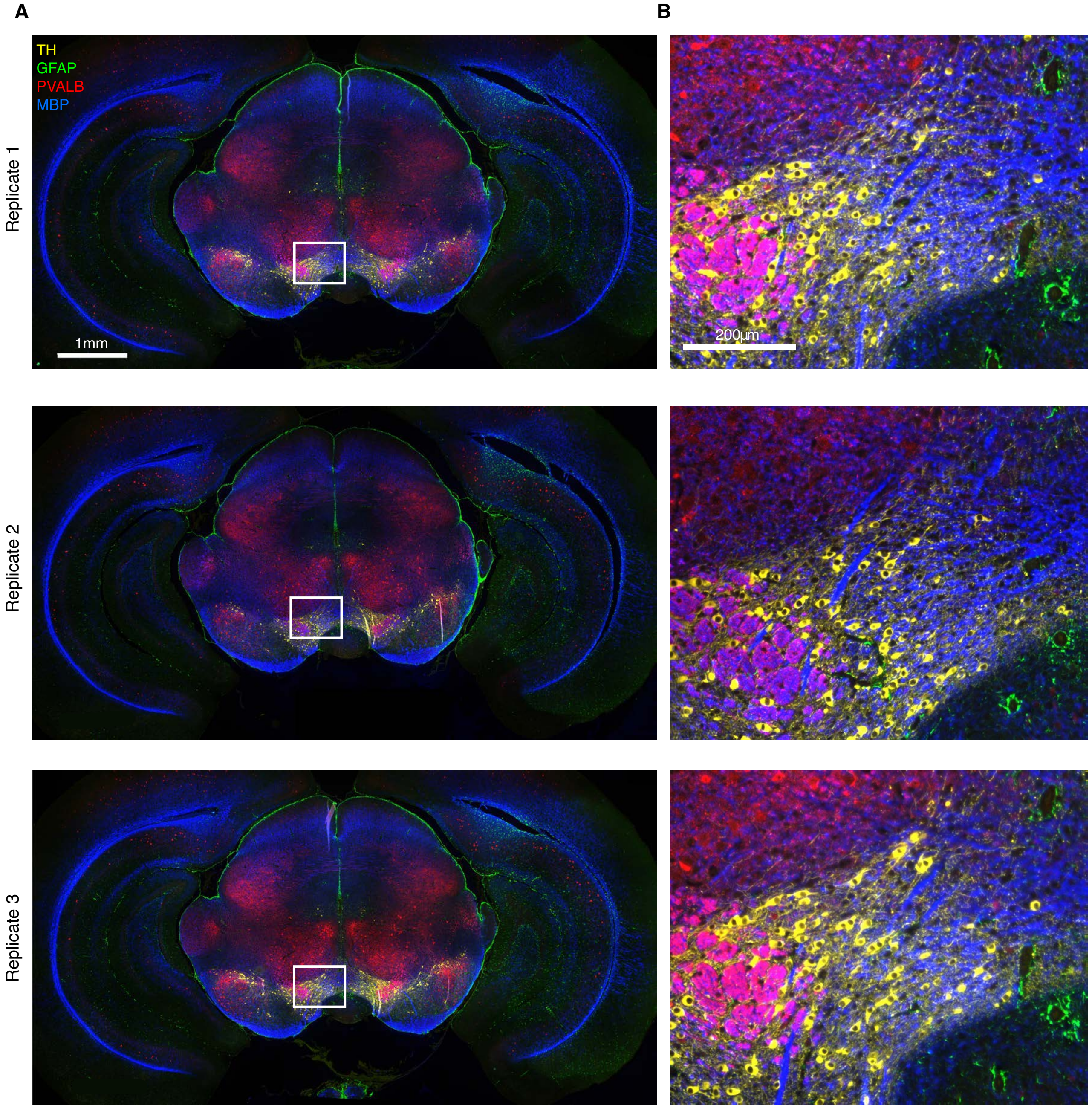

**Figure S14. Replicates for 4-plex protein imaging using HCR 2°IHC in FFPE mouse brain sections (cf. Figures 3DE).** (A) 4-channel epifluorescence images for 3 replicate FFPE mouse brain sections. (B) Zoom of the depicted region. Ch1: target protein TH (Alexa488). Ch2: target protein GFAP (Alexa546). Ch3: target protein PVALB (Alexa647). Ch4: target protein MBP (Alexa750). Sample: FFPE C57BL/6 mouse brain section (coronal); thickness: 5 $\mu$m.

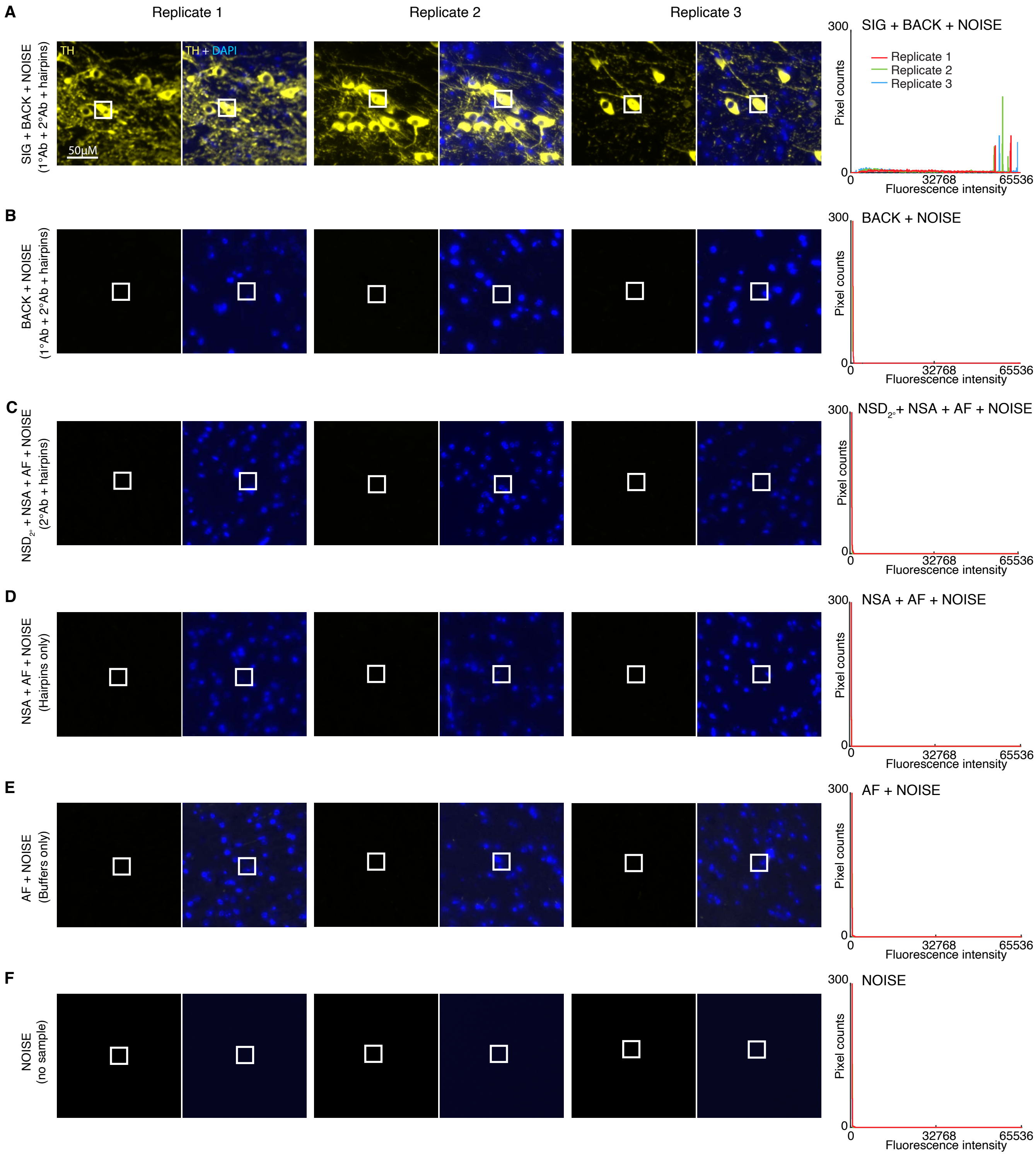

**Figure S15. Measurement of signal, background, background components, and noise for protein target TH using HCR 2°IHC in FFPE mouse brain sections (cf. Figures 3DE).** Use experiment of Type 1 in Table S8A (1°Ab probe + 2°Ab probe + hairpins) to measure (A) SIG+BACK+NOISE in a region of high expression and (B) BACK+NOISE in a region of no/low expression. (C) Use experiment of Type 4 in Table S8B (2°Ab probes + hairpins) to measure $NSD_{2^\circ}$+NSA+AF+NOISE in a region of high expression. (D) Use experiment of Type 2 in Table S8B (no probes, hairpins only) to measure NSA+AF+NOISE in a region of high expression. Use experiment of Type 3 in Table S8B (no probes, no hairpins) to measure (E) AF+NOISE in a region of high expression and (F) NOISE in a region with no sample. Left: epifluorescence image collected with the microscope exposure time optimized to avoid saturating SIG+BACK+NOISE pixels; DAPI channel facilitates placement of rectangles. Right: pixel intensity histograms for representative regions (three rectangles per experiment type for each of three replicate FFPE mouse brain sections). Ch1: target protein TH (Alexa488). Ch5: DAPI. Sample: FFPE C57BL/6 mouse brain section (coronal); thickness: 5 $\mu$m.

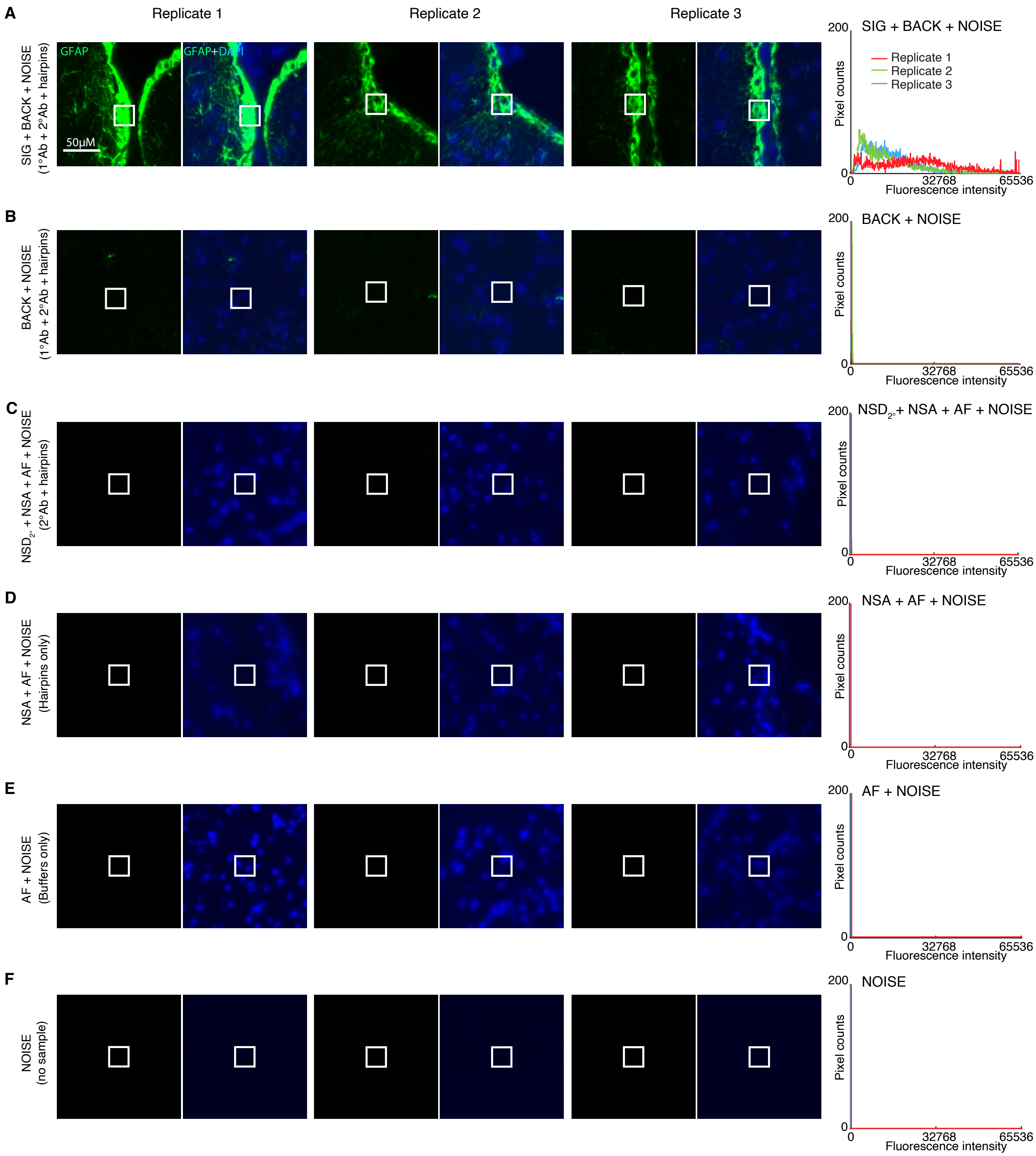

**Figure S16. Measurement of signal, background, background components, and noise for protein target GFAP using HCR 2°IHC in FFPE mouse brain sections (cf. Figures 3DE).** Use experiment of Type 1 in Table S8A (1°Ab probe + 2°Ab probe + hairpins) to measure (A) SIG+BACK+NOISE in a region of high expression and (B) BACK+NOISE in a region of no/low expression. (C) Use experiment of Type 4 in Table S8B (2°Ab probes + hairpins) to measure $NSD_{2°}$+NSA+AF+NOISE in a region of high expression. (D) Use experiment of Type 2 in Table S8B (no probes, hairpins only) to measure NSA+AF+NOISE in a region of high expression. Use experiment of Type 3 in Table S8B (no probes, no hairpins) to measure (E) AF+NOISE in a region of high expression and (F) NOISE in a region with no sample. Left: epifluorescence image collected with the microscope exposure time optimized to avoid saturating SIG+BACK+NOISE pixels; DAPI channel facilitates placement of rectangles. Right: pixel intensity histograms for representative regions (three rectangles per experiment type for each of three replicate FFPE mouse brain sections). Ch2: target protein GFAP (Alexa546). Ch5: DAPI. Sample: FFPE C57BL/6 mouse brain section (coronal); thickness: 5 $\mu$m.

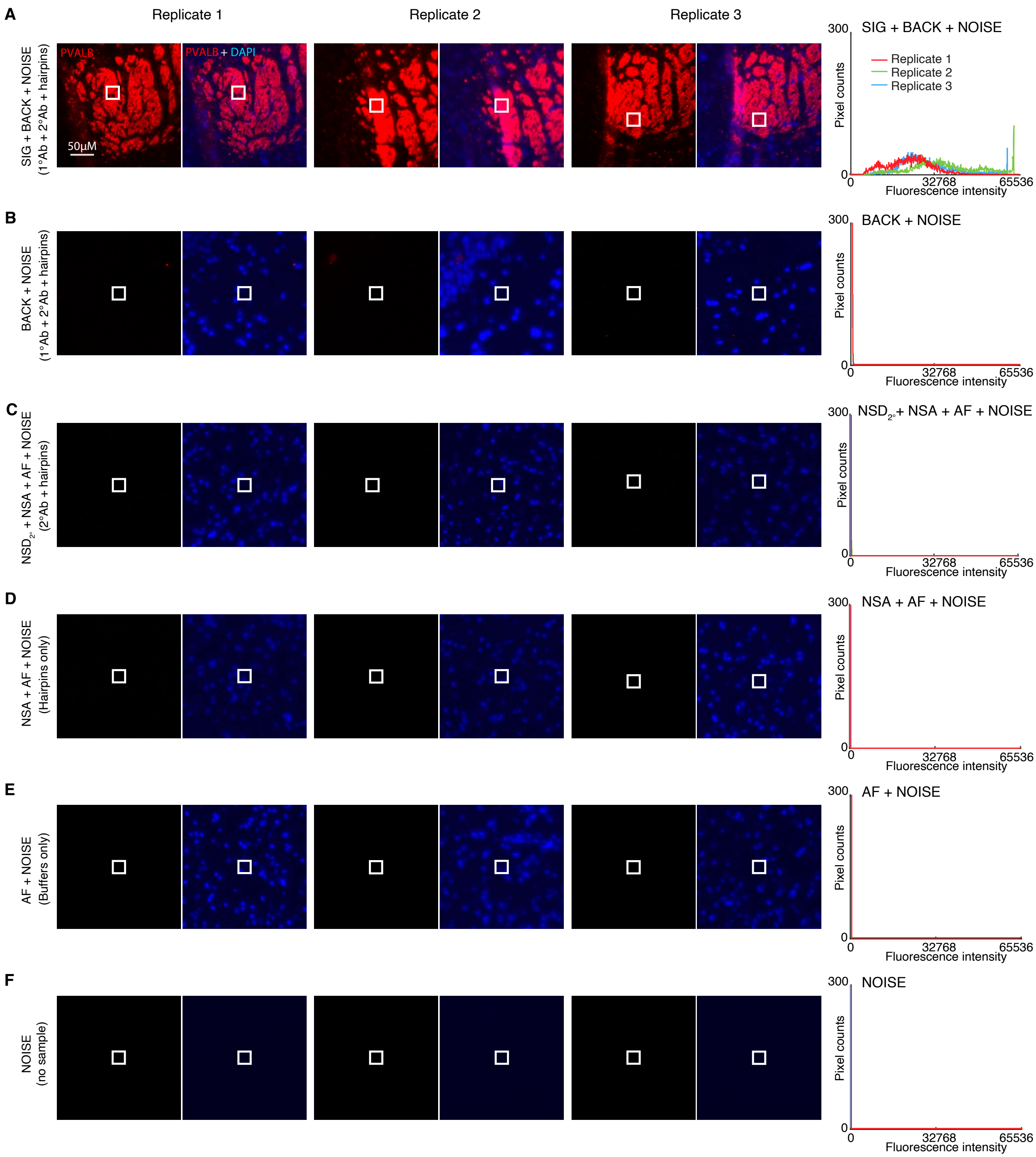

**Figure S17. Measurement of signal, background, background components, and noise for protein target PVALB using HCR 2°IHC in FFPE mouse brain sections (cf. Figures 3DE).** Use experiment of Type 1 in Table S8A (1°Ab probe + 2°Ab probe + hairpins) to measure (A) SIG+BACK+NOISE in a region of high expression and (B) BACK+NOISE in a region of no/low expression. (C) Use experiment of Type 4 in Table S8B (2°Ab probes + hairpins) to measure $NSD_{2^\circ}$+NSA+AF+NOISE in a region of high expression. (D) Use experiment of Type 2 in Table S8B (no probes, hairpins only) to measure NSA+AF+NOISE in a region of high expression. Use experiment of Type 3 in Table S8B (no probes, no hairpins) to measure (E) AF+NOISE in a region of high expression and (F) NOISE in a region with no sample. Left: epifluorescence image collected with the microscope exposure time optimized to avoid saturating SIG+BACK+NOISE pixels; DAPI channel facilitates placement of rectangles. Right: pixel intensity histograms for representative regions (three rectangles per experiment type for each of three replicate FFPE mouse brain sections). Ch3: target protein PVALB (Alexa647). Ch5: DAPI. Sample: FFPE C57BL/6 mouse brain section (coronal); thickness: 5 $\mu$m.

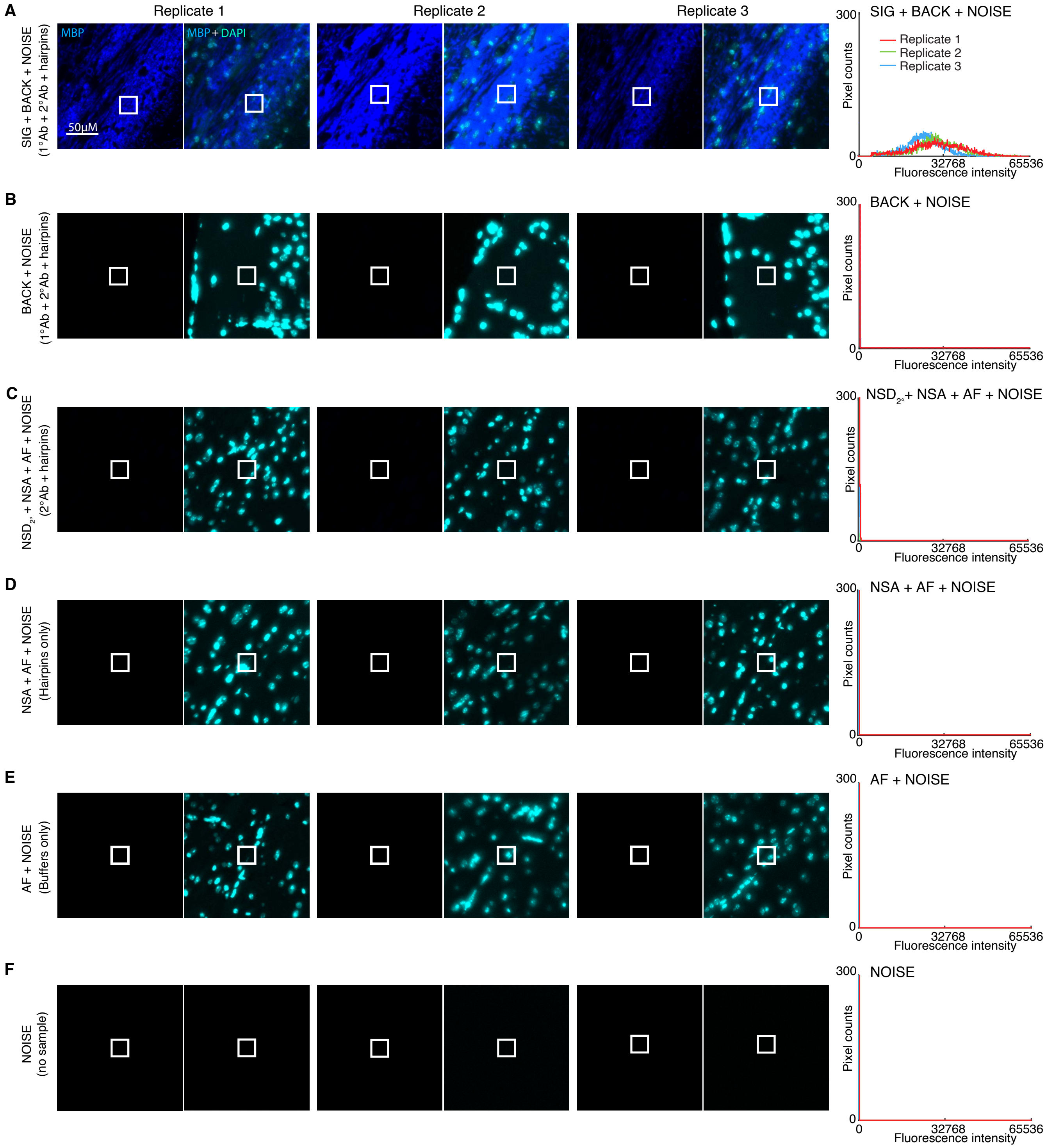

**Figure S18. Measurement of signal, background, background components, and noise for protein target MBP using HCR 2°IHC in FFPE mouse brain sections (cf. Figures 3DE).** Use experiment of Type 1 in Table S8A (1°Ab probe + 2°Ab probe + hairpins) to measure (A) SIG+BACK+NOISE in a region of high expression and (B) BACK+NOISE in a region of no/low expression. (C) Use experiment of Type 4 in Table S8B (2°Ab probes + hairpins) to measure $NSD_{2^\circ}$+NSA+AF+NOISE in a region of high expression. (D) Use experiment of Type 2 in Table S8B (no probes, hairpins only) to measure NSA+AF+NOISE in a region of high expression. Use experiment of Type 3 in Table S8B (no probes, no hairpins) to measure (E) AF+NOISE in a region of high expression and (F) NOISE in a region with no sample. Left: epifluorescence image collected with the microscope exposure time optimized to avoid saturating SIG+BACK+NOISE pixels; DAPI channel facilitates placement of rectangles. Right: pixel intensity histograms for representative regions (three rectangles per experiment type for each of three replicate FFPE mouse brain sections). Ch4: target protein MBP (Alexa750). Ch5: DAPI. Sample: FFPE C57BL/6 mouse brain section (coronal); thickness: 5 $\mu$m.

| | | Ch1: TH | Ch2: GFAP | Ch3: PVALB | Ch4: MBP | Reagents | | Expression | Figure | |
|---|---|---|---|---|---|---|---|---|---|---|
| | Quantity | B4-Alexa488 | B1-Alexa546 | B5-Alexa647 | B3-Alexa750 | 1°Ab | 2°Ab-init | Hairpins | region | panel |
| **A** | SIG+BACK+NOISE | 32 000 ± 3000 | 18 000 ± 6000 | 30 000 ± 8000 | 29 000 ± 3000 | ✓ | ✓ | ✓ | high | A |
| | BACK+NOISE | 640 ± 70 | 900 ± 200 | 340 ± 40 | 225 ± 12 | ✓ | ✓ | ✓ | low/no | B |
| | NOISE | 211 ± 16 | 114 ± 4 | 165 ± 2 | 116.5 ± 0.4 | ✓ | ✓ | ✓ | no sample | F |
| | SIG | 31 000 ± 3000 | 18 000 ± 6000 | 30 000 ± 8000 | 28 000 ± 3000 | | | | | |
| | BACK | 430 ± 80 | 800 ± 200 | 170 ± 40 | 109 ± 12 | | | | | |
| | SIG/BACK | 72 ± 14 | 23 ± 11 | 170 ± 60 | 260 ± 40 | | | | | |
| **B** | $NSD_{2^\circ}$+NSA+AF+NOISE | 540 ± 30 | 185 ± 11 | 297 ± 11 | 240 ± 30 | | ✓ | ✓ | high | C |
| | NSA+AF+NOISE | 500 ± 40 | 178 ± 2 | 237 ± 5 | 149 ± 4 | | | ✓ | high | D |
| | AF+NOISE | 490 ± 20 | 168 ± 6 | 217 ± 2 | 128 ± 3 | | | | high | E |
| | NSD | 140 ± 50 | 700 ± 200 | 100 ± 40 | 110 ± 30 | | | | | |
| | $NSD_{1^\circ}$ | 110 ± 30 | 700 ± 200 | 40 ± 40 | 10 ± 30 | | | | | |
| | $NSD_{2^\circ}$ | 40 ± 40 | $< 17$ | 60 ± 13 | 100 ± 30 | | | | | |
| | NSA | 10 ± 40 | 10 ± 6 | 19 ± 6 | 21 ± 5 | | | | | |
| | AF | 280 ± 30 | 54 ± 7 | 53 ± 3 | 12 ± 3 | | | | | |

**Table S15. Estimated signal-to-background, background components, and noise for 4-plex protein imaging using HCR 2°IHC in FFPE mouse brain sections (cf. Figures 3DE).** (A) Estimated signal-to-background (SIG/BACK) based on methods of Section S2.6.2. (B) Estimated background components (AF, NSA, $NSD_{1^\circ}$, $NSD_{2^\circ}$, NSD) based on methods of Section S2.6.3. Mean ± standard error of the mean, $N = 3$ replicate FFPE mouse brain sections. Analysis based on representative rectangular regions (examples depicted in Figures S15–S18).

#### S5.4 Protein imaging with high signal-to-background in whole-mount zebrafish embryos using HCR 2°IHC.

Here, we demonstrate protein imaging in whole-mount vertebrate embryos using HCR 2°IHC (cf. Figure 3). The reagents for these 1-channel studies are:

- **Ch1:** target protein Elavl3/Elavl4, probe 1°mAb mouse IgG2b anti-Elavl3/Elavl4, probe 2°pAb goat anti-mouse IgG2b labeled with B1 initiator, amplifier B1-Alexa647.

Additional studies are presented as follows:

- Figure S19 displays confocal images depicting representative regions used for measurement of signal and background for $N = 3$ replicate whole-mount zebrafish embryos.
- Table S16 displays estimated values for signal, background, and signal-to-background.

**Protocol:** HCR 2°IHC (Section S4.4) using unlabeled primary antibody probes and initiator-labeled secondary antibody probes with HCR signal amplification.
**Sample:** Whole-mount zebrafish embryos; fixed 27 hpf.
**Microscopy:** Confocal.

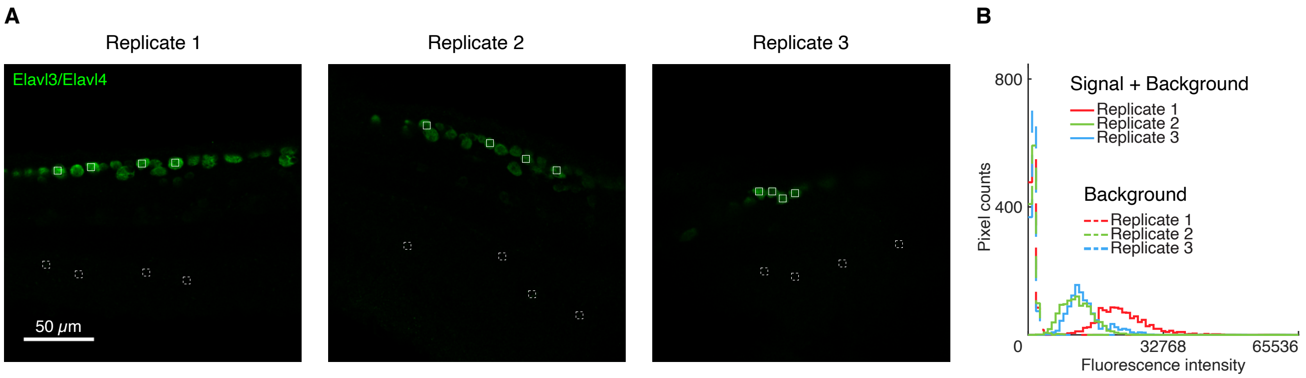

**Figure S19. Measurement of signal and background for protein imaging using HCR 2°IHC in whole-mount zebrafish embryos (cf. Figure 3).** Use experiment of Type 1 in Table S8A (1°Ab probe + 2°Ab probe + hairpins) to measure (A) SIG+BACK in regions of high expression and (B) BACK in regions of no/low expression. Confocal images. For each of three replicate embryos, a representative optical section was selected based on the expression depth of the target protein. Pixel size: $0.312 \times 0.312$ $\mu$m. (B) Pixel intensity histograms for SIG+BACK (pixels within solid boundary) and BACK (pixels within dashed boundary). Ch1: target protein Elavl3/Elavl4 (Alexa647). Sample: whole-mount zebrafish embryos; fixed 27 hpf.

| Target protein | Fluorophore | BACK | SIG+BACK | SIG | SIG/BACK |
|---|---|---|---|---|---|
| Elavl3/Elavl4 | Alexa647 | $800 \pm 100$ | $16\,000 \pm 3000$ | $15\,000 \pm 3000$ | $20 \pm 4$ |

**Table S16. Estimated signal-to-background for protein imaging using HCR 2°IHC in whole-mount zebrafish embryos (cf. Figure 3).** Instrument noise is negligible using confocal microscopy so calculations use the approximation NOISE $\approx 0$. Mean $\pm$ standard error of the mean, $N = 3$ replicate embryos. Analysis based on rectangular regions depicted in Figure S19 using methods of Section S2.6.2.

#### S5.5 Estimating HCR IHC polymer length (cf. Figures 2 and 3)

The gain due to HCR signal amplification corresponds to the mean HCR polymer length, which can be described in terms of the mean number of HCR hairpins per polymer. Here, we estimate HCR amplification gain in the context of HCR 1°IHC and HCR 2°IHC in both mammalian cells on a slide and FFPE mouse brain sections. For each method, we estimate:

- SIG using hairpins h1 and h2 to that HCR polymerization can proceed as normal.
- $\text{SIG}_{\text{h1}}$ using only hairpin h1 so that each HCR initiator can bind only one HCR hairpin and polymerization cannot proceed.

The HCR amplification gain is then estimated as $\text{SIG}/\text{SIG}_{\text{h1}}$. Results are summarized in Table S17.

| Method | Sample | Target protein | Amplifier | SIG | $\text{SIG}_{\text{h1}}$ | $\text{SIG}/\text{SIG}_{\text{h1}}$ | Table |
|---|---|---|---|---|---|---|---|
| HCR 1°IHC | mammalian cells on a slide | PCNA | B5-Alexa647 | $39\,700 \pm 900$ | $290 \pm 40$ | $134 \pm 17$ | S18 |
| HCR 2°IHC | mammalian cells on a slide | PCNA | B5-Alexa647 | $36\,000 \pm 2000$ | $280 \pm 40$ | $130 \pm 20$ | S19 |
| HCR 1°IHC | FFPE mouse brain sections | TH | B3-Alexa647 | $12\,000 \pm 1100$ | $51 \pm 4$ | $230 \pm 30$ | S20 |
| HCR 2°IHC | FFPE mouse brain sections | TH | B3-Alexa647 | $24\,000 \pm 3000$ | $53 \pm 4$ | $450 \pm 50$ | S21 |

**Table S17. Estimates of HCR amplification gain (mean polymer length) in the context of HCR 1°IHC and HCR 2°IHC in mammalian cells on a slide and FFPE mouse brain sections.** Mean $\pm$ standard error of the mean. For mammalian cells on a slide, $N = 15$ representative rectangular regions (one rectangle in each of 5 individual cells on each of 3 replicate wells on a multi-well slide). For FFPE mouse brain sections, $N = 3$ replicate sections. Analysis based on representative rectangular regions (examples depicted in Figures S20-S23) using methods of Sections S2.6.2 and S2.6.4.

##### S5.5.1 HCR 1°IHC in mammalian cells on a slide

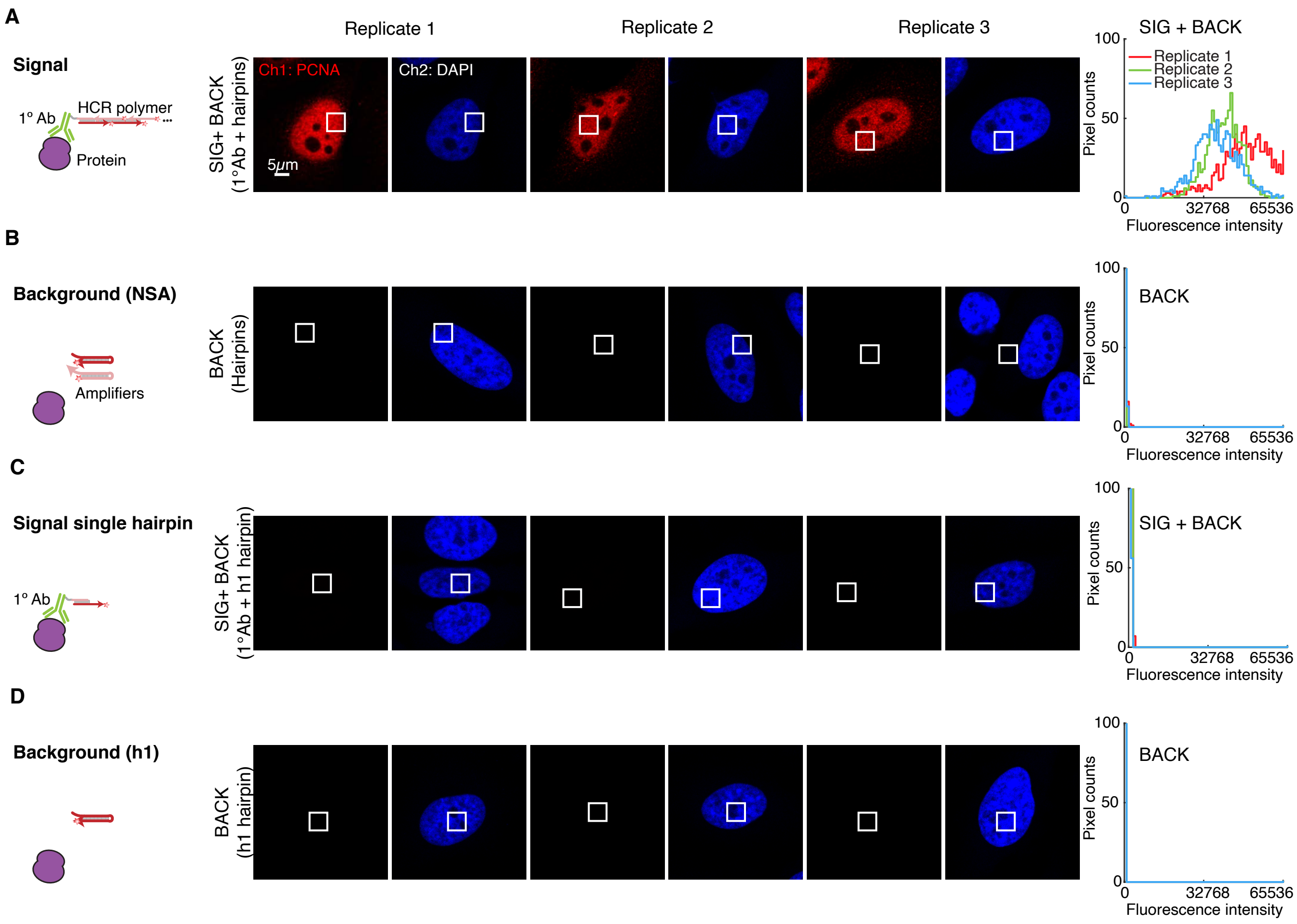

**Figure S20. Measurement of HCR amplification gain (mean polymer length) for HCR 1°IHC in mammalian cells on a slide (cf. Figure 2C).** (A,B) HCR 1°IHC. (C,D) HCR 1°IHC (h1 only). For each of 2 methods there are 2 rows: the top row measures SIG+BACK and the bottom row measures an approximation for BACK (see Table S18 for details on the background approximation used for each method). Left: Schematic of reagents used. Middle: confocal images collected with the microscope gain optimized to avoid saturating SIG+BACK pixels using HCR 1°IHC (panel A); DAPI channel facilitates placement of rectangles; single optical section. Pixel size: $0.198 \times 0.198$ $\mu$m. Right: pixel intensity histograms for representative regions (one rectangle in each of 5 individual cells in each of 3 replicate wells on a multi-well slide). Ch1: target protein PCNA (Alexa647). Ch2: DAPI. Sample: HeLa cells.

| | Quantity | Reagents | Value | Figure |
|---|---|---|---|---|
| **A** | SIG+BACK | 1°Ab-i1 + h1 + h2 | 39 700 ± 900 | S20A |
| | NSA+AF | h1 + h2 | 84 ± 2 | S20B |
| | SIG | | 39 600 ± 900 | |
| **B** | $\mathrm{SIG_{h1}}$+BACK | 1°Ab-i1 + h1 | 360 ± 40 | S20C |
| | $\mathrm{NSA_{h1}}$+AF | h1 | 61.0 ± 0.6 | S20D |
| | $\mathrm{SIG_{h1}}$ | | 290 ± 40 | |
| **C** | SIG/$\mathrm{SIG_{h1}}$ | | 134 ± 17 | |

**Table S18. Estimate of HCR amplification gain (mean polymer length) for HCR 1°IHC in mammalian cells on a slide (cf. Figure 2C).** (A) Estimated signal for HCR 1°IHC using HCR signal amplification (hairpins h1 and h2). The signal estimate SIG is calculated using the background approximation BACK $\approx$ NSA+AF. (B) Estimated signal $\mathrm{SIG_{h1}}$ without HCR signal amplification (using only hairpin h1 so that HCR polymerization is not possible and only a single h1 hairpin can bind initiator i1). The signal estimate $\mathrm{SIG_{h1}}$ is calculated using the background approximation $\mathrm{BACK_{h1}} \approx \mathrm{NSA_{h1}}$+AF. (C) Estimated HCR amplification gain SIG/$\mathrm{SIG_{h1}}$ (i.e., mean HCR polymer length). Instrument noise is negligible using confocal microscopy so calculations use the approximation NOISE $\approx 0$. Mean $\pm$ standard error of the mean, $N = 15$ representative rectangular regions (one rectangle in each of 5 individual cells on each of 3 replicate wells on a multi-well slide). Analysis based on representative rectangular regions (examples depicted in Figures S20) using methods of Sections S2.6.2 and S2.6.4.

##### S5.5.2 HCR 2°IHC in mammalian cells on a slide

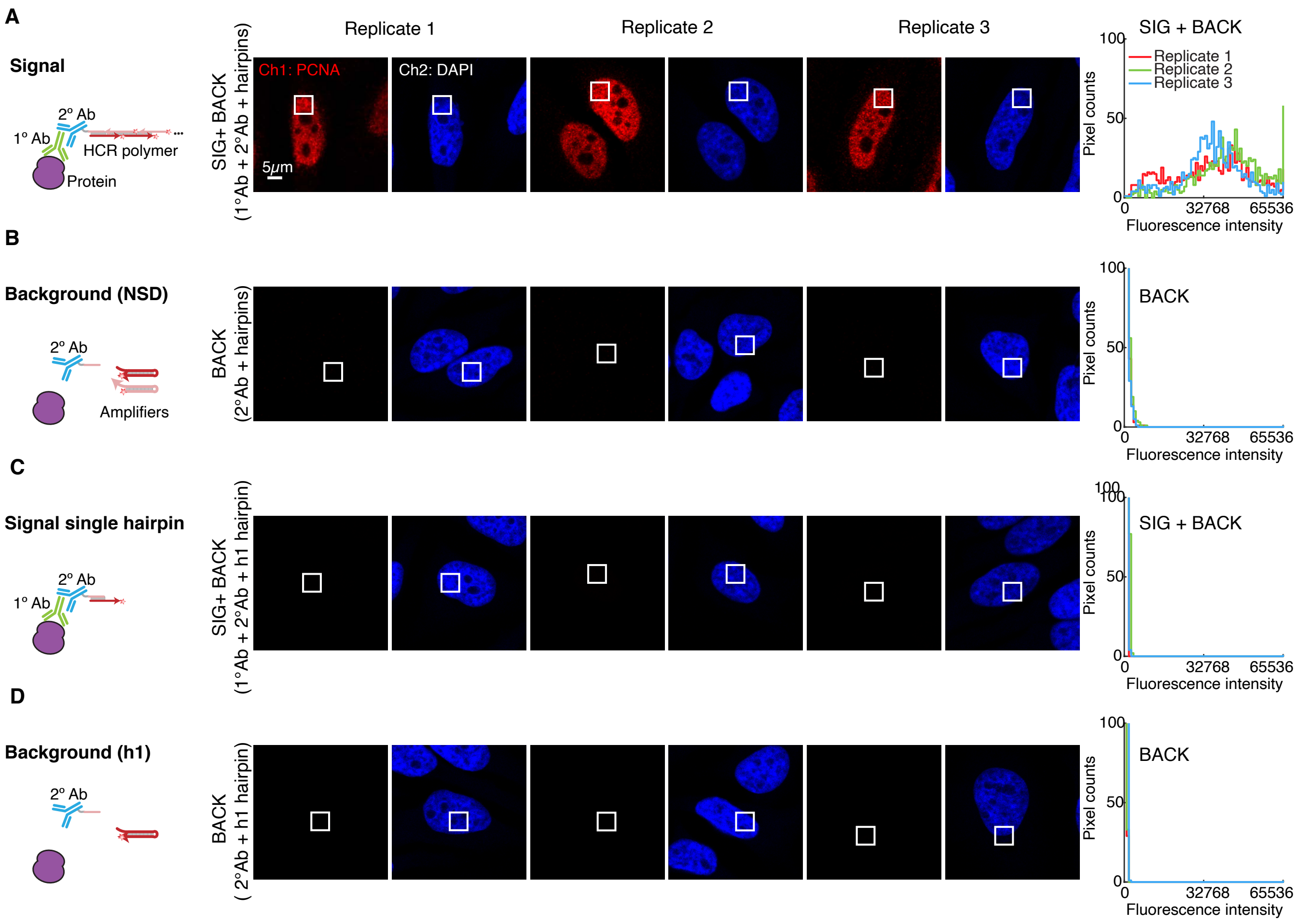

**Figure S21. Measurement of HCR amplification gain (mean polymer length) for HCR 2°IHC in mammalian cells on a slide (cf. Figure 3C).** (A,B) HCR 2°IHC. (C,D) HCR 2°IHC (h1 only). For each of 2 methods there are 2 rows: the top row measures SIG+BACK and the bottom row measures an approximation for BACK (see Table S19 for details on the background approximation used for each method). Left: Schematic of reagents used. Middle: confocal images collected with the microscope gain optimized to avoid saturating SIG+BACK pixels using HCR 2°IHC (panel A); DAPI channel facilitates placement of rectangles; single optical section. Pixel size: $0.198 \times 0.198$ $\mu$m. Right: pixel intensity histograms for representative regions (one rectangle in each of 5 individual cells in each of 3 replicate wells on a multi-well slide). Ch1: target protein PCNA (Alexa647). Ch2: DAPI. Sample: HeLa cells.

| | Quantity | Reagents | Value | | Figure |
|---|---|---|---|---|---|
| **A** | SIG+BACK | 1°Ab + 2°Ab-i1 + h1 + h2 | 37 000 | ± 2000 | S21A |
| | $NSD_{2^\circ}$+NSA+AF | 2°Ab-i1 + h1 + h2 | 460 | ± 20 | S21B |
| | SIG | | 36 000 | ± 2000 | |
| **B** | $SIG_{h1}$+$BACK_{h1}$ | 1°Ab + 2°Ab-i1 + h1 | 390 | ± 40 | S21C |
| | $NSD_{2^\circ h1}$+$NSA_{h1}$+AF | 2°Ab-i1 + h1 | 119 | ± 6 | S21D |
| | $SIG_{h1}$ | | 280 | ± 40 | |
| **C** | SIG/$SIG_{h1}$ | | 130 | ± 20 | |

**Table S19. Estimate of HCR amplification gain (mean polymer length) for HCR 2°IHC in mammalian cells on a slide (cf. Figure 3C).** (A) Estimated signal for HCR 2°IHC using HCR signal amplification (hairpins h1 and h2). The signal estimate SIG is calculated using the background approximation BACK $\approx$ $NSD_{2^\circ}$+NSA+AF. (B) Estimated signal $SIG_{h1}$ without HCR signal amplification (using only hairpin h1 so that HCR polymerization is not possible and only a single h1 hairpin can bind initiator i1). The signal estimate $SIG_{h1}$ is calculated using the background approximation BACK $\approx$ $NSD_{2^\circ h1}$+$NSA_{h1}$+AF. (C) Estimated HCR amplification gain SIG/$SIG_{h1}$ (i.e., mean HCR polymer length). Instrument noise is negligible using confocal microscopy so calculations use the approximation NOISE $\approx 0$. Mean $\pm$ standard error of the mean, $N = 15$ representative rectangular regions (one rectangle in each of 5 individual cells on each of 3 replicate wells on a multi-well slide). Analysis based on representative rectangular regions (examples depicted in Figures S21) using methods of Sections S2.6.2 and S2.6.4.

##### S5.5.3 HCR 1°IHC in FFPE mouse brain sections

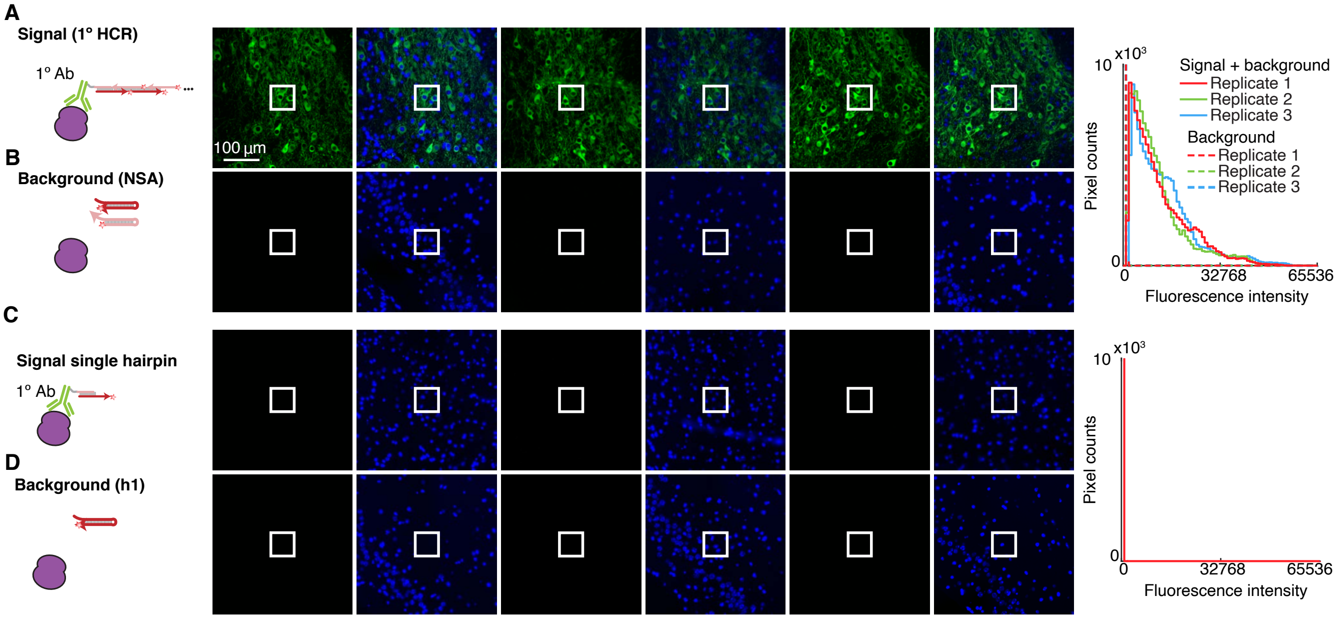

**Figure S22. Measurement of HCR amplification gain (mean polymer length) for HCR 1°IHC in FFPE mouse brain sections (cf. Figures 2DE).** (A,B) HCR 1°IHC. (C,D) HCR 1°IHC (h1 only). For each of 2 methods there are 2 rows: the top row measures SIG+BACK+NOISE in a region of high expression and the bottom row measures BACK+NOISE in a region of no/low expression. Left: Schematic of reagents used. Middle: epifluorescence images collected with the microscope exposure time optimized to avoid saturating SIG+BACK+NOISE pixels using HCR 1°IHC (panel A); DAPI channel facilitates placement of rectangles. Pixel size: 0.16×0.16 $\mu$m. Right: pixel intensity histograms for representative regions (three rectangles per experiment type for each of three replicate FFPE mouse brain sections). Ch1: target protein TH (Alexa647). Ch2: DAPI. Sample: FFPE C57BL/6 mouse brain section (coronal); thickness: 5 $\mu$m.

| | Quantity | Expression region | Value | | Figure |
|---|---|---|---|---|---|
| **A** | SIG+BACK | high | 12 000 | ± 1100 | S22A |
| | BACK | no/low | 70 | ± 30 | S22B |
| | SIG | | 11 900 | ± 1100 | |
| **B** | $\text{SIG}_{\text{h1}}$+BACK | high | 65 | ± 5 | S22C |
| | $\text{BACK}_{\text{h1}}$ | no/low | 14 | ± 1 | S22D |
| | $\text{SIG}_{\text{h1}}$ | | 51 | ± 4 | |
| **C** | SIG/$\text{SIG}_{\text{h1}}$ | | 230 | ± 30 | |

**Table S20. Estimate of HCR amplification gain (mean polymer length) for HCR 1°IHC in FFPE mouse brain sections (cf. Figure 2DE).** (A) Estimated signal for HCR 1°IHC using HCR signal amplification (hairpins h1 and h2). SIG+BACK characterized for rectangular regions of high expression; BACK characterized for rectangular regions of no/low expression. (B) Estimated signal $\text{SIG}_{\text{h1}}$ without HCR signal amplification (using only hairpin h1 so that HCR polymerization is not possible and only a single h1 hairpin can bind initiator i1). $\text{SIG}_{\text{h1}}$+$\text{BACK}_{\text{h1}}$ characterized for rectangular regions of high expression; $\text{BACK}_{\text{h1}}$ characterized for rectangular regions of no/low expression. (C) Estimated HCR amplification gain SIG/$\text{SIG}_{\text{h1}}$ (i.e., mean HCR polymer length). Mean ± standard error of the mean, $N = 3$ replicate FFPE mouse brain sections. Analysis based on representative rectangular regions (examples depicted in Figures S22) using methods of Sections S2.6.2 and S2.6.4.

##### S5.5.4 HCR 2°IHC in FFPE mouse brain sections

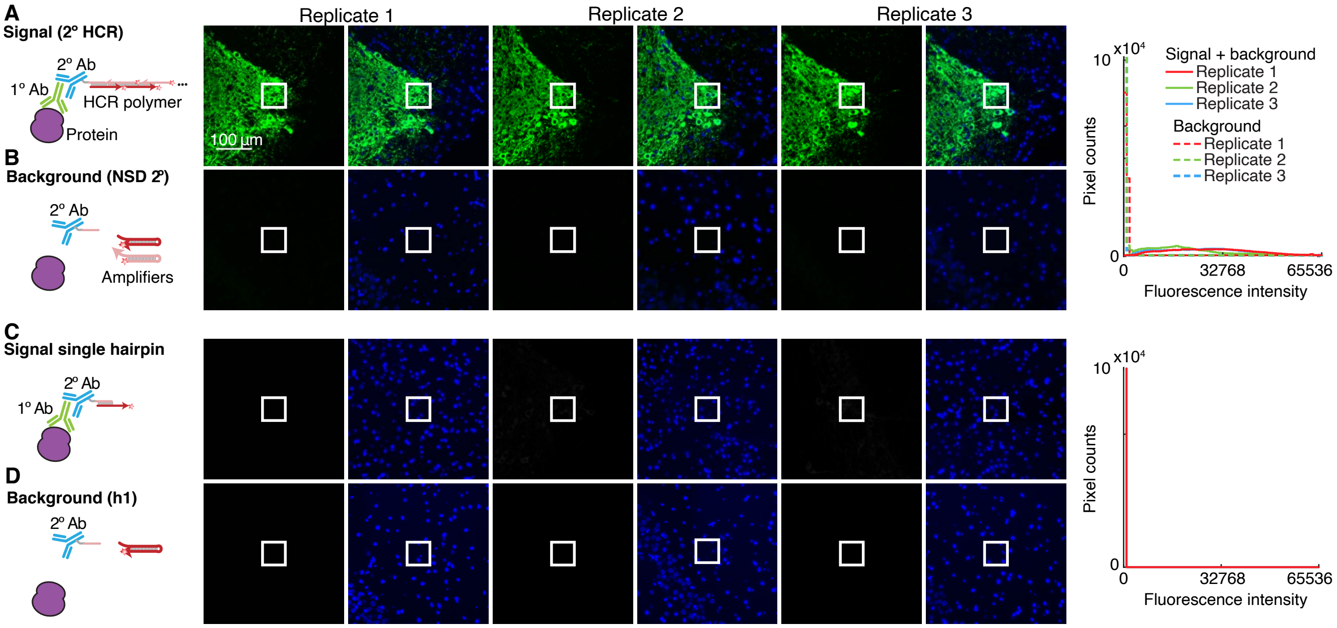

**Figure S23. Measurement of HCR amplification gain (mean polymer length) for HCR 2°IHC in FFPE mouse brain sections (cf. Figure 3DE).** (A,B) HCR 2°IHC. (C,D) HCR 2°IHC (h1 only). For each of 2 methods there are 2 rows: the top row measures SIG+BACK+NOISE in a region of high expression and the bottom row measures BACK+NOISE in a region of no/low expression. Left: Schematic of reagents used. Middle: epifluorescence images collected with the microscope exposure time optimized to avoid saturating SIG+BACK+NOISE pixels using HCR 1°IHC (panel A); DAPI channel facilitates placement of rectangles. Pixel size: 0.16×0.16 $\mu$m. Right: pixel intensity histograms for representative regions (three rectangles per experiment type for each of three replicate FFPE mouse brain sections). Ch1: target protein TH (Alexa647). Ch2: DAPI. Sample: FFPE C57BL/6 mouse brain section (coronal); thickness: 5 $\mu$m.

| | Quantity | Expression region | Value | | Figure |
|---|---|---|---|---|---|
| **A** | SIG+BACK | high | 24 000 | ± 3000 | S23A |
| | BACK | no/low | 170 | ± 90 | S23B |
| | SIG | | 24 000 | ± 3000 | |
| **B** | $SIG_{h1}$+$BACK_{h1}$ | high | 57 | ± 6 | S23C |
| | $BACK_{h1}$ | no/low | 4 | ± 3 | S23D |
| | $SIG_{h1}$ | | 53 | ± 4 | |
| **C** | SIG/$SIG_{h1}$ | | 450 | ± 50 | |

**Table S21. Estimate of HCR amplification gain (mean polymer length) for HCR 2°IHC in FFPE mouse brain sections (cf. Figure 3DE).** (A) Estimated signal for HCR 2°IHC using HCR signal amplification (hairpins h1 and h2). SIG+BACK characterized for rectangular regions of high expression; BACK characterized for rectangular regions of no/low expression. (B) Estimated signal $SIG_{h1}$ without HCR signal amplification (using only hairpin h1 so that HCR polymerization is not possible and only a single h1 hairpin can bind initiator i1). $SIG_{h1}$+$BACK_{h1}$ characterized for rectangular regions of high expression; $BACK_{h1}$ characterized for rectangular regions of no/low expression. (C) Estimated HCR amplification gain SIG/$SIG_{h1}$ (i.e., mean HCR polymer length). Mean ± standard error of the mean, $N = 3$ replicate FFPE mouse brain sections. Analysis based on representative rectangular regions (examples depicted in Figures S23) using methods of Sections S2.6.2 and S2.6.4.

#### S5.6 qHCR imaging: protein relative quantitation with subcellular resolution in an anatomical context (cf. Figure 4)

Additional studies are presented as follows:

- Section S5.6.1 presents a crowding study to test whether HCR amplification polymers for different targets interact within the cell.
- Section S5.6.2 provides replicates for redundant 2-channel imaging of target protein TH using HCR 1°IHC in FFPE mouse brain sections.
- Section S5.6.3 provides replicates for redundant 2-channel imaging of target proteins KRT17 and KRT19 using HCR 2°IHC in FFPE human breast tissue sections.

##### S5.6.1 Testing for a crowding effect

In order to perform multiplexed quantitative imaging using HCR, it is important that there is not a crowding effect in which amplification polymers tethered to one target molecule affect the signal intensity for a different target molecule. To test for a possible crowding effect, we imaged two target proteins (SC35 and PCNA) that are highly expressed in the nucleus individually (1-target studies) and also simultaneously (2-target studies) within HeLa cells. Figure S24 compares the signal intensity distributions for 1-target and 2-target studies, revealing similar intensity distributions whether targets were detected alone or together, suggesting that there is not a significant crowding effect (either antagonistic or synergistic). Intensities are plotted for subcellular $2{\times}2$ $\mu$m voxels that fall entirely within a cell nucleus as determined based on a DAPI mask.

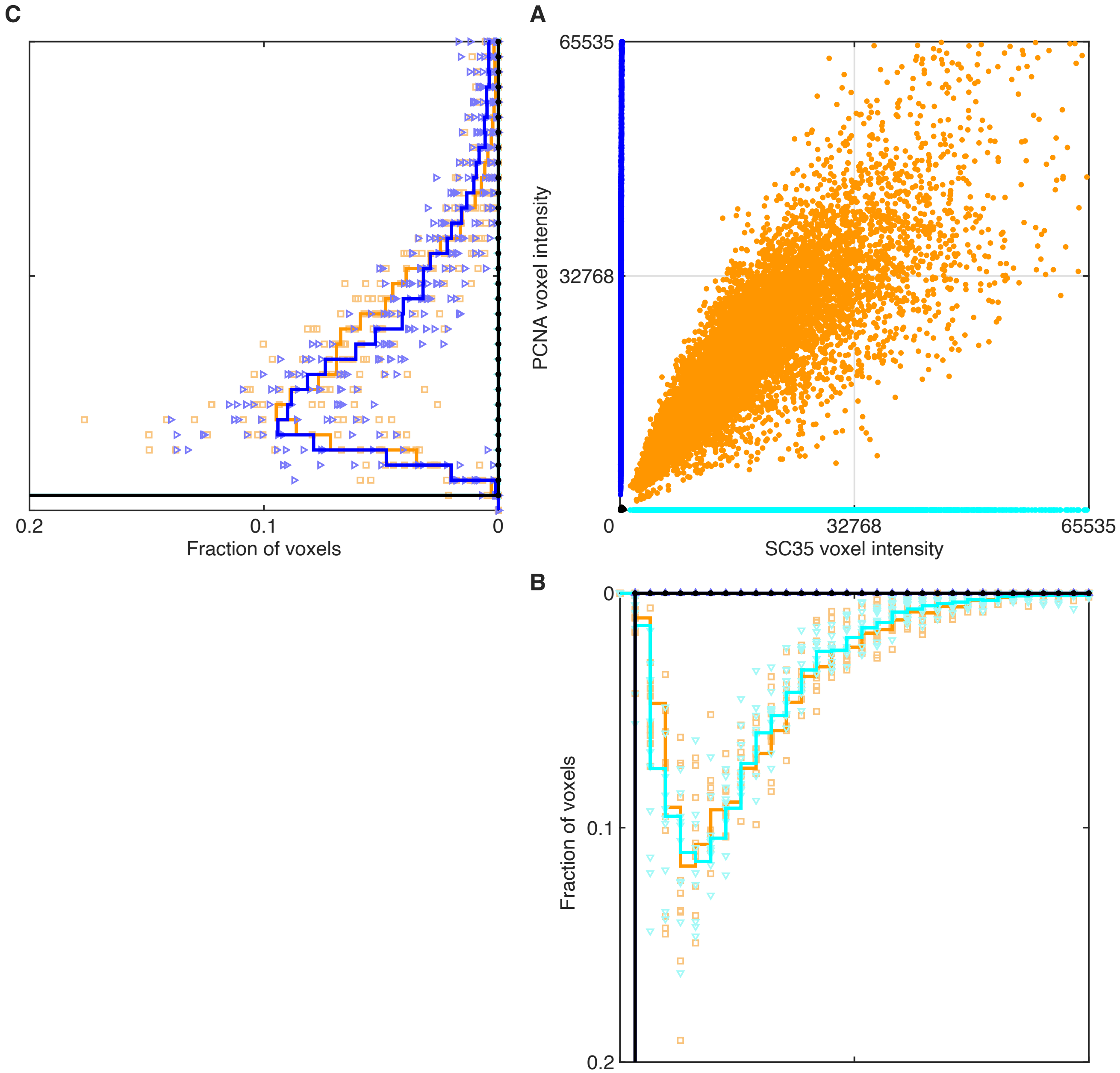

**Figure S24. Comparison of fluorescence intensity distributions for one-target and two-target experiments.** Detection of target proteins SC35 and PCNA with unlabeled primary antibodies and initiator-labeled secondary antibodies that trigger orthogonal spectrally-distinct HCR amplifiers. Ch1: Target protein SC35, probe 1°mAb mouse IgG1 anti-SC35, probe 2°mAb goat anti-mouse IgG1-B2, amplifier B2-Alexa546. Ch2: Target protein PCNA, probe 1°mAb mouse IgG2a anti-PCNA, probe 2°mAb goat anti-mouse IgG2a-B5, amplifier B5-Alexa647. (A) Raw voxel intensity scatter plot: SC35 vs PCNA. (B) Raw voxel intensity histogram for SC35. (C) Raw voxel intensity histogram for PCNA. In panels B and C, solid lines denote average histograms over cells in 10 replicate wells on a multi-well slide while symbols denote individual histograms (1 histogram per replicate well). Orange data: signal plus background for SC35 and PCNA (Figure S25). Cyan data: signal plus background for SC35 and background for PCNA (Figure S26). Blue data: background for SC35 and signal plus background for PCNA (Figure S27). Black data (near origin): background for SC35 and PCNA (Figure S28). Voxel size: 2.0×2.0 $\mu$m. Sample: HeLa cells.

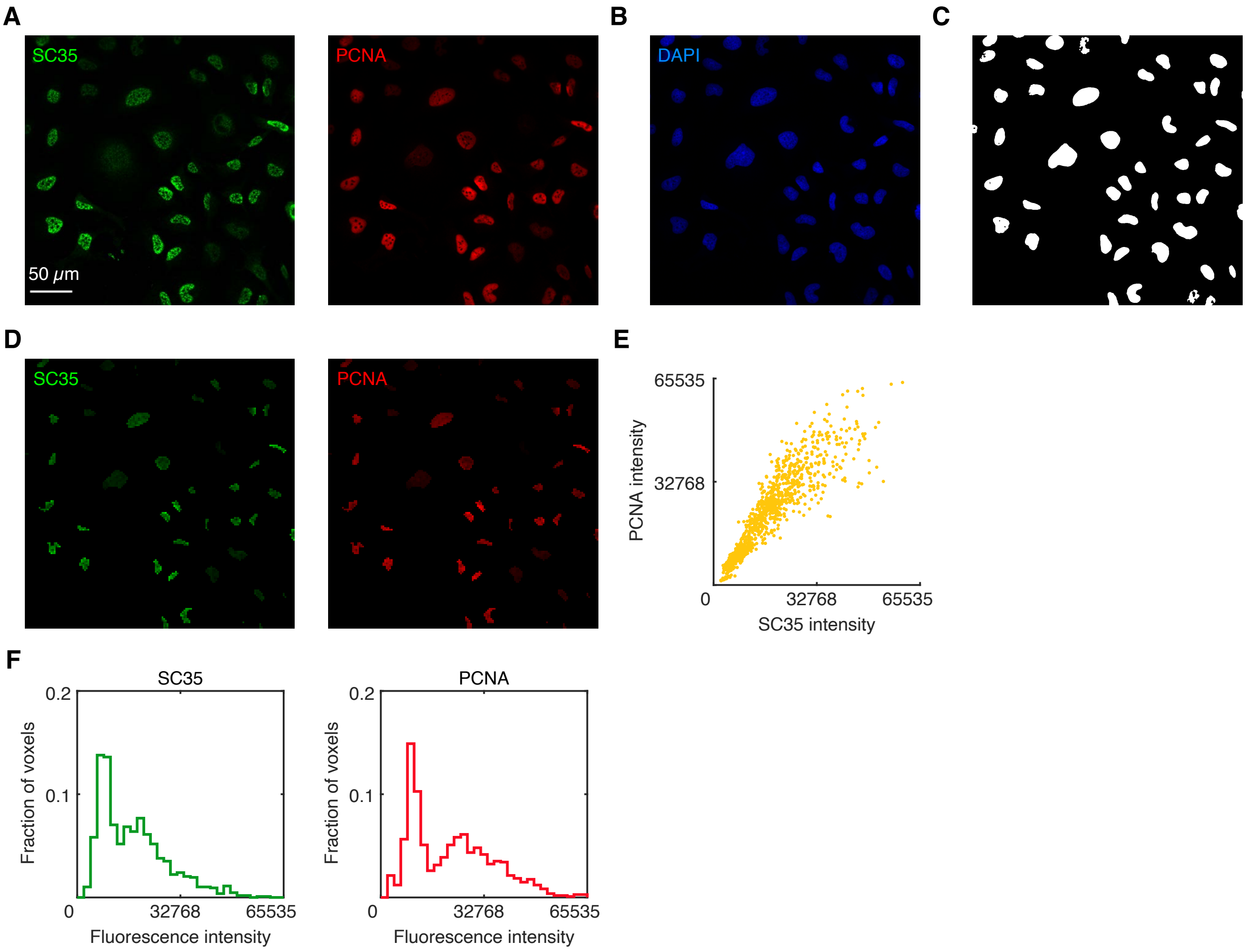

**Figure S25. Characterizing signal plus background for SC35 and PCNA in a 2-target experiment.** Detection of target proteins SC35 and PCNA with two unlabeled primary antibodies and two initiator-labeled secondary antibodies that trigger orthogonal spectrally-distinct HCR amplifiers. Ch1: Target protein SC35, probe 1°mAb mouse IgG1 anti-SC35, probe 2°mAb goat anti-mouse IgG1-B2, amplifier B2-Alexa546. Ch2: Target protein PCNA, probe 1°mAb mouse IgG2a anti-PCNA, probe 2°mAb goat anti-mouse IgG2a-B5, amplifier B5-Alexa647. (A) SC35 and PCNA channels from 3-channel confocal image; single optical section. Pixel size: $0.31\times0.31$ $\mu$m. (B) DAPI channel from 3-channel confocal image; single optical section. Pixel size: $0.31\times0.31$ $\mu$m. (C) Nuclear mask based on the DAPI staining of panel B; Gaussian blur filter followed by pixel thresholding. (D) Subcellular voxels falling entirely within the mask of panel C. Voxel size: $2.0\times2.0$ $\mu$m. (E) Raw voxel intensity scatter plots for the masked regions of panel D representing signal plus background for SC35 and PCNA. (F) Raw voxel intensity histograms for the masked regions of panel D representing signal plus background for SC35 and PCNA. Same microscope settings used for all replicates in Figures S25–S28. Sample: HeLa cells.

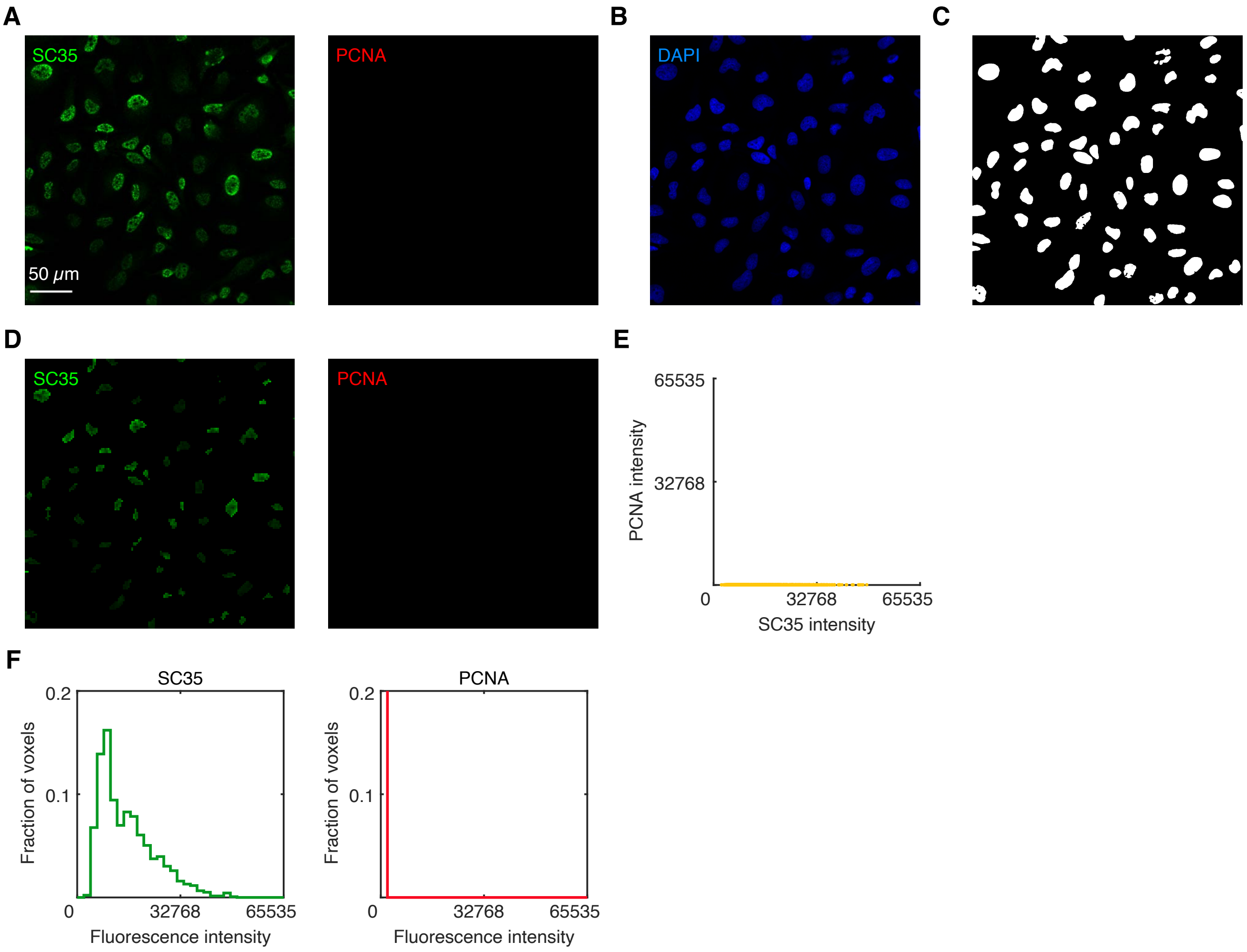

**Figure S26. Characterizing signal plus background for SC35 in a 1-target experiment.** Detection of target protein SC35 with an unlabeled primary antibody probe and initiator-labeled secondary antibody probe. Ch1: Target protein SC35, probe 1°mAb mouse IgG1 anti-SC35, probe 2°mAb goat anti-mouse IgG1-B2, amplifier B2-Alexa546. Ch2: no probe, no amplifier. (A) SC35 and PCNA channels from 3-channel confocal image; single optical section. Pixel size: $0.31 \times 0.31$ $\mu$m. (B) DAPI channel from 3-channel confocal image; single optical section. Pixel size: $0.31 \times 0.31$ $\mu$m. (C) Nuclear mask based on the DAPI staining of panel B; Gaussian blur filter followed by pixel thresholding. (D) Subcellular voxels falling entirely within the mask of panel C. Voxel size: $2.0 \times 2.0$ $\mu$m. (E) Raw voxel intensity scatter plots for the masked regions of panel D representing signal plus background for SC35 and background for PCNA. (F) Raw voxel intensity histograms for the masked regions of panel D representing signal plus background for SC35 and background for PCNA. Same microscope settings used for all replicates in Figures S25–S28. Sample: HeLa cells.

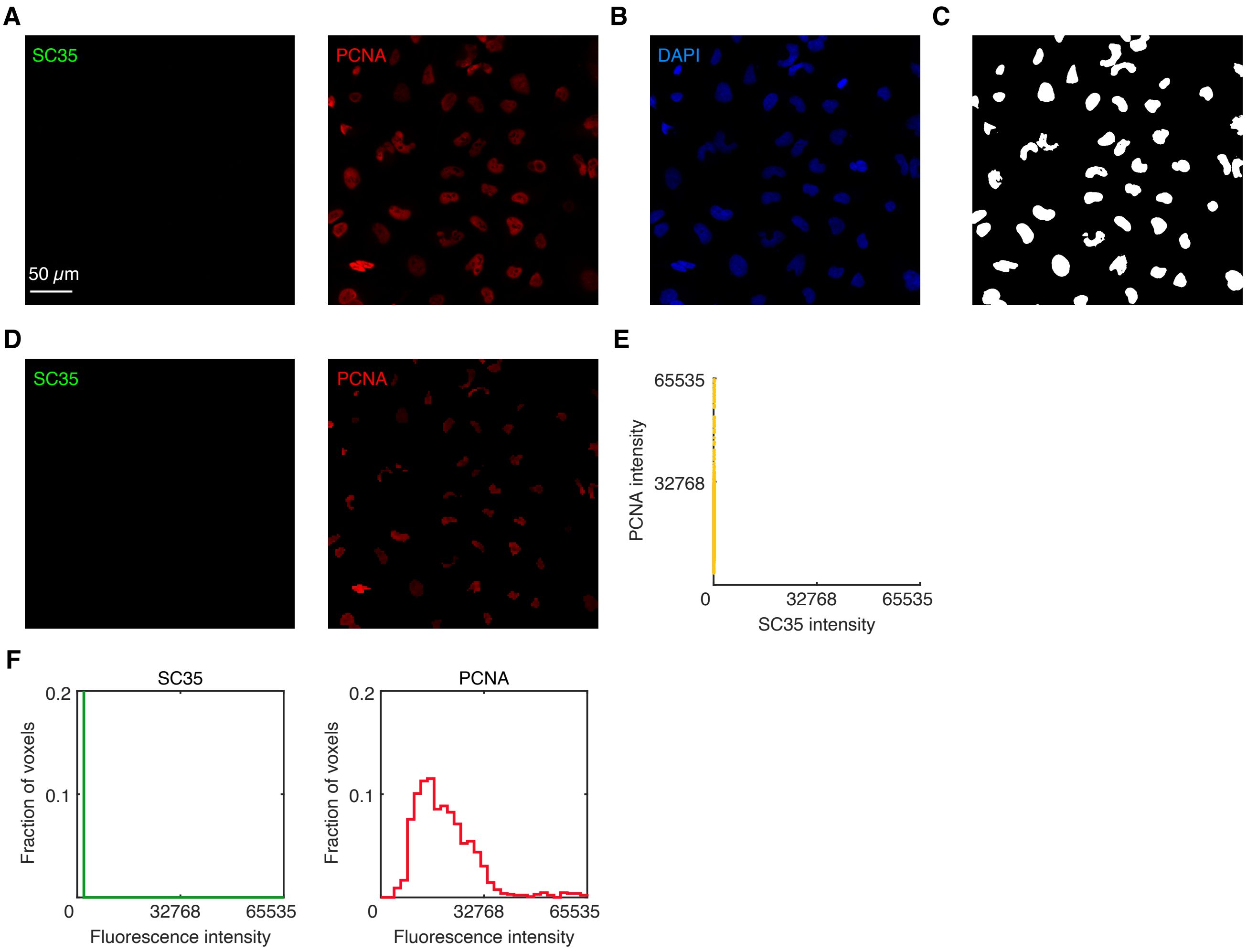

**Figure S27. Characterizing signal plus background for PCNA in a 1-target experiment.** Detection of target protein PCNA with an unlabeled primary antibody probe and initiator-labeled secondary antibody probe. Ch1: no probe, no amplifier. Ch2: Target protein PCNA, probe 1°mAb mouse IgG2a anti-PCNA, probe 2°mAb goat anti-mouse IgG2a-B5, amplifier B5-Alexa647. (A) SC35 and PCNA channels from 3-channel confocal image; single optical section. Pixel size: $0.31 \times 0.31$ $\mu$m. (B) DAPI channel from 3-channel confocal image; single optical section. Pixel size: $0.31 \times 0.31$ $\mu$m. (C) Nuclear mask based on the DAPI staining of panel B; Gaussian blur filter followed by pixel thresholding. (D) Subcellular voxels falling entirely within the mask of panel C. Voxel size: $2.0 \times 2.0$ $\mu$m. (E) Raw voxel intensity scatter plots for the masked regions of panel D representing background for SC35 and signal plus background for PCNA. (F) Raw voxel intensity histograms for the masked regions of panel D representing background for SC35 and signal plus background for PCNA. Same microscope settings used for all replicates in Figures S25–S28. Sample: HeLa cells.

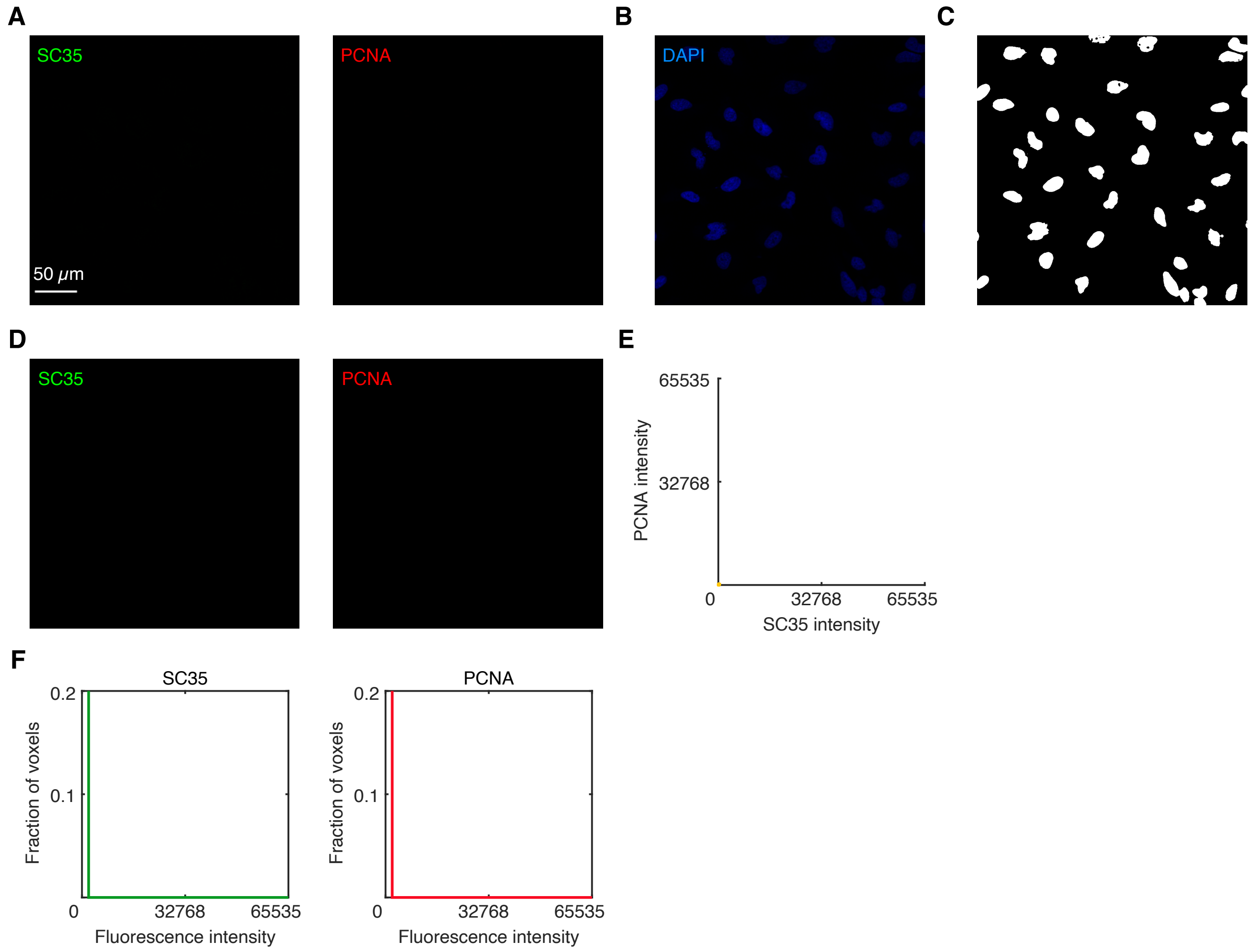

**Figure S28. Characterizing background for SC35 and PCNA.** Background is estimated using the standard HCR 2°IHC protocol omitting probes (BACK ≈ $NSD_{2°}$+NSA+AF+NOISE; see Section S2.6 for definitions). (A) SC35 and PCNA channels from 3-channel confocal image; single optical section. Pixel size: 0.31×0.31 μm. (B) DAPI channel from 3-channel confocal image; single optical section. Pixel size: 0.31×0.31 μm. (C) Nuclear mask based on the DAPI staining of panel B; Gaussian blur filter followed by pixel thresholding. (D) Subcellular voxels falling entirely within the mask of panel C. Voxel size: 2.0×2.0 μm. (E) Raw voxel intensity scatter plots for the masked regions of panel D representing background for SC35 and PCNA. (F) Raw voxel intensity histograms for the masked regions of panel D representing background for SC35 and PCNA. Same microscope settings used for all replicates in Figures S25–S28. Sample: HeLa cells.

##### S5.6.2 Redundant 2-channel imaging of target protein TH using HCR 1°IHC in FFPE mouse brain sections

Here, we perform redundant 2-channel imaging of target protein TH using HCR 1°IHC in FFPE mouse brain sections. The target is detected with two initiator-labeled primary antibodies that bind different epitopes (e1 and e2) on the target protein and trigger orthogonal spectrally-distinct HCR amplifiers. The reagents for this 2-channel experiment are:

- **Ch1:** Target protein TH, probe 1°$mAb_{e1}$ EP1533Y anti-TH labeled with B1 initiator, amplifier B1-Alexa647.
- **Ch2:** Target protein TH, probe 1°$mAb_{e2}$ EP1532Y anti-TH labeled with B3 initiator, amplifier B3-Alexa750.

Additional studies are presented as follows:

- Figure S29 displays 2-plex images and 2-channel voxel intensity scatter plots for $N = 3$ replicate FFPE mouse brain sections.
- Table S22 displays estimated values for signal, background, and signal-to-background for each channel.

**Protocol:** HCR 1°IHC (Section S3.2) using initiator-labeled primary antibody probes and HCR signal amplification.
**Sample:** FFPE C57BL/6 mouse brain section (coronal); thickness: 5 $\mu$m.
**Microscopy:** Epifluorescence.

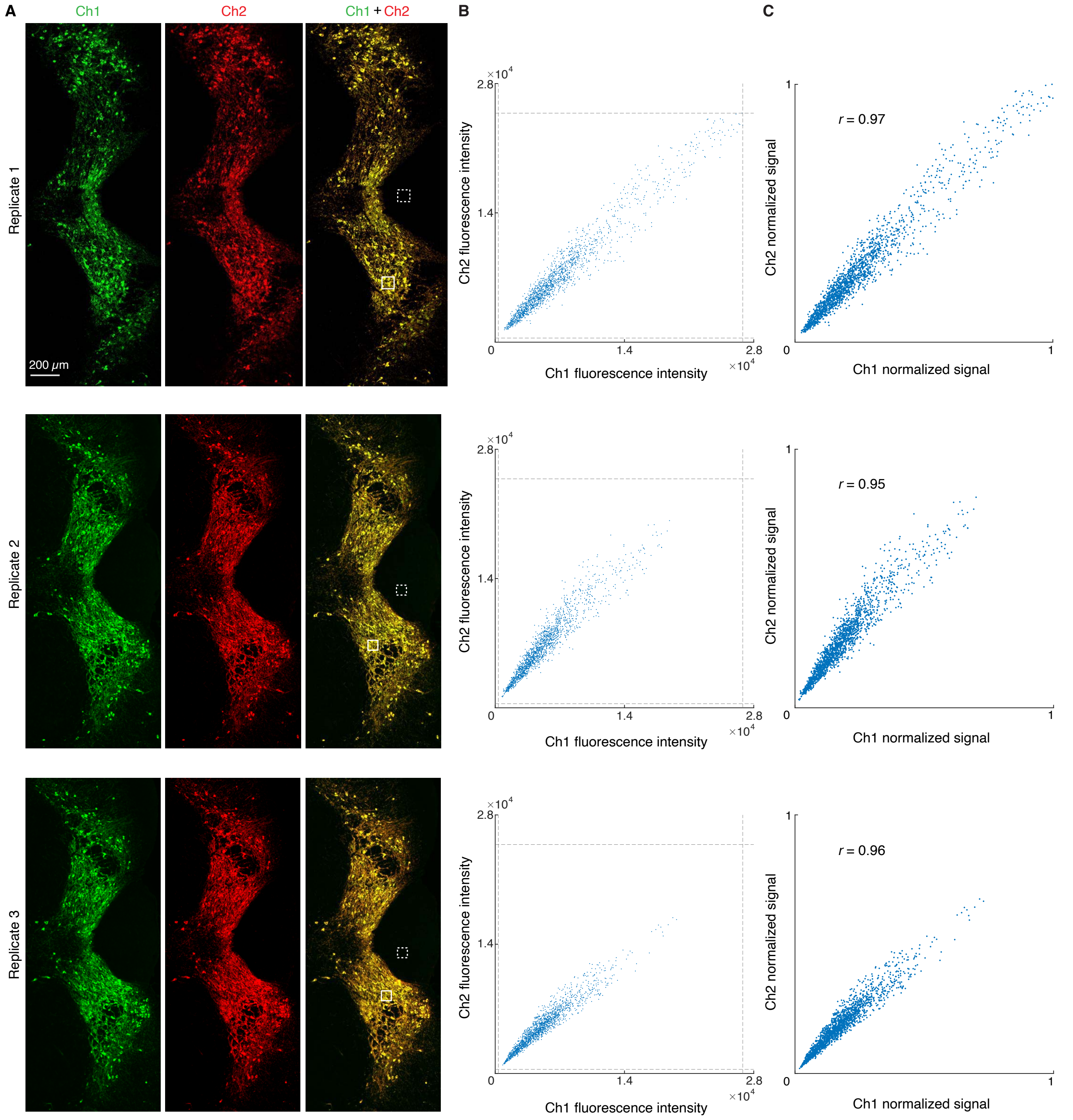

**Figure S29. Redundant 2-channel detection of target protein TH in FFPE mouse brain sections (cf. Figure 4).** (A) Epifluorescence images: individual channels and merge. Solid boundaries denote representative regions of high expression; dashed boundaries denote representative regions of no/low expression. Pixel size: 0.2×0.2 $\mu$m. (B) Raw voxel intensity scatter plots representing signal plus background plus noise for voxels within solid boundaries of panel A. Voxel size: 2.0×2.0 $\mu$m. Dashed lines represent BOT and TOP values (Table S22) used to normalize data for panel C using methods of Section S2.6.5. (C) Normalized voxel intensity scatter plots representing estimated normalized signal (Pearson correlation coefficient, $r$). Ch1: target protein TH (Alexa 546). Ch2: target protein TH (Alexa647). Sample: FFPE C57BL/6 mouse brain section (coronal); thickness: 5 $\mu$m.

| Channel | Target protein | Fluorophore | BACK+NOISE | SIG+BACK+NOISE | SIG | SIG/BACK | BOT | TOP |
|---|---|---|---|---|---|---|---|---|
| Ch1 | TH | Alexa647 | $253 \pm 8$ | $5600 \pm 700$ | $5400 \pm 700$ | $45 \pm 9$ | 253 | 26800 |
| Ch2 | TH | Alexa750 | $270 \pm 20$ | $6000 \pm 1000$ | $6000 \pm 1000$ | $41 \pm 9$ | 270 | 25000 |

**Table S22. Estimated signal-to-background for redundant 2-channel detection of target protein TH in FFPE mouse brain sections (cf. Figure 4).** Mean $\pm$ standard error, $N = 3$ replicate embryos. Analysis based on rectangular regions depicted in Figure S29A using methods of Section S2.6.2. BOT and TOP values used to calculate normalized voxel intensities for scatter plots of Figures 4C and S29C using methods of Section S2.6.5.

##### S5.6.3 Redundant 2-channel imaging of target proteins KRT17 and KRT19 using HCR 2°IHC in FFPE human breast tissue sections

Here, we perform redundant 2-channel imaging of target proteins KRT17 and KRT19 using HCR 2°IHC in FFPE human breast tissue sections. In each case, the target protein is detected with a 1°Ab probe, which is then redundantly detected by two batches of 2°Ab probes labeled with different HCR initiators that trigger orthogonal spectrally distinct HCR amplifiers. For redundant 2-channel imaging of KRT17, the reagents are:

- **Ch1:** Target protein KRT17, probe 1°pAb rabbit IgG anti-KRT17, probe 2°pAb donkey anti-rabbit labeled with B4 initiator, amplifier B4-Alexa546.
- **Ch2:** Target protein KRT17, probe 1°pAb rabbit IgG anti-KRT17, probe 2°pAb donkey anti-rabbit labeled with B3 initiator, amplifier B3-Alexa647.

For redundant 2-channel imaging of KRT19, the reagents are:

- **Ch1:** Target protein KRT19, probe 1°mAb mouse IgG1 anti-KRT19, probe 2°pAb goat anti-mouse IgG1 labeled with B2 initiator, amplifier B2-Alexa546.
- **Ch2:** Target protein KRT19, probe 1°mAb mouse IgG1 anti-KRT19, probe 2°pAb goat anti-mouse IgG1 labeled with B5 initiator, amplifier B5-Alexa647.

Additional studies are presented as follows:

- Figure S30 displays 2-plex images and 2-channel voxel intensity scatter plots for target protein KRT17 for $N = 3$ replicate FFPE human breast tissue sections.
- Figure S31 displays 2-plex images and 2-channel voxel intensity scatter plots for target protein KRT19 for $N = 3$ replicate FFPE human breast tissue sections.
- Table S23 displays values used for signal normalization in Figures S30 and S31.
- Figure S32 display representative regions used for measurement of signal and background for protein targets KRT17 and KRT19.
- Table S24 displays estimated values for signal, background, and signal-to-background for each channel for both KRT17 and KRT19.

**Protocol:** HCR 2°IHC (Section S4.3) using unlabeled primary antibody probes and initiator-labeled secondary antibody probes with HCR signal amplification.
**Sample:** FFPE human breast tissue section; thickness: 5 $\mu$m.
**Microscopy:** Confocal.

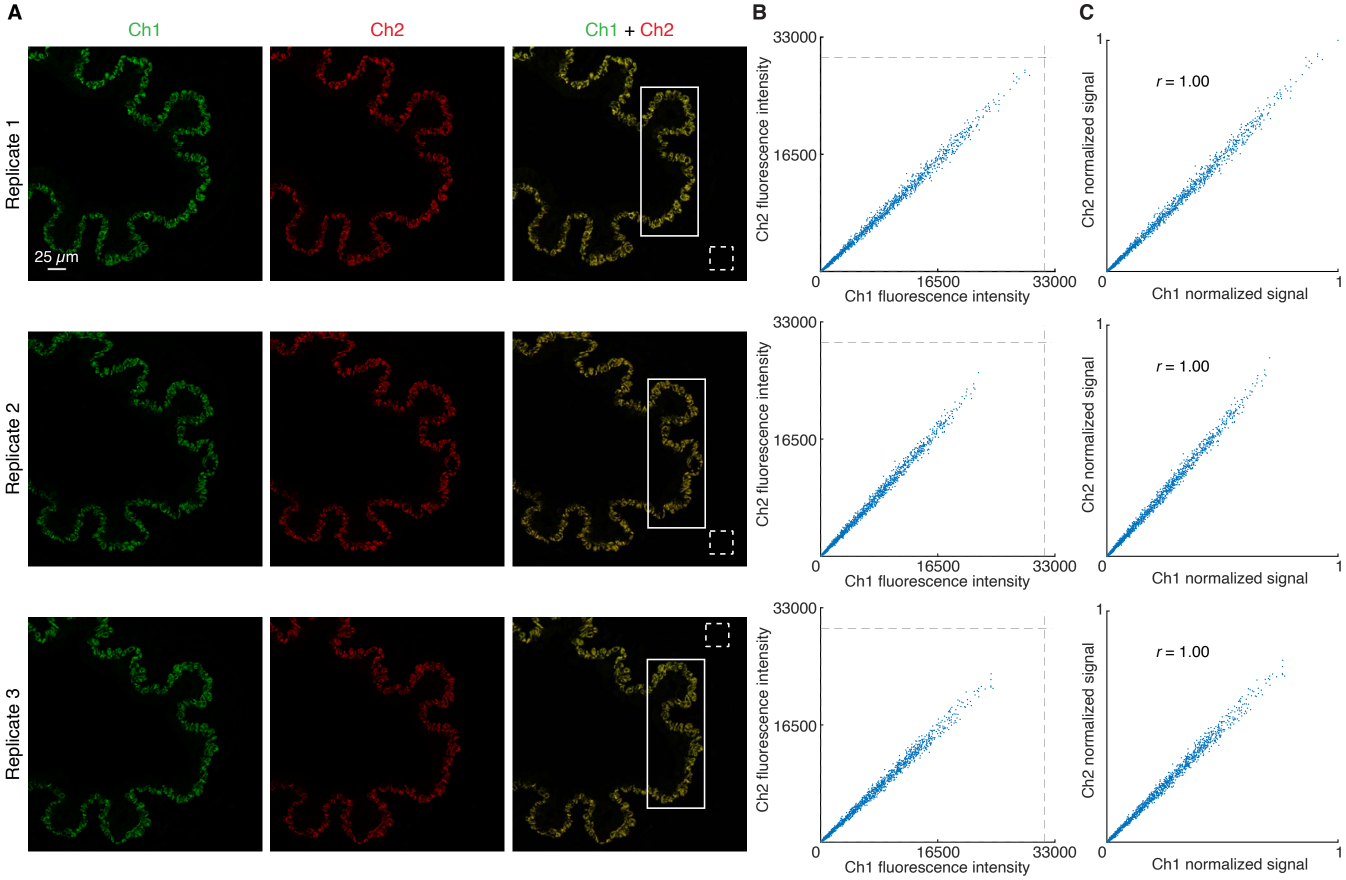

**Figure S30. Redundant 2-channel detection of target protein KRT17 in FFPE human breast tissue sections (cf. Figure 4).** (A) Confocal images: individual channels and merge; single optical section. Solid boundaries denote regions of variable expression; dashed boundaries denote regions of no/low expression. Pixel size: $0.312\times0.312$ $\mu$m. Sample: FFPE human breast tissue section; thickness: 5 $\mu$m. (B) Raw voxel intensity scatter plots representing signal plus background for voxels within solid boundaries of panel A. Voxel size: $2.0\times2.0$ $\mu$m. Dashed lines represent BOT and TOP values (Table S23) used to normalize data for panel C using methods of Section S2.6.5. (C) Normalized voxel intensity scatter plots representing estimated normalized signal (Pearson correlation coefficient, $r$). Ch1: KRT17 (Alexa546). Ch2: KRT17 (Alexa647). Sample: FFPE human breast tissue section; thickness: 5 $\mu$m.

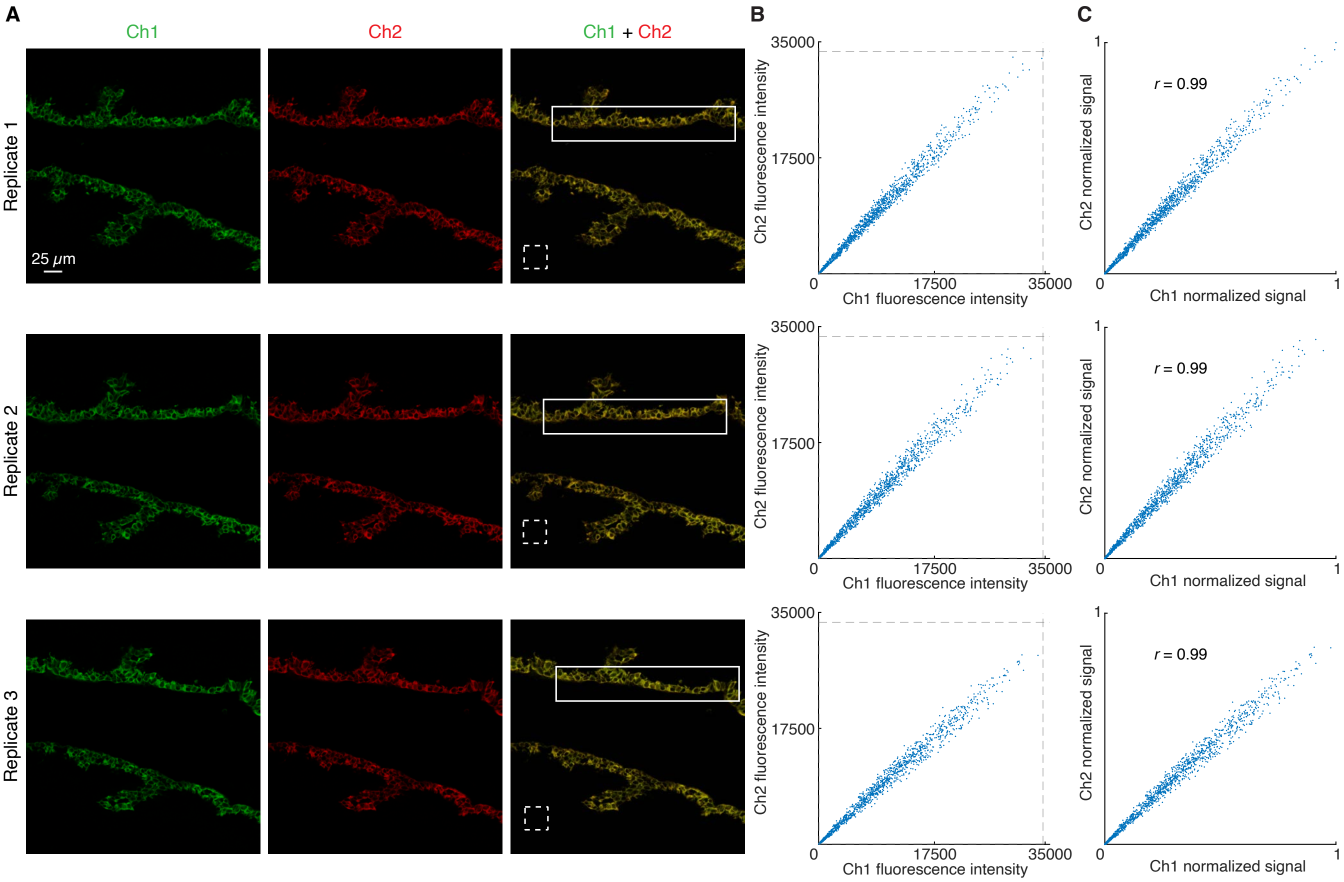

**Figure S31. Redundant 2-channel detection of target protein KRT19 in FFPE human breast tissue sections (cf. Figure 4).** (A) Confocal images: individual channels and merge; single optical section. Solid boundaries denote regions of variable expression; dashed boundaries denote regions of no/low expression. Pixel size: $0.312 \times 0.312$ $\mu$m. Sample: FFPE human breast tissue section. Thickness: 5 $\mu$m. (B) Raw voxel intensity scatter plots representing signal plus background for voxels within solid boundaries of panel A. Voxel size: $2.0 \times 2.0$ $\mu$m. Dashed lines represent BOT and TOP values (Table S23) used to normalize data for panel C using methods of Section S2.6.5. (C) Normalized voxel intensity scatter plots representing estimated normalized signal (Pearson correlation coefficient, $r$). Ch1: KRT19 (Alexa546). Ch2: KRT19 (Alexa647). Sample: FFPE human breast tissue section; thickness: 5 $\mu$m.

| Channel | Target protein | Fluorophore | BOT | TOP |
|---|---|---|---|---|
| Ch1 | KRT17 | Alexa546 | 70 | 31500 |
| Ch2 | KRT17 | Alexa647 | 70 | 30100 |
| Ch1 | KRT19 | Alexa546 | 70 | 34000 |
| Ch2 | KRT19 | Alexa647 | 63 | 33500 |

**Table S23. BOT and TOP values used to calculate normalized voxel intensities for scatter plots of Figures 4C, S30C, and S31C using methods of Section S2.6.5** Analysis based on rectangular regions depicted in Figures S30A and S31A.

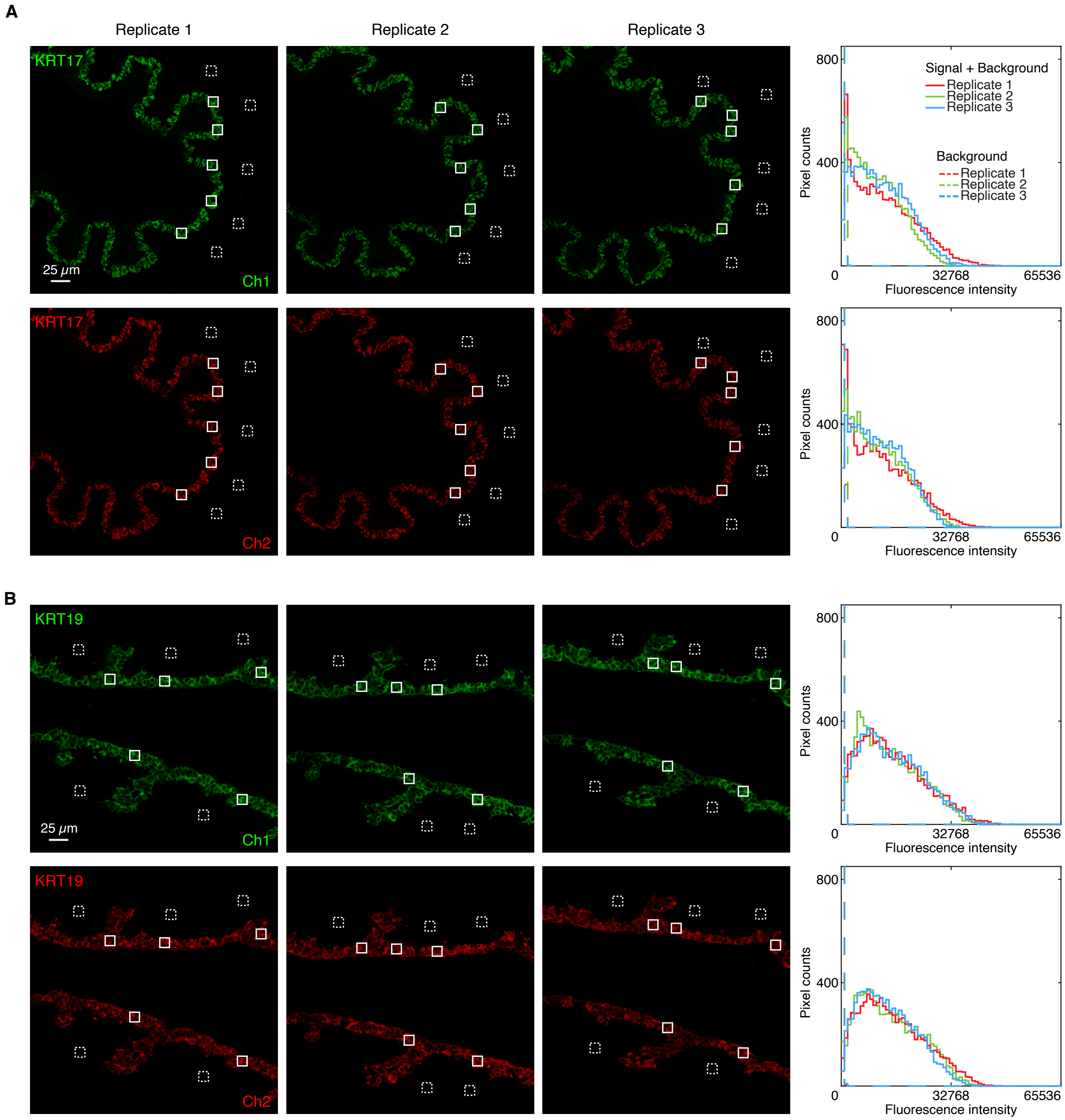

**Figure S32. Measurement of signal and background for redundant 2-channel detection of target proteins KRT17 and KRT19 in FFPE human breast tissue sections (cf. Figure 4).** (A) Redundant 2-channel imaging of target protein KRT17. Ch1: Alexa 546. Ch2: Alexa 647. (B) Redundant 2-channel imaging of target protein KRT19. Ch1: Alexa546. Ch2: Alexa647. Left: confocal images; single optical section. Solid boundaries denote representative regions of high expression; dashed boundaries denote representative regions of no/low expression. Right: pixel intensity histograms for the depicted representative regions. Sample: FFPE human breast tissue section; thickness: 5 $\mu$m.

| Channel | Target protein | Fluorophore | BACK | SIG+BACK | SIG | SIG/BACK |
|---|---|---|---|---|---|---|
| Ch1 | KRT17 | Alexa546 | $110 \pm 30$ | $11\,200 \pm 600$ | $11\,000 \pm 600$ | $100 \pm 30$ |
| Ch2 | KRT17 | Alexa647 | $100 \pm 20$ | $11\,000 \pm 500$ | $10\,900 \pm 500$ | $110 \pm 30$ |
| Ch1 | KRT19 | Alexa546 | $99 \pm 6$ | $14\,000 \pm 1000$ | $14\,000 \pm 1000$ | $140 \pm 10$ |
| Ch2 | KRT19 | Alexa647 | $83 \pm 4$ | $14\,200 \pm 900$ | $14\,100 \pm 900$ | $170 \pm 10$ |

**Table S24. Estimated signal-to-background for redundant 2-channel detection of target proteins KRT17 and KRT19 in FFPE human breast tissue sections (cf. Figure 4).** Instrument noise is negligible using confocal microscopy so calculations use the approximation NOISE $\approx 0$. Mean $\pm$ standard error, $N = 3$ replicate human breast tissue sections. Analysis based on rectangular regions depicted in Figure S32 using methods of Section S2.6.2.

#### S5.7 Replicates and signal-to-background measurements for simultaneous multiplexed protein and mRNA imaging using HCR 1°IHC and HCR RNA-ISH (cf. Figure 5)

##### S5.7.1 Mammalian cells on a slide

For 4-plex simultaneous protein and mRNA imaging using HCR 1°IHC + HCR RNA-ISH in mammalian cells on a slide, the 5 channels are (2 RNAs + 2 proteins + DAPI):

- **Ch1:** Target protein PCNA, probe 1°mAb mouse IgG2a anti-PCNA labeled with B5 initiator, amplifier B5-Alexa488.
- **Ch2:** Target protein HSP60, probe 1°mAb rabbit IgG anti-HSP60 labeled with B3 initiator, amplifier B3-Alexa546.
- **Ch3:** Target RNA *U6*, probe set with 2 split-initiator probe pairs, amplifier B1-Alexa594.
- **Ch4:** Target mRNA *ACTB*, probe set with 10 split-initiator pairs, amplifier B2-Alexa647.
- **Ch5:** DAPI.

Additional studies are presented as follows:

- Figure S33 displays 4-plex images for $N = 3$ replicate wells on a multi-well slide (cf. Figure 5B).
- Figure S34 displays representative regions of individual channels used for measurement of signal and background for each target.
- Table S25 displays estimated values for signal, background, and signal-to-background for each target.

**Protocol:** Simultaneous HCR 1°IHC + HCR RNA-ISH (Section S3.1) using initiator-labeled primary antibody probes for protein targets, split-initiator DNA probes for RNA targets, and simultaneous HCR signal amplification for all targets.
**Sample:** HeLa cells.
**Microscopy:** Confocal.

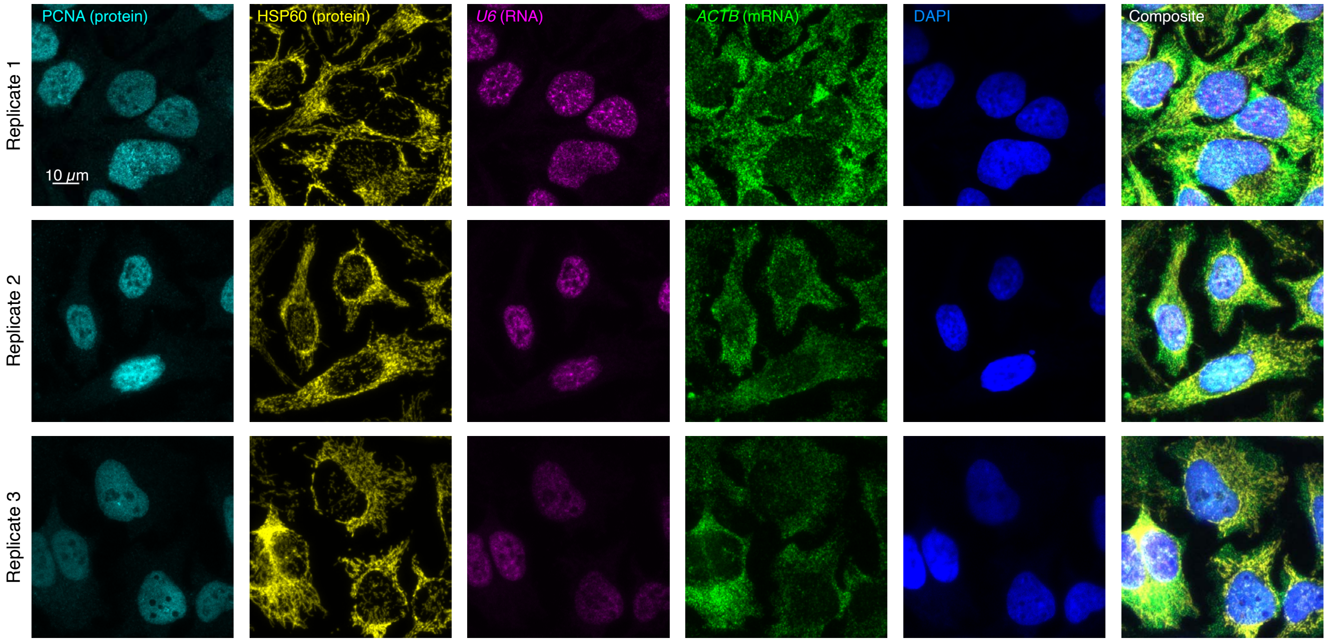

**Figure S33. Replicates for 4-plex simultaneous protein and mRNA imaging using HCR 1°IHC and HCR RNA-ISH in mammalian cells on slides (cf. Figures 5B).** 5-channel confocal images for 3 replicate wells on a multi-well slide; maximum intensity z-projection. Ch1: target protein PCNA (Alexa488). Ch2: target protein HSP60 (Alexa546). Ch3: target RNA *U6* (Alexa594). Ch4: target mRNA *ACTB* (Alexa647). Ch5: DAPI. Sample: HeLa cells.

| Channel | Target | Type | Fluorophore | BACK | SIG+BACK | SIG | SIG/BACK |
|---|---|---|---|---|---|---|---|
| Ch1 | PCNA | protein | Alexa488 | $276 \pm 4$ | $26\,600 \pm 1700$ | $26\,400 \pm 1700$ | $95 \pm 6$ |
| Ch2 | HSP60 | protein | Alexa546 | $134.4 \pm 1.3$ | $37\,100 \pm 1300$ | $37\,000 \pm 1300$ | $280 \pm 10$ |
| Ch3 | *U6* | RNA | Alexa594 | $100.9 \pm 0.7$ | $2150 \pm 130$ | $2050 \pm 130$ | $20.3 \pm 1.3$ |
| Ch4 | *ACTB* | mRNA | Alexa647 | $182 \pm 8$ | $20\,000 \pm 2000$ | $19\,000 \pm 2000$ | $107 \pm 12$ |

**Table S25. Estimated signal-to-background for 4-plex simultaneous protein and mRNA imaging using HCR 1°IHC and HCR RNA-ISH in mammalian cells on a slide (cf. Figure 5B).** The signal estimate SIG is calculated using the background approximation BACK $\approx$ NSA+AF. Instrument noise is negligible using confocal microscopy so calculations use the approximation NOISE $\approx 0$. Mean $\pm$ standard error of the mean, $N = 15$ representative rectangular regions (one rectangle in each of 5 individual cells in each of 3 replicate wells on a multi-well slide). Analysis based on representative rectangular regions (examples depicted in Figure S34) using methods of Section S2.6.2.

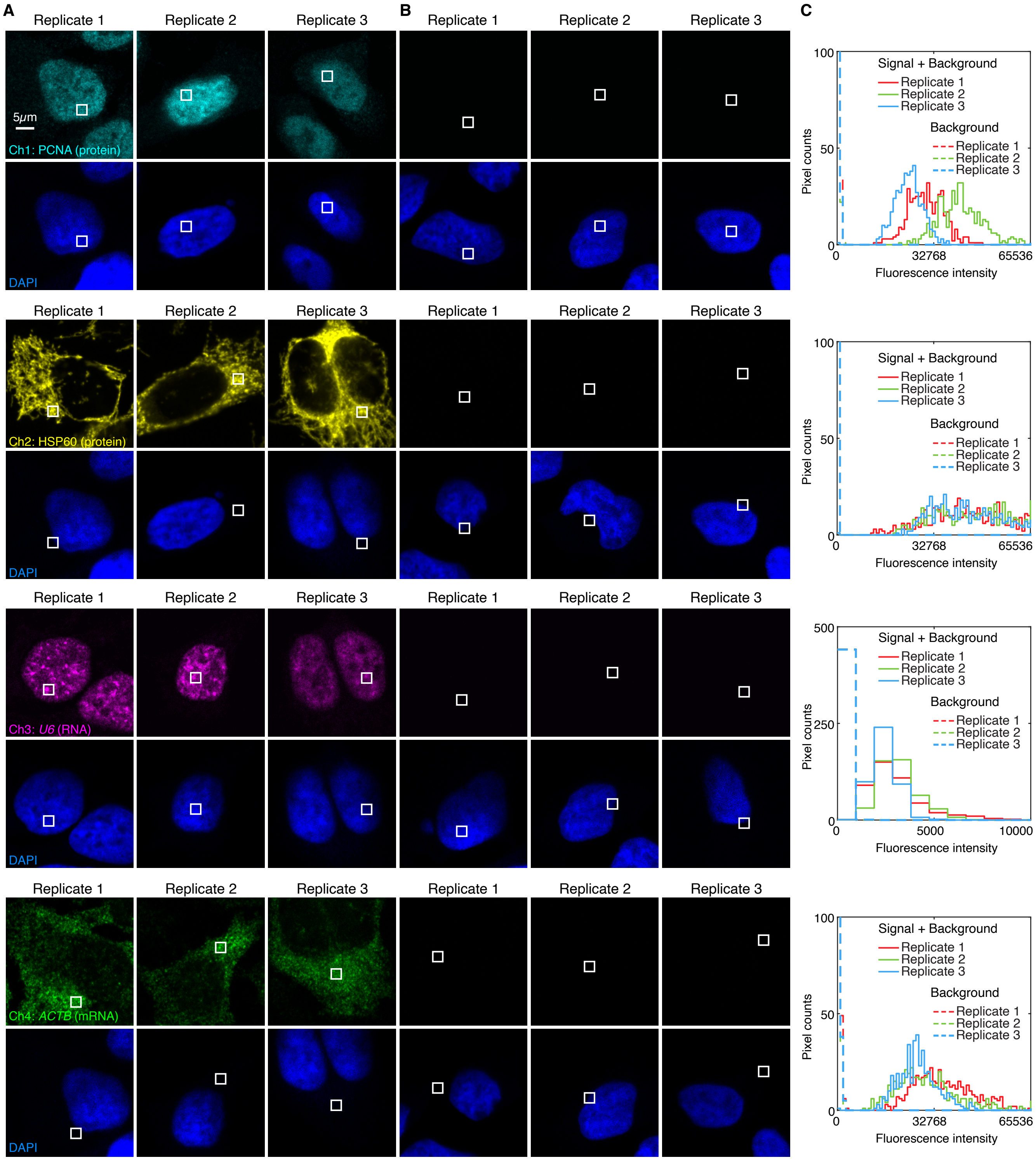

**Figure S34. Measurement of signal and background for 4-plex simultaneous protein and mRNA imaging using HCR 1°IHC and HCR RNA-ISH in mammalian cells on a slide (cf. Figure 5B).** (A) Use experiment of Type 1 in Table S7A (1°Ab probe + hairpins) to measure SIG+BACK in a region of high expression. (B) Use experiment of Type 2 in Table S7B (hairpins only) to measure NSA+AF in a region of maximum background. (C) Pixel intensity histograms for representative regions (one rectangle in each of 5 individual cells in each of 3 replicate wells on a multi-well slide). Confocal images collected with the microscope gain optimized to avoid saturating SIG+BACK pixels; DAPI channel facilitates placement of representative rectangles; single optical section. Ch1: target protein PCNA (Alexa488). Ch2: target protein HSP60 (Alexa546). Ch3: target RNA *U6* (Alexa594). Ch4: target mRNA *ACTB* (Alexa647). Ch5: DAPI. Sample: HeLa cells.

##### S5.7.2 FFPE mouse brain sections

For 4-plex simultaneous protein and mRNA imaging using HCR 1°IHC + HCR RNA-ISH in FFPE mouse brain sections, the 5 channels are (2 proteins + 2 RNAs + DAPI):

- **Ch1:** Target protein TH, probe 1°Ab rabbit IgG monoclonal anti-TH labeled with B1 initiator, amplifier B1-Alexa488.
- **Ch2:** Target protein MBP, probe 1°Ab rabbit IgG monoclonal anti-MB labeled with B5 initiator, amplifier B5-Alexa546.
- **Ch3:** Target mRNA *Prkcd*, probe set with 31 split-initiator probe pairs, amplifier B2-Alexa647.
- **Ch4:** Target mRNA *Slc17a7*, probe set with 36 split-initiator pairs, amplifier B4-Alexa750.
- **Ch5:** DAPI.

Additional studies are presented as follows:

- Figure S35 displays 4-plex images for $N = 3$ replicate FFPE mouse brain sections (cf. Figures 5CD).
- Figure S36 displays representative regions of individual channels used for measurement of signal and background for each target.
- Table S26 displays estimated values for signal, background, noise, and signal-to-background for each target.

**Protocol:** Simultaneous HCR 1°IHC + HCR RNA-ISH (Section S3.2) using initiator-labeled primary antibody probes for protein targets, split-initiator DNA probes for RNA targets, and simultaneous HCR signal amplification for all targets.
**Sample:** FFPE C57BL/6 mouse brain section (coronal); thickness: 5 $\mu$m.
**Microscopy:** Epifluorescence.

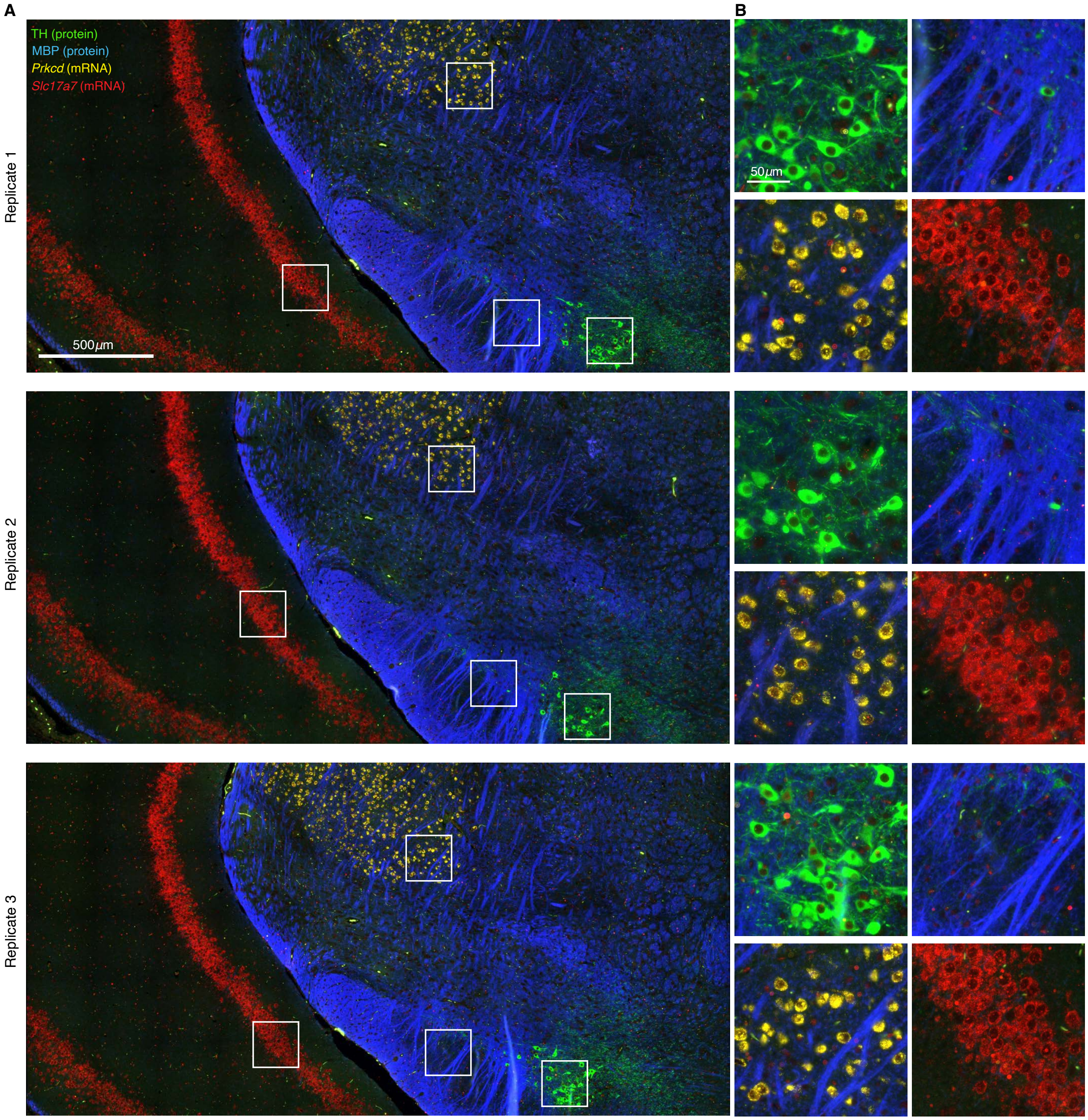

**Figure S35. Replicates for 4-plex simultaneous protein and mRNA imaging using HCR 1°IHC and HCR RNA-ISH in FFPE mouse brain sections (cf. Figures 5CD).** (A) 4-channel epifluorescence images for 3 replicate FFPE mouse brain sections. (B) Zoom of the depicted regions. Ch1: target protein TH (Alexa488). Ch2: target protein MBP (Alexa546). Ch3: target mRNA *Prkcd* (Alexa647). Ch4: target mRNA *Slc17a7* (Alexa750). Sample: FFPE C57BL/6 mouse brain section (coronal); thickness: 5 $\mu$m.

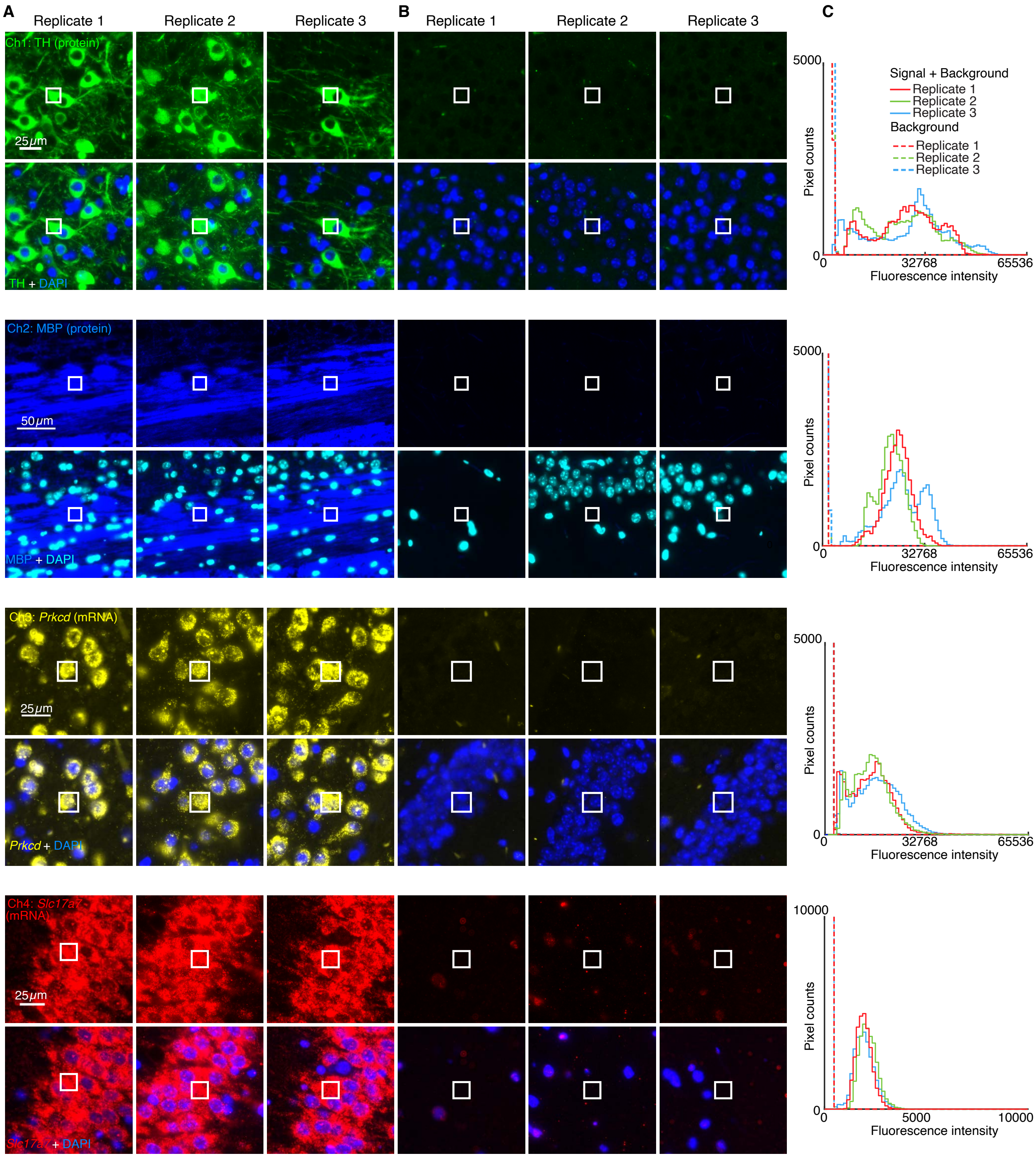

**Figure S36. Measurement of signal, background, and noise for 4-plex simultaneous protein and mRNA imaging using HCR 1°IHC and HCR RNA-ISH in FFPE mouse brain sections (cf. Figure 5CD).** Use experiment of Type 1 in Table S7A (1°Ab probe + hairpins) to measure: (A) SIG+BACK+NOISE (region of high expression), (B) BACK+NOISE (region of no/low expression) and NOISE (region with no sample; not depicted). (C) Pixel intensity histograms for representative regions (three rectangles per experiment type for each of 3 replicate mouse brain sections). Epifluorescence images collected with the microscope exposure time optimized to avoid saturating SIG+BACK pixels; DAPI channel facilitates placement of representative rectangles. Ch1: target protein TH (Alexa488). Ch2: target protein MBP (Alexa546). Ch3: target mRNA *Prkcd* (Alexa647). Ch4: target mRNA *Slc17a7* (Alexa750). Ch5: DAPI. Sample: FFPE C57BL/6 mouse brain section (coronal); thickness: 5 μm.

| Channel | Target | Type | Fluorophore | NOISE | BACK+NOISE | SIG+BACK+NOISE | BACK | SIG | SIG/BACK |
|---|---|---|---|---|---|---|---|---|---|
| Ch1 | TH | protein | Alexa488 | $425 \pm 7$ | $1900 \pm 50$ | $25\,800 \pm 900$ | $1470 \pm 50$ | $23\,900 \pm 900$ | $16.2 \pm 0.9$ |
| Ch2 | MBP | protein | Alexa546 | $500 \pm 60$ | $1160 \pm 40$ | $23\,000 \pm 2000$ | $670 \pm 70$ | $22\,000 \pm 2000$ | $33 \pm 5$ |
| Ch3 | *Prkcd* | mRNA | Alexa647 | $800 \pm 100$ | $1720 \pm 140$ | $14\,900 \pm 1100$ | $960 \pm 170$ | $13\,200 \pm 1800$ | $14 \pm 3$ |
| Ch4 | *Slc17a7* | mRNA | Alexa750 | $200 \pm 10$ | $240 \pm 10$ | $1950 \pm 120$ | $42 \pm 14$ | $1710 \pm 120$ | $41 \pm 14$ |

**Table S26. Estimated signal-to-background for 4-plex simultaneous protein and mRNA imaging using HCR 1°IHC and HCR RNA-ISH in FFPE mouse brain sections (cf. Figure 5CD).** Mean $\pm$ standard error of the mean, $N = 3$ replicate FFPE mouse brain sections. Analysis based on representative rectangular regions (examples depicted in Figure S36) using methods of Section S2.6.2.

#### S5.8 Replicates and signal-to-background measurements for simultaneous multiplexed protein and mRNA imaging using HCR 2°IHC and HCR RNA-ISH (cf. Figure 6)

##### S5.8.1 Mammalian cells on a slide

For 4-plex simultaneous protein and mRNA imaging using HCR 2°IHC + HCR RNA-ISH in mamalian cells on a slide, the 5 channels are (2 RNAs + 2 proteins + DAPI):

- **Ch1:** Target protein PCNA, probe 1°mAb mouse IgG2a anti-PCNA , probe 2°pAb goat anti-mouse IgG2a labeled with B5 initiator, amplifier B5-Alexa488.
- **Ch2:** Target protein HSP60, probe 1°mAb rabbit anti-Hsp60, probe 2°pAb donkey anti-rabbit labeled with B4 initiator, amplifier B4-Alexa546.
- **Ch3:** Target RNA *U6*, probe set with 2 split-initiator probe pairs, amplifier B1-Alexa594.
- **Ch4:** Target mRNA *HSP60*, probe set with 18 split-initiator probe pairs, amplifier B2-Alexa647.
- **Ch5:** DAPI.

Additional studies are presented as follows:

- Figure S37 displays 4-plex images for $N = 3$ replicate wells on a multi-well slide (cf. Figure 6B).
- Figure S38 displays representative regions of individual channels used for measurement of signal and background for each target.
- Table S27 displays estimated values for signal, background, and signal-to-background for each target.

**Protocol:** Simultaneous HCR 2°IHC + HCR RNA-ISH (Section S4.1) using unlabeled primary antibody probes and initiator-labeled secondary antibody probes for protein targets, split-initiator DNA probes for RNA targets, and simultaneous HCR signal amplification for all targets.
**Sample:** HeLa cells.
**Microscopy:** Confocal.

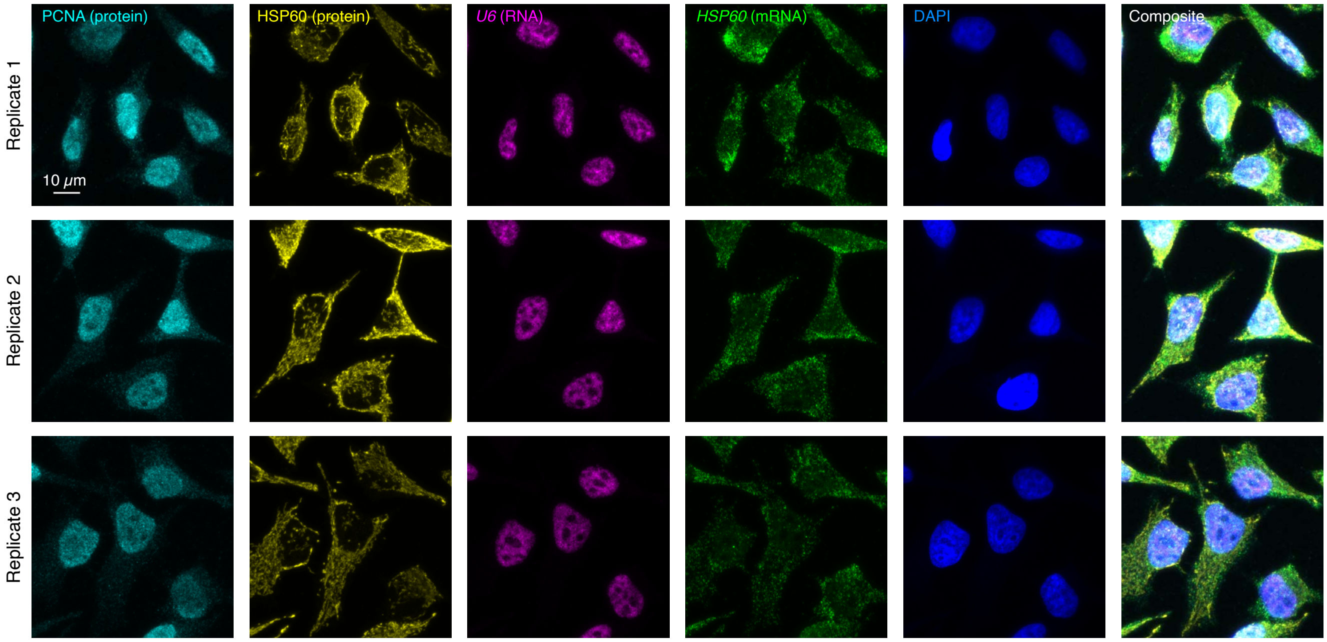

**Figure S37. Replicates for 4-plex simultaneous protein and mRNA imaging using HCR 2°IHC and HCR RNA-ISH in mammalian cells on slides (cf. Figures 6B).** 5-channel confocal images for 3 replicate wells in a multi-well slide; maximum intensity z-projection. Ch1: target protein PCNA (Alexa488). Ch2: target protein Hsp60 (Alexa546). Ch3: target RNA *U6* (Alexa594). Ch4: target mRNA *Hsp60* (Alexa647). Ch5: DAPI. Sample: HeLa cells.

| Channel | Target | Type | Fluorophore | BACK | SIG+BACK | SIG | SIG/BACK |
|---|---|---|---|---|---|---|---|
| Ch1 | PCNA | protein | Alexa488 | $453 \pm 12$ | $13\,700 \pm 1000$ | $12\,200 \pm 1000$ | $27 \pm 2$ |
| Ch2 | Hsp60 | protein | Alexa546 | $158 \pm 5$ | $38\,000 \pm 3000$ | $38\,000 \pm 3000$ | $240 \pm 20$ |
| Ch3 | *U6* | RNA | Alexa594 | $135 \pm 3$ | $38\,000 \pm 2000$ | $38\,000 \pm 2000$ | $279 \pm 17$ |
| Ch4 | *Hsp60* | mRNA | Alexa647 | $202 \pm 9$ | $9300 \pm 1100$ | $9100 \pm 1100$ | $45 \pm 6$ |

**Table S27. Estimated signal-to-background for 4-plex simultaneous protein and mRNA imaging using HCR 2°IHC and HCR RNA-ISH in mammalian cells on a slide (cf. Figure 6B).** For protein targets, the signal estimate SIG is calculated using the background approximation BACK $\approx$ $NSD_{2^\circ}$+NSA+AF. For RNA targets, the signal estimate SIG is calculated using the background approximation BACK $\approx$ NSA+AF. Instrument noise is negligible using confocal microscopy so calculations use the approximation NOISE $\approx 0$. Mean $\pm$ standard error of the mean, $N = 15$ representative rectangular regions (one rectangle in each of 5 individual cells in each of 3 replicate wells on a multi-well slide). Analysis based on representative rectangular regions (examples depicted in Figure S38) using methods of Section S2.6.2.

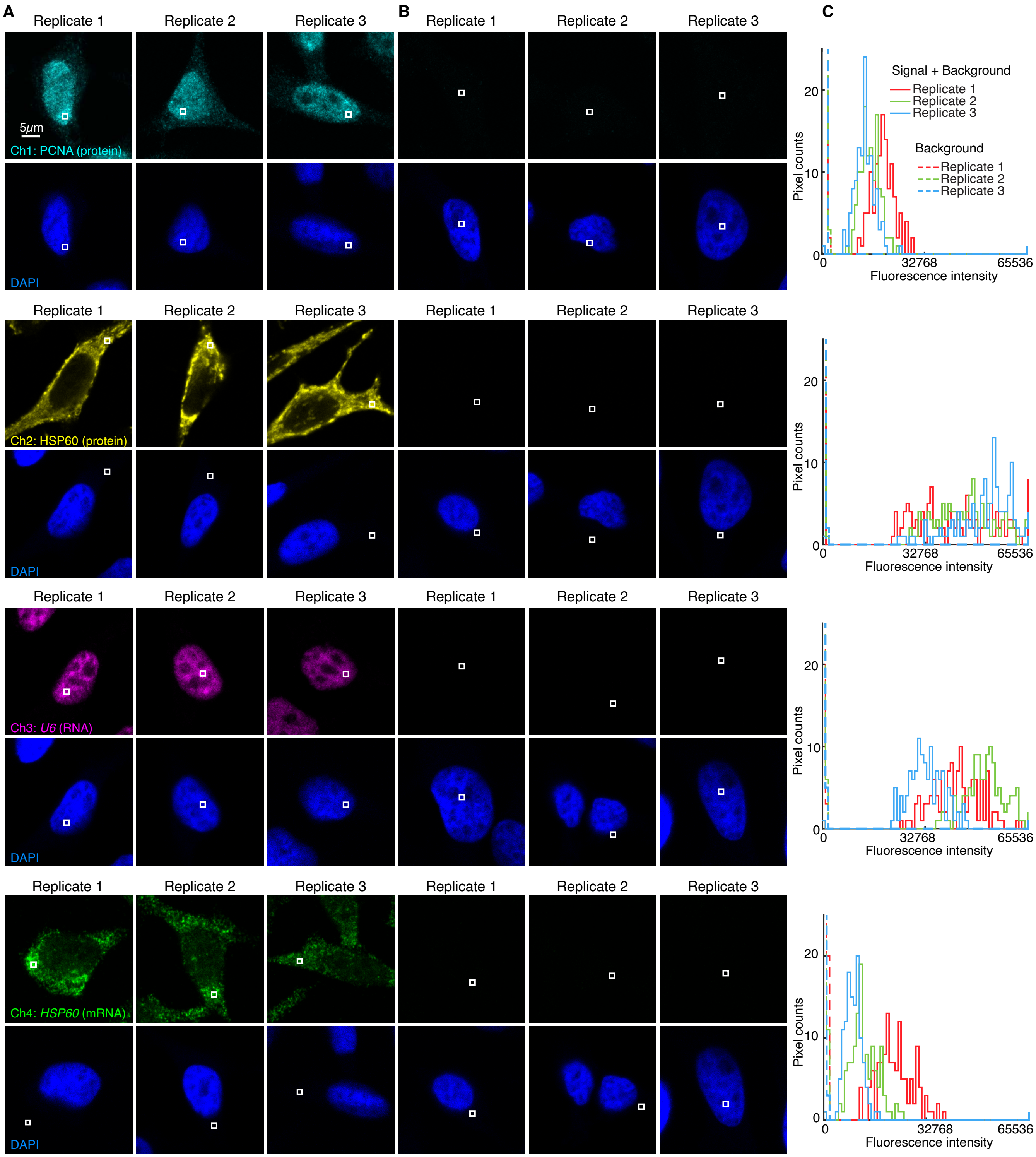

**Figure S38. Measurement of signal and background for 4-plex simultaneous protein and mRNA imaging using HCR 2°IHC and HCR RNA-ISH in mammalian cells on a slide (cf. Figure 6B).** (A) Use experiment of Type 1 in Table S8A (1°Ab probe + 2°Ab probe + hairpins) to measure SIG+BACK in a region of high expression. (B) For RNA targets, use experiment of Type 2 in Table S8B (2°Ab probe + hairpins) to measure NSA+AF in a region of maximum background. For protein targets, use experiment of Type 4 in Table S8B (2°Ab probe + hairpins) to measure $NSD_{2^\circ}$+NSA+AF in a region of maximum background. (C) Pixel intensity histograms for representative regions (one rectangle in each of 5 individual cells in each of 3 replicate wells on a multi-well slide). Confocal images collected with the microscope gain optimized to avoid saturating SIG+BACK pixels; DAPI channel facilitates placement of representative rectangles; single optical section. Ch1: target protein PCNA (Alexa488). Ch2: target protein HSP60 (Alexa546). Ch3: target RNA *U6* (Alexa594). Ch4: target mRNA *Hsp60* (Alexa647). Ch5: DAPI. Sample: HeLa cells.

##### S5.8.2 FFPE mouse brain sections

For 4-plex simultaneous protein and mRNA imaging using HCR 2°IHC + HCR RNA-ISH in FFPE mouse brain sections, the 5 channels are (2 proteins + 2 RNAs + DAPI):

- **Ch1:** Target protein TH, probe 1°pAb probe sheep IgG anti-TH, probe 2°pAb donkey anti-sheep labeled with B4 initiator, amplifier B4-Alexa488.
- **Ch2:** Target protein MBP, probe 1°mAb probe rat IgG2A anti-MBP, probe 2°pAb donkey anti-rat IgG labeled with B3 initiator, amplifier B3-Alexa546.
- **Ch3:** Target mRNA *Prkcd*, probe set with 31 split-initiator probe pairs, amplifier B1-Alexa647.
- **Ch4:** Target mRNA *Slc17a7*, probe set with 36 split-initiator probe pairs, amplifier B2-Alexa750.
- **Ch5:** DAPI.

Additional studies are presented as follows:

- Figure S39 displays 4-plex images for $N = 3$ replicate FFPE mouse brain sections (cf. Figures 6CD).
- Figure S40 displays representative regions of individual channels used for measurement of signal and background for each target.
- Table S28 displays estimated values for signal, background, noise, and signal-to-background for each target.

**Protocol:** Simultaneous HCR 2°IHC + HCR RNA-ISH (Section S4.2) using unlabeled primary antibody probes and initiator-labeled secondary antibody probes for protein targets, split-initiator DNA probes for RNA targets, and simultaneous HCR signal amplification for all targets.
**Sample:** FFPE C57BL/6 mouse brain section (coronal); thickness: 5 $\mu$m.
**Microscopy:** Epifluorescence.

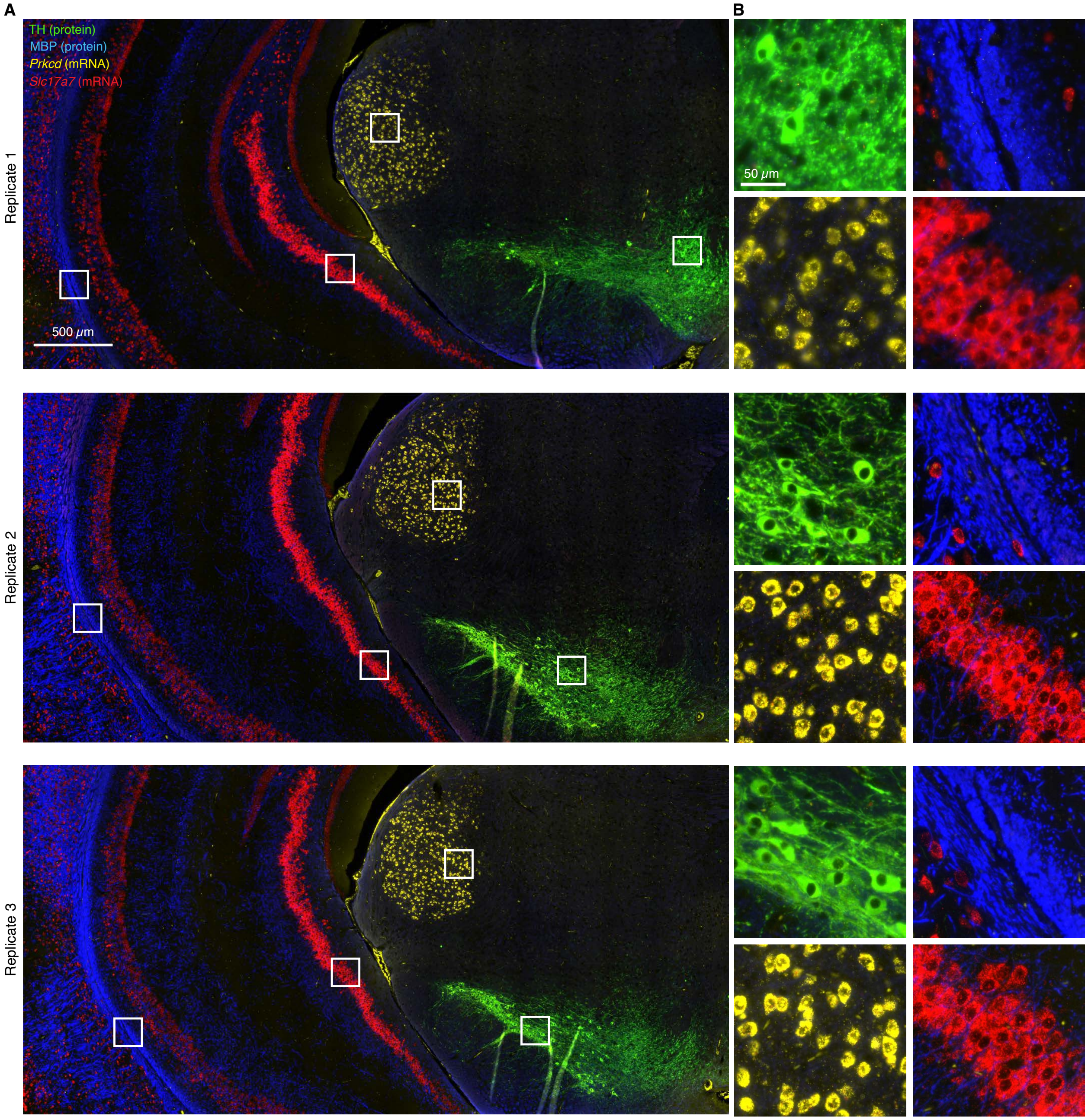

**Figure S39. Replicates for 4-plex simultaneous protein and mRNA imaging using HCR 2°IHC and HCR RNA-ISH in FFPE mouse brain sections (cf. Figures 6CD).** (A) 4-channel epifluorescence images for 3 replicate FFPE mouse brain sections. (B) Zoom of the depicted regions. Ch1: target protein TH (Alexa488). Ch2: target protein MBP (Alexa546). Ch3: target mRNA *Prkcd* (Alexa647). Ch4: target mRNA *Slc17a7* (Alexa750). Sample: FFPE C57BL/6 mouse brain section (coronal); thickness: 5 μm.

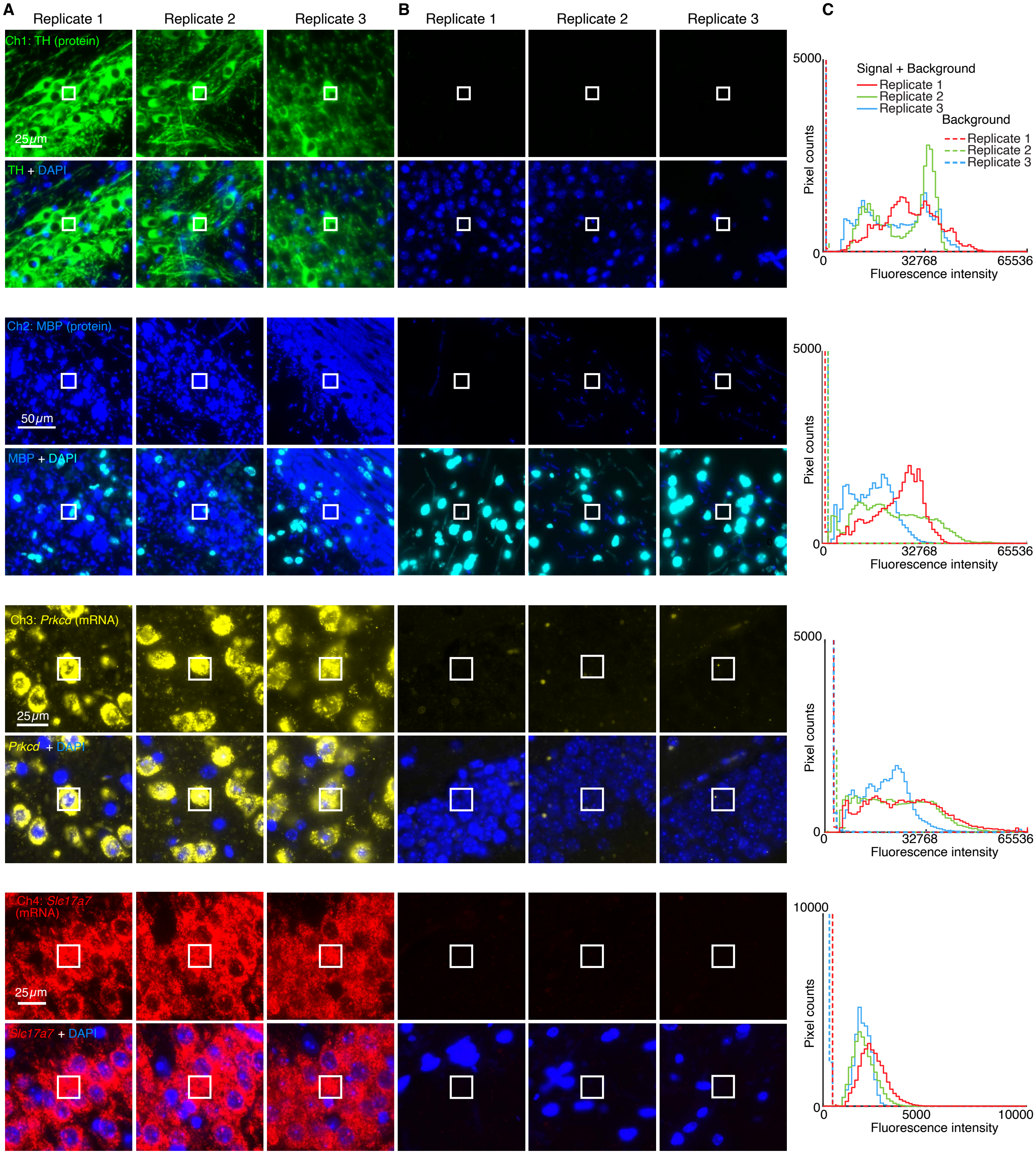

**Figure S40. Measurement of signal, background, and noise for 4-plex simultaneous protein and mRNA imaging using HCR 2°IHC and HCR RNA-ISH in FFPE mouse brain sections (cf. Figure 6CD).** Use experiment of Type 1 in Table S8A (1°Ab probe + 2°Ab probe + hairpins) to measure (A) SIG+BACK+NOISE in a region of high expression, (B) BACK+NOISE in a region of no/low expression, and NOISE in a region with no sample (not depicted). (C) Pixel intensity histograms for representative regions (three rectangles per experiment type for each of 3 replicate mouse brain sections). Epifluorescence images collected with the microscope exposure time optimized to avoid saturating SIG+BACK pixels; DAPI channel facilitates placement of representative rectangles. Ch1: target protein TH (Alexa488). Ch2: target protein MBP (Alexa546). Ch3: target mRNA *Prkcd* (Alexa647). Ch4: target mRNA *Slc17a7* (Alexa750). Ch5: DAPI. Sample: FFPE C57BL/6 mouse brain section (coronal); thickness: 5 $\mu$m.

| Channel | Target | Type | Fluorophore | NOISE | BACK+NOISE | SIG+BACK+NOISE | BACK | SIG | SIG/BACK |
|---|---|---|---|---|---|---|---|---|---|
| Ch1 | TH | protein | Alexa488 | $160 \pm 40$ | $390 \pm 30$ | $16\,000 \pm 3000$ | $234 \pm 5$ | $15\,000 \pm 3000$ | $65 \pm 17$ |
| Ch2 | MBP | protein | Alexa546 | $220 \pm 120$ | $490 \pm 40$ | $11\,000 \pm 3000$ | $270 \pm 80$ | $11\,000 \pm 3000$ | $40 \pm 16$ |
| Ch3 | *Prkcd* | mRNA | Alexa647 | $770 \pm 120$ | $1700 \pm 300$ | $23\,000 \pm 3000$ | $1000 \pm 300$ | $21\,000 \pm 3000$ | $22 \pm 8$ |
| Ch4 | *Slc17a7* | mRNA | Alexa750 | $195 \pm 8$ | $229 \pm 18$ | $2100 \pm 300$ | $30 \pm 20$ | $1800 \pm 300$ | $50 \pm 30$ |

**Table S28. Estimated signal-to-background for 4-plex simultaneous protein and mRNA imaging using HCR 2°IHC and HCR RNA-ISH in FFPE mouse brain sections (cf. Figure 6CD).** Mean ± standard error of the mean, $N = 3$ replicate mouse brain sections. Analysis based on a representative rectangular regions (examples depicted in Figure S40) using methods of Section S2.6.2.

**Krienen, F. M., Goldman, M., Zhang, Q., C. H. del Rosario, R., Florio, M., Machold, R., Saunders, A., Levandowski, K., Zaniewski, H., Schuman, B., Wu, C., Lutservitz, A., Mullally, C. D., Reed, N., Bien, E., Bortolin, L., Fernandez-Otero, M., Lin, J. D., Wysoker, A., Nemesh, J., Kulp, D., Burns, M., Tkachev, V., Smith, R., Walsh, C. A., Dimidschstein, J., Rudy, B., S. Kean, L., Berretta, S., Fishell, G., Feng, G.,

**& McCarroll, S. A.** (2020). Innovations present in the primate interneuron repertoire. *Nature*, **586**(7828), 262–269.

**Kumar, V., Krolewski, D. M., Hebda-Bauer, E. K., Parsegian, A., Martin, B., Foltz, M., Akil, H., & Watson, S. J.** (2021). Optimization and evaluation of fluorescence in situ hybridization chain reaction in cleared fresh-frozen brain tissues. *Brain Struct. Funct.*, **226**(2), 481–499.

**Lacin, H., Chen, H. M., Long, X., Singer, R. H., Lee, T., & Truman, J. W.** (2019). Neurotransmitter identity is acquired in a lineage-restricted manner in the Drosophila CNS. *eLife*, **8**, e43701.

**Li, Y., He, X., Kawaguchi, R., Zhang, Y., Wang, Q., Monavarfeshani, A., Yang, Z., Chen, B., Shi, Z., Meng, H., Zhou, S., Zhu, J., Jacobi, A., Swarup, V., Popovich, P. G., Geschwind, D. H., & He, Z.** (2020). Microglia-organized scar-free spinal cord repair in neonatal mice. *Nature*, **587**(7835), 613–618.

**Lignell, A., Kerosuo, L., Streichan, S. J., Cai, L., & Bronner, M. E.** (2017). Identification of a neural crest stem cell niche by spatial genomic analysis. *Nat. Commun.*, **8**(1), 1830.

**Ling, I. T. C., & Sauka-Spengler, T.** (2019). Early chromatin shaping predetermines multipotent vagal neural crest into neural, neuronal and mesenchymal lineages. *Nat. Cell Biol.*, **21**(12), 1504–1517.

**Liu, Y., Zou, R. S., He, S., Nihongaki, Y., Li, X., Razavi, S., Wu, B., & Ha, T.** (2020). Very fast CRISPR on demand. *Science*, **368**(6496), 1265–1269.

**Lovett-Barron, M., Chen, R., Bradbury, S., Andalman, A. S., Wagle, M., Guo, S., & Deisseroth, K.** (2020). Multiple convergent hypothalamus–brainstem circuits drive defensive behavior. *Nat. Neurosci.*, **23**(8), 959–967.

**Mantri, M., Scuderi, G. J., Abedini-Nassab, R., Wang, M. F. Z., McKellar, D., Shi, H., Grodner, B., Butcher, J. T., & De Vlaminck, I.** (2021). Spatiotemporal single-cell RNA sequencing of developing chicken hearts identifies interplay between cellular differentiation and morphogenesis. *Nat. Commun.*, **12**(1), 1771.

**Marconi, A., Hancock-Ronemus, A., & Gillis, J. A.** (2020). Adult chondrogenesis and spontaneous cartilage repair in the skate, Leucoraja erinacea. *eLife*, **9**(May), e53414.

**May-Zhang, A. A., Tycksen, E., Southard-Smith, A. N., Deal, K. K., Benthal, J. T., Buehler, D. P., Adam, M., Simmons, A. J., Monaghan, J. R., Matlock, B. K., Flaherty, D. K., Potter, S. S., Lau, K. S., & Southard-Smith, E. M.** (2021). Combinatorial transcriptional profiling of mouse and human enteric neurons identifies shared and disparate subtypes in situ. *Gastroenterology*, **160**(3), 755–770.e26.

**McLennan, R., Schumacher, L. J., Morrison, J. A., Teddy, J. M., Ridenour, D. A., Box, A. C., Semerad, C. L., Li, H., McDowell, W., Kay, D., Maini, P. K., Baker, R. E., & Kulesa, P. M.** (2015). Neural crest migration is driven by a few trailblazer cells with a unique molecular signature narrowly confined to the invasive front. *Development*, **142**(11), 2014–2025.

**Meinecke, L., Sharma, P. P., Du, H., Zhang, L., Nie, Q., & Schilling, T. F.** (2018). Modeling craniofacial development reveals spatiotemporal constraints on robust patterning of the mandibular arch. *PLOS Comput. Biol.*, **14**(11), e1006569.

**Metcalfe, K. S., Murali, R., Mullin, S. W., Connon, S. A., & Orphan, V. J.** (2021). Experimentally-validated correlation analysis reveals new anaerobic methane oxidation partnerships with consortium-level heterogeneity in diazotrophy. *ISME J.*, **15**(2), 377–396.

**Mich, J. K., Graybuck, L. T., Hess, E. E., Mahoney, J. T., Kojima, Y., Ding, Yi, Somasundaram, S., Miller, J. A., Kalmbach, B. E., Radaelli, C., Gore, B. B., Weed, N., Omstead, V., Bishaw, Y., Shapovalova, N. V., Martinez, R. A., Fong, O., Yao, S., Mortrud, M., Chong, P., Loftus, L., Bertagnolli, D., Goldy, J., Casper, T., Dee, N., Opitz-Araya, X., Cetin, A., Smith, K. A., Gwinn, R. P., Cobbs, C., Ko, A. L., Ojemann, J. G., Keene, C. D., Silbergeld, D. L., Sunkin, S. M., Gradinaru, V., Horwitz, G. D., Zeng, H., Tasic, B., Lein, E. S., Ting, J. T., & Levi, B. P.** (2021). Functional enhancer elements drive subclass-selective expression from mouse to primate neocortex. *Cell Rep.*, **34**(13), 108754.

**Michael, V., Goffinet, J., Pearson, J., Wang, F., Tschida, K., & Mooney, R.** (2020). Circuit and synaptic organization of forebrain-to-midbrain pathways that promote and suppress vocalization. *eLife*, **9**, e63493.

**Michki, N. S., Li, Y., Sanjasaz, K., Zhao, Y., Shen, F. Y., Walker, L. A., Cao, W., Lee, C.-Y., & Cai, D.** (2021). The molecular landscape of neural differentiation in the developing *Drosophila* brain revealed by targeted scRNA-seq and multi-informatic analysis. *Cell Rep.*, **35**(4), 109039.

**Milewska, A., Kula-Pacurar, A., Wadas, J., Suder, A., Szczepanski, A., Dabrowska, A., Owczarek, K., Marcello, A., Ochman, M., Stacel, T., Rajfur, Z., Sanak, M., Labaj, P., Branicki, W., & Pyrc, K.** (2020). Replication of severe acute respiratory syndrome coronavirus 2 in human respiratory epithelium. *J. Virol.*, **94**(15), e00957–20.

**Moriano-Gutierrez, S., Bongrand, C., Essock-Burns, T., Wu, L., McFall-Ngai, Margaret J., & Ruby, E. G.** (2020). The noncoding small RNA SsrA is released by *Vibrio fischeri* and modulates critical host responses. *PLOS Biol.*, **18**(11), e3000934.

**Mu, W., Li, S., Xu, J., Guo, X., Wu, H., Chen, Z., Qiao, L., Helfer, G., Lu, F., Liu, C., & Wu, Q. F.** (2021). Hypothalamic Rax + tanycytes contribute to tissue repair and tumorigenesis upon oncogene activation in mice. *Nat. Commun.*, **12**(1), 2288.

**Nandagopal, N., Santat, L. A., & Elowitz, M. B.** (2019). Cis-activation in the Notch signaling pathway. *eLife*, **8**, e37880.

**Nikolakakis, K., Lehnert, E., McFall-Ngai, M. J., & Ruby, E. G.** (2015). Use of hybridization chain reaction-fluorescent in situ hybridization to track gene expression by both partners during initiation of symbiosis. *Appl. Environ. Microbiol.*, **81**(14), 4728–4735.

**O'Brown, N. M., Megason, S. G., & Gu, C.** (2019). Suppression of transcytosis regulates zebrafish blood-brain barrier function. *eLife*, **8**, e47326.

**Park, Y.-G., Sohn, C. H., Chen, R., McCue, M., Yun, D. H., Drummond, G. T., Ku, T., Evans, N. B., Oak, H. C., Trieu, W., Choi, H., Jin, X., Lilascharoen, V., Wang, J., Truttmann, M. C., Qi, H. W., Ploegh, H. L., Golub, T. R., Chen, S.-C., Frosch, M. P., Kulik, H. J., Lim, B. K., & Chung, K.** (2018). Protection of tissue physicochemical properties using polyfunctional crosslinkers. *Nat. Biotechnol.*, **37**(1), 73–83.

**Patriarchi, T., Cho, J. R., Merten, K., Howe, M. W., Marley, A., Xiong, W. H., Folk, R. W., Broussard, G. J., Liang, R., Jang, M. J., Zhong, H., Dombeck, D., Zastrow, M. V., Nimmerjahn, A., Gradinaru, V., Williams, J. T., & Tian, L.** (2018). Ultrafast neuronal imaging of dopamine dynamics with designed genetically encoded sensors. *Science*, **360**(6396).

**Pond, A. J. R., Hwang, S., Verd, B., & Steventon, B.** (2021). A deep learning approach for staging embryonic tissue isolates with small data. *PLOS ONE*, **16**(1), e0244151.

**Ren, J., Isakova, A., Friedmann, D., Zeng, J., Grutzner, S. M., Pun, A., Zhao, G. Q., Kolluru, S. S., Wang, R., Lin, R., Li, P., Li, A., Raymond, J. L., Luo, Q., Luo, M., Quake, S. R., & Luo, L.** (2019). Single-cell transcriptomes and whole-brain projections of serotonin neurons in the mouse dorsal and median raphe nuclei. *eLife*, **8**, e49424.

**Rodriguez, C. M., Wright, S. E., Kearse, M. G., Haenfler, J. M., Flores, B. N., Liu, Y., Ifrim, M. F., Glineburg, M. R., Krans, A., Jafar-Nejad, P., Sutton, M. A., Bassell, G. J., Parent, J. M., Rigo, F., Barmada, S. J., & Todd, P. K.** (2020). A native function for RAN translation and CGG repeats in regulating fragile X protein synthesis. *Nat. Neurosci.*, **23**(3), 386–397.

**Rosenthal, A. Z., Zhang, X. N., Lucey, K. S., Ottesen, E. A., Trivedi, V., Choi, H. M. T., Pierce, N. A., & Leadbetter, J. R.** (2013). Localizing transcripts to single cells suggests an important role of uncultured deltaproteobacteria in the termite gut hydrogen economy. *Proc. Natl. Acad. Sci. U. S. A.*, **110**(40), 16163–16168.

**Schloissnig, S., Kawaguchi, A., Nowoshilow, S., Falcon, F., Otsuki, L., Tardivo, P., Timoshevskaya, N., Keinath, M. C., Smith, J. J., Voss, S. R., & Tanaka, E. M.** (2021). The giant axolotl genome uncovers the evolution, scaling, and transcriptional control of complex gene loci. *Proc. Natl. Acad. Sci.*, **118**(15).

**Shah, S., Lubeck, E., Zhou, W., & Cai, L.** (2016a). In situ transcription profiling of single cells reveals spatial organization of cells in the mouse hippocampus. *Neuron*, **92**, 342–357.

**Shah, S., Lubeck, E., Schwarzkopf, M., He, T.-F., Greenbaum, A., Sohn, C. H., Lignell, A., Choi, H. M. T., Gradinaru, V., Pierce, N. A., & Cai, L.** (2016b). Single-molecule RNA detection at depth via hybridization chain reaction and tissue hydrogel embedding and clearing. *Development*, **143**, 2862–2867.

**Simões, F. C., Cahill, T. J., Kenyon, A., Gavriouchkina, D., Vieira, J. M., Sun, X., Pezzolla, D., Ravaud, C., Masmanian, E., Weinberger, M., Mayes, S., Lemieux, M. E., Barnette, D. N., Gunadasa-Rohling, M., Williams, R. M., Greaves, D. R., Trinh, L. A., Fraser, S. E., Dallas, S. L., Choudhury, R. P., Sauka-Spengler, T., & Riley, P. R.** (2020). Macrophages directly contribute collagen to scar formation during zebrafish heart regeneration and mouse heart repair. *Nat. Commun.*, **11**(1), 600.

**Sui, Q., Zhu, J., Li, X., Knight, G. E., He, C., Burnstock, G., Yuan, H., & Xiang, Z.** (2016). A modified protocol for the detection of three different mRNAs with a new-generation in situ hybridization chain reaction on frozen sections. *J. Mol. Histol.*, **47**(6), 511–529.

**Sylwestrak, E. L., Rajasethupathy, P., Wright, M. A., Jaffe, A., & Deisseroth, K.** (2016). Multiplexed intact-tissue transcriptional analysis at cellular resolution. *Cell*, **164**(4), 792–804.

**Ton, Q. V., Leino, D., Mowery, S. A., Bredemeier, N. O., Lafontant, P. J., Lubert, A., Gurung, S., Farlow, J. L., Foroud, T. M., Broderick, J., & Sumanas, S.** (2018). Collagen COL22A1 maintains vascular stability and mutations in COL22A1 are potentially associated with intracranial aneurysms. *Dis. Model. Mech.*, **11**(dmm033654).

**Trivedi, V., Choi, H. M. T., Fraser, S. E., & Pierce, N. A.** (2018). Multidimensional quantitative analysis of mRNA expression within intact vertebrate embryos. *Development*, **145**, dev156869.

**Tsai, T. Y.-C., Sikora, M., Xia, P., Colak-Champollion, T., Knaut, H., Heisenberg, C., & Megason, S. G.** (2020). An adhesion code ensures robust pattern formation during tissue morphogenesis. *Science*, **370**(6512), 113–116.

**Tu, R., Duan, B., Song, X., Chen, S., Scott, A., Hall, K., Blanck, J., DeGraffenreid, D., Li, H., Perera, A., Haug, J., & Xie, T.** (2021). Multiple niche compartments orchestrate stepwise germline stem cell progeny differentiation. *Curr. Biol.*, **31**(4), 827–839.e3.

**van den Brink, S. C., Alemany, A., van Batenburg, V., Moris, N., Blotenburg, M., Vivié, J., Baillie-Johnson, P., Nichols, J., Sonnen, K. F., Martinez Arias, A., & van Oudenaarden, A.** (2020). Single-cell and spatial transcriptomics reveal somitogenesis in gastruloids. *Nature*, **582**(7812), 405–409.

**von Buchholtz, L. J., Lam, R. M., Emrick, J. J., Chesler, A. T., & Ryba, N. J. P.** (2020). Assigning transcriptomic class in the trigeminal ganglion using multiplex in situ hybridization and machine learning. *Pain*, **161**(9), 2212–2224.

**Wang, H., Holland, P. W. H., & Takahashi, T.** (2019). Gene profiling of head mesoderm in early zebrafish development: Insights into the evolution of cranial mesoderm. *EvoDevo*, **10**(1), 14.

**Weinberger, M, Simões, F. C., Patient, R., Sauka-Spengler, T., & Riley, P. R.** (2020). Functional heterogeneity within the developing zebrafish epicardium. *Dev. Cell*, **52**(5), 574–590.e6.

**Wells, A. I., Grimes, K. A., Kim, K., Branche, E., Bakkenist, C. J., DePas, W. H., Shresta, S., & Coyne, C. B.** (2021). Human FcRn expression and Type I Interferon signaling control Echovirus 11 pathogenesis in mice. *PLOS Pathog.*, **17**(1), e1009252.

**Williams, R. M., Candido-Ferreira, I., Repapi, E., Gavriouchkina, D., Senanayake, U., Ling, I. T. C., Telenius, J., Taylor, S., Hughes, J., & Sauka-Spengler, T.** (2019). Reconstruction of the global neural crest gene regulatory network in vivo. *Dev. Cell*, **51**(2), 255–276.e7.

**Wurster, S., Ruiz, O. E., Samms, K. M., Tatara, A. M., Albert, N. D., Kahan, P. H., Nguyen, A. T., Mikos, A. G., Kontoyiannis, D. P., & Eisenhoffer, G. T.** (2021). EGF-mediated suppression of cell extrusion during mucosal damage attenuates opportunistic fungal invasion. *Cell Rep.*, **34**(12), 108896.

**Yamaguchi, T., Kawakami, S., Hatamoto, M., Imachi, H., Takahashi, M., Araki, N., Yamaguchi, T., & Kubota, K.** (2015). In situ DNA-hybridization chain reaction (HCR): A facilitated in situ HCR system for the detection of environmental microorganisms. *Environ. Microbiol.*, **17**(7), 2532–2541.

**Young, W. S., & Song, J.** (2020). Characterization of oxytocin receptor expression within various neuronal populations of the mouse dorsal hippocampus. *Front. Mol. Neurosci.*, **13**.

**Zhuang, P., Zhang, H., Welchko, R. M., Thompson, R. C., Xu, S., & Turner, D. L.** (2020). Combined microRNA and mRNA detection in mammalian retinas by in situ hybridization chain reaction. *Sci. Rep.*, **10**(1), 351.
